# Post-translational regulation of photosynthetic activity via the TOR kinase in plants

**DOI:** 10.1101/2023.05.05.539554

**Authors:** Stefano D’Alessandro, Florent Velay, Régine Lebrun, Marwa Mehrez, Shanna Romand, Rim Saadouni, Céline Forzani, Sylvie Citerne, Marie-Hélène Montané, Christophe Robaglia, Benoît Menand, Christian Meyer, Ben Field

## Abstract

Chloroplasts are the powerhouse of the plant cell, yet they are resource-intensive and will cause photooxidative damage if their activity overshoots the demands of growth. The adjustment of chloroplast activity to match growth is therefore vital for stress acclimation. Here we identify a novel post-translational mechanism linking the conserved eukaryotic TOR kinase that promotes growth and the guanosine tetraphosphate (ppGpp) signaling pathway of prokaryotic origin that regulates chloroplast activity, and photosynthesis in particular. We show that RelA SpoT Homologue 3 (RSH3), a nuclear-encoded chloroplastic enzyme responsible for ppGpp biosynthesis, interacts directly with the TOR complex via a plant-specific N-terminal region (NTR) which is hyper-phosphorylated in a TOR-dependent manner. Downregulation of TOR activity reduces NTR phosphorylation, enhances ppGpp synthesis by RSH3, and causes a ppGpp-dependent decrease in photosynthetic capacity. Altogether we demonstrate that the TOR-RSH3 signaling axis is a novel and direct post-translational mechanism that allows chloroplast activity to be matched with plant growth, setting a new precedent for the regulation of organellar function by TOR.

**One sentence summary:** The TOR kinase post-translationally controls guanosine tetraphosphate signaling to regulate plant photosynthetic activity.

## Main text

The use of sunlight to fix carbon and produce chemical energy during photosynthesis is the basis of almost all life on the planet. However, the photosynthetic machinery is also resource intensive and chloroplasts must be tightly regulated to prevent photooxidative stress. Guanosine tetraphosphate (ppGpp) is a signaling nucleotide that regulates growth and stress acclimation in the majority of prokaryotes (Bange et al., 2021). In plants, ppGpp signaling negatively regulates photosynthesis, and is required for normal growth and stress acclimation (Maekawa et al., 2015; Sugliani et al., 2016; Mehrez et al., 2022). TARGET OF RAPAMYCIN (TOR) is a nucleocytosolic Ser/Thr kinase that plays an evolutionary conserved role in eukaryotes by promoting growth in response to favorable environmental cues (Burkart and Brandizzi, 2021; Pacheco et al., 2021). Nutrient limitation or environmental stress lead to the inactivation of TOR, which slows growth and promotes nutrient recycling. TOR also influences photosynthesis in plants and algae, a phenomenon that up to now was only explained by transcriptional regulation of nuclear-encoded chloroplast genes (Dong et al., 2015; Sun et al., 2016; Imamura et al., 2018; Upadhyaya and Rao, 2019; D’Alessandro, 2022). Here we set out to determine whether TOR is involved in the post-transcriptional regulation of photosynthesis.

Using the TOR complex subunit SEC13 protein 8 (LST8) as bait in an untargeted yeast two hybrid (Y2H) screen, we identified the bifunctional ppGpp synthase / hydrolase enzymes RSH2 and RSH3 as LST8 interactors (Fig. 1A). LST8 is nucleocytosolic (Fig. S1), while RSH3 is a nuclear-encoded chloroplast enzyme expected to reside only transiently in the cytosol before chloroplast import and processing. Therefore, we adopted a proximity labelling approach to determine whether LST8 interacts with the RSH3 precursor protein *in planta*. LST8 fused to the promiscuous biotin ligase TurboID (Zhang et al., 2019) with a triple haemagglutinin (HA) tag (TID-LST8, LST8-TID) was co-expressed with RSH3-GFP or CFP targeted to the chloroplast by the Rubisco Small Subunit 5A Chloroplast Transit Peptide (CTP) (SSU-CFP). TID-LST8 and LST8-TID biotinylated proteins at the molecular weight of RSH3-GFP only in the samples co-expressing RSH3-GFP and not in the SSU-CFP control (Fig. 1B). SSU-CFP was not biotinylated (Fig. 1B, grey arrows), indicating that RSH3-GFP biotinylation is specific. We further confirmed that biotinylation is specific by purifying biotinylated proteins and showing that RSH3-GFP is biotinylated preferentially by TID-LST8 compared to a TID-YFP control (Fig. 1C).

**Figure 1.**
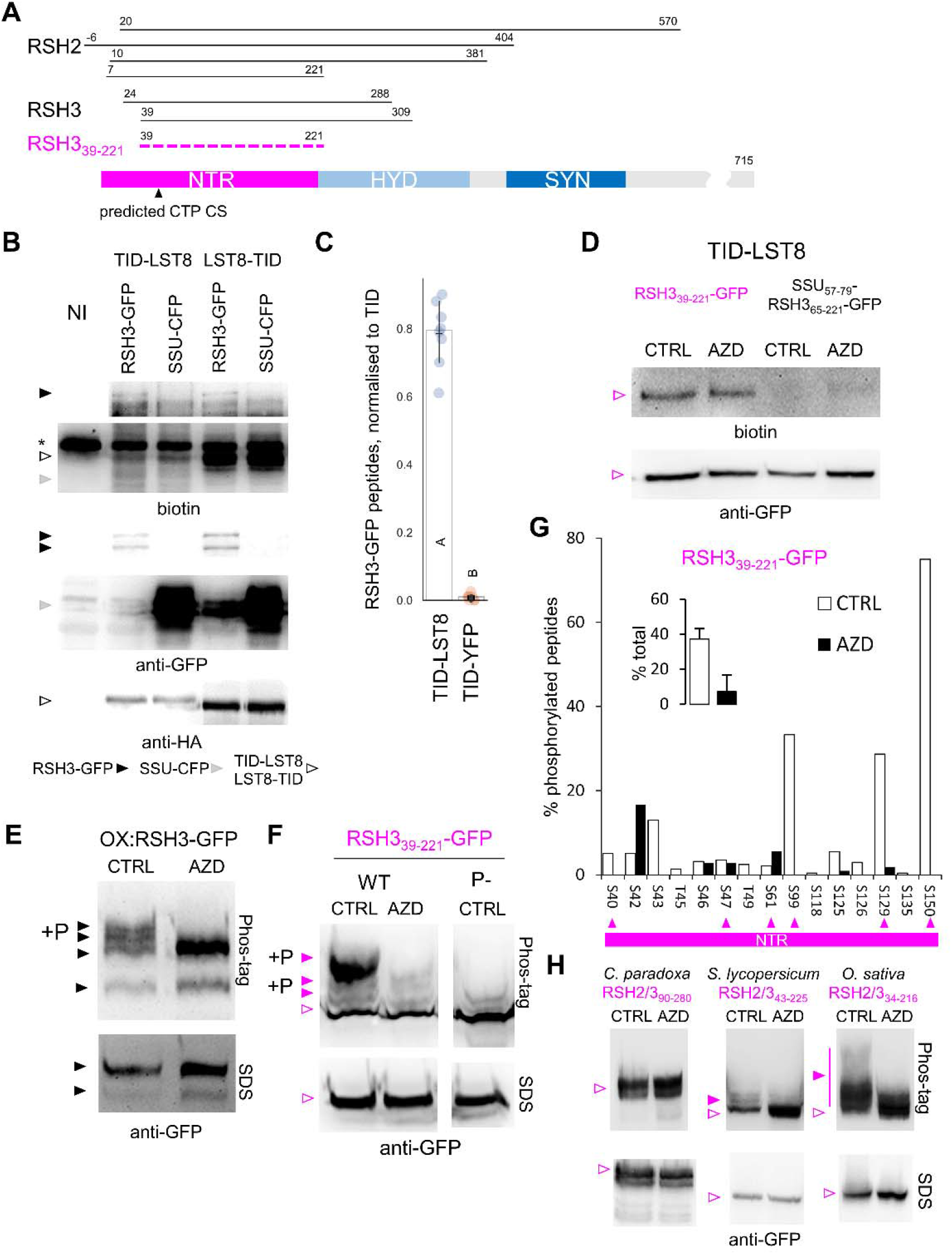
RSH enzymes interact with a subunit of the TOR complex and undergo TOR-dependent phosphorylation. (A) Alignment of the Y2H LST8 interacting regions of RSH2/3, showing the minimal interaction zone (magenta dashed line). TargetP predicted CTP cleavage site (CS) shown (Almagro Armenteros et al., 2019). (B) Blots of protein extracts from *N. benthamiana* co-expressing TID LST8 fusions and RSH3-GFP or SSU-CFP. (C) Liquid chromatography mass spectrometry (LC-MS) identification of RSH3-GFP peptides in the biotinylated protein fraction from *N. benthamiana* co-expressing RSH3-GFP with TID-LST8 or TID-YFP. Blots of protein extracts from (D) *N. benthamiana* co-expressing TID-LST8 and RSH3_39-221_-GFP or SSU_57-79_-RSH3_65-221_-GFP, (E) *A. thaliana* OXRSH3-GFP seedlings and (F) *N. benthamiana* expressing RSH3_39-221_-GFP or a phosphodefective (P-) form and treated ±AZD. (G) Map of RSH3_39-221_-GFP phosphorylation sites identified by LC-MS following immunoprecipitation from *N. benthamiana*. Arrows indicate phosphorylation in putative TOR-dependent contexts (see also Fig. S5). (H) Blots of protein extracts from *N. benthamiana* expressing RSH3_39-221_ homologous regions from *C. paradoxa, S. lycopersicum* and *O. sativa* ±AZD. NTR, N-terminal region; SYN, synthetase domain ; HYD, hydrolase domain; SDS, SDS-PAGE separation; Phos-tag, Phos-tag SDS-PAGE separation.

We next sought to determine whether RSH3_39-221,_ the minimal RSH2/3 LST8 interaction zone identified by Y2H, was sufficient for interaction with LST8 *in planta*. RSH3_39-221_-GFP was strongly biotinylated by TID-LST8 (Fig. 1D). A modified control protein SSU_59-75_-RSH3_65-221_-GFP, where the region corresponding to the predicted CTP was substituted with an equivalent region of the SSU CTP (Fig. S2A), was not biotinylated despite accumulating to the same level in the cytosol. Biotinylation was also not affected by inhibition of TOR with AZD-8055 (AZD), an ATP competitive inhibitor specific for TOR (Montané and Menand, 2013). The RSH3_39-221_ region is therefore sufficient for interaction with LST8 *in planta*, and residues 39-64 are required.

We observed that RSH3-GFP accumulates to very low levels *in planta* (Fig. 1B, S3B), shows dual bands by immunoblot (Fig. 1B, 1E, S3B), and only has sporadic chloroplast localization (Fig S3A). To determine whether the RSH3 N-terminal region (NTR) regulates RSH3 localization and stability we substituted the predicted CTP with the SSU CTP while either preserving the remaining NTR (SSU-RSH3_65-END_-GFP) or eliminating the majority of the NTR (SSU-RSH3*_195-END_-GFP) (Figure S2B). Replacement of the predicted CTP alone did not cause a major change in protein localization (Fig. S3B). However, elimination of the NTR resulted in strong accumulation of the mature form of RSH3 in the chloroplast (Fig. S3A,B). Therefore, the full NTR is involved in controlling RSH3 accumulation and localization.

We next found that the RSH3 CTP is much longer than predicted. We observed that the probable mature forms of RSH3 were of a similar size whether the NTR was present or not (Fig. S3B). This suggested the presence of additional downstream processing sites. Analysis of a series of N-terminal GFP fusions (Figure S2C) showed that RSH3_1-165_-GFP was the first to show clear chloroplast localization and processing (Fig. S4A, B). Indeed, the mature form was similar in size to GFP, suggesting that the chloroplast cleavage site was close to position 165 (Fig. S4A). Accumulation of a larger mature protein demonstrated that this cleavage site was also retained in RSH3_1-175_-GFP. These observations are consistent with the increased probability of cleavage between 160-161 (Fig. S4C). Interestingly, the dual chloroplast and nucleocytoplasmic localization of RSH3_1-175-_GFP strongly resembled that of full length RSH3, and there was reduced accumulation compared to RSH3_1-165_-GFP. Motifs important for destabilization may therefore lie between RSH3 positions 165 and 175. In conclusion, the RSH3 NTR contains a remarkably long CTP, more than double the average (Bienvenut et al., 2012), and is responsible for destabilizing RSH3 and conferring a dual nucleocytosolic and chloroplastic localization.

**Fig. S1.**
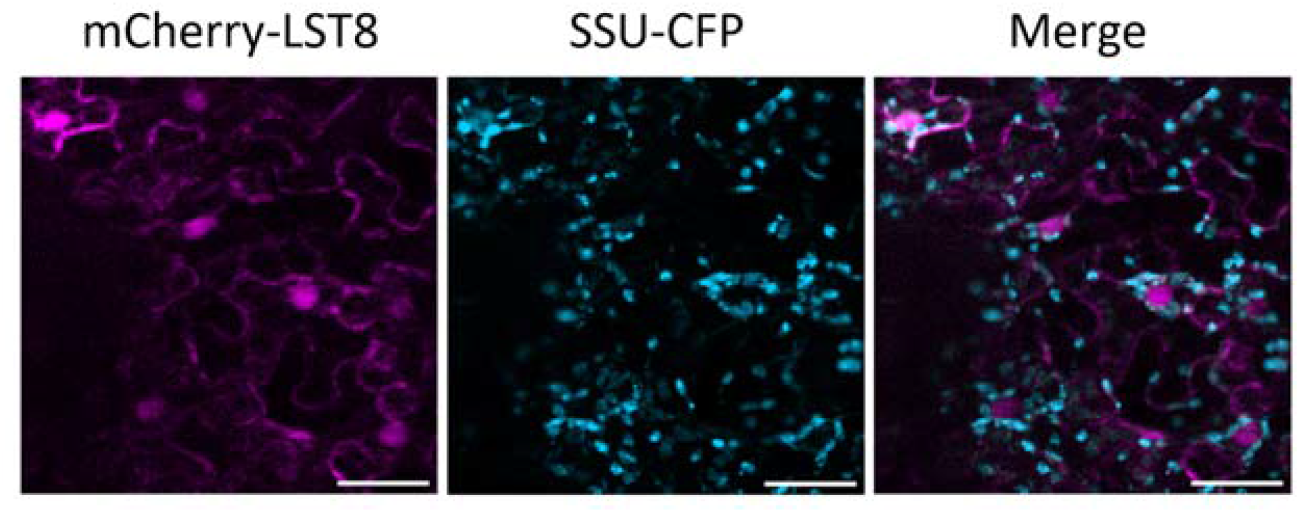
LST8 shows a nucleocytosolic localisation. Fluorescence microscopy images of *N. benthamiana* leaves expressing mCherry-LST8 and the chloroplast marker SSU-CFP. Scale bar, 50 μm.

**Fig. S2.**
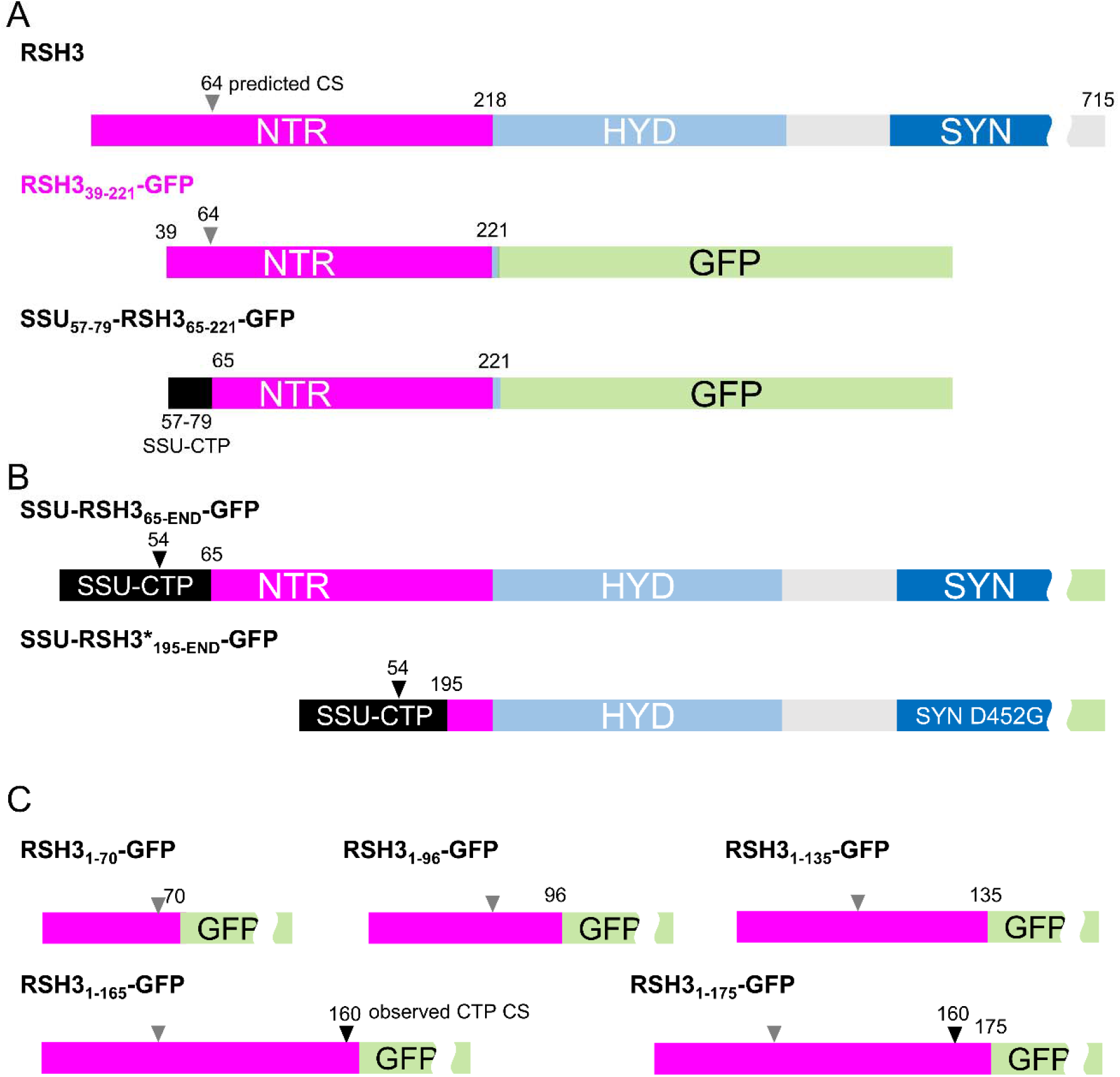
Outline of different RSH3 fusion proteins. (A) RSH3 fusions used for analyzing RSH3 control by TOR in *N. benthamiana*. (B) RSH3 with long or truncated N-terminal region targeted to chloroplast using the Rubisco small subunit CTP. Asterisk (*) indicates an inactivating mutation in the RSH3 synthetase domain (SYN D452G). (C) Series of RSH3 NTR truncations fused to GFP for determining the minimal sequence required for chloroplast targeting. Chloroplast targeting peptide (CTP), N’-terminal region (NTR, magenta), ppGpp hydrolase domain (HYD, light blue), synthetase domain (SYN, blue). TargetP predicted chloroplast import cleavage site (CS) indicated by grey arrows, and observed cleavage sites indicated by black arrows.

**Fig. S3.**
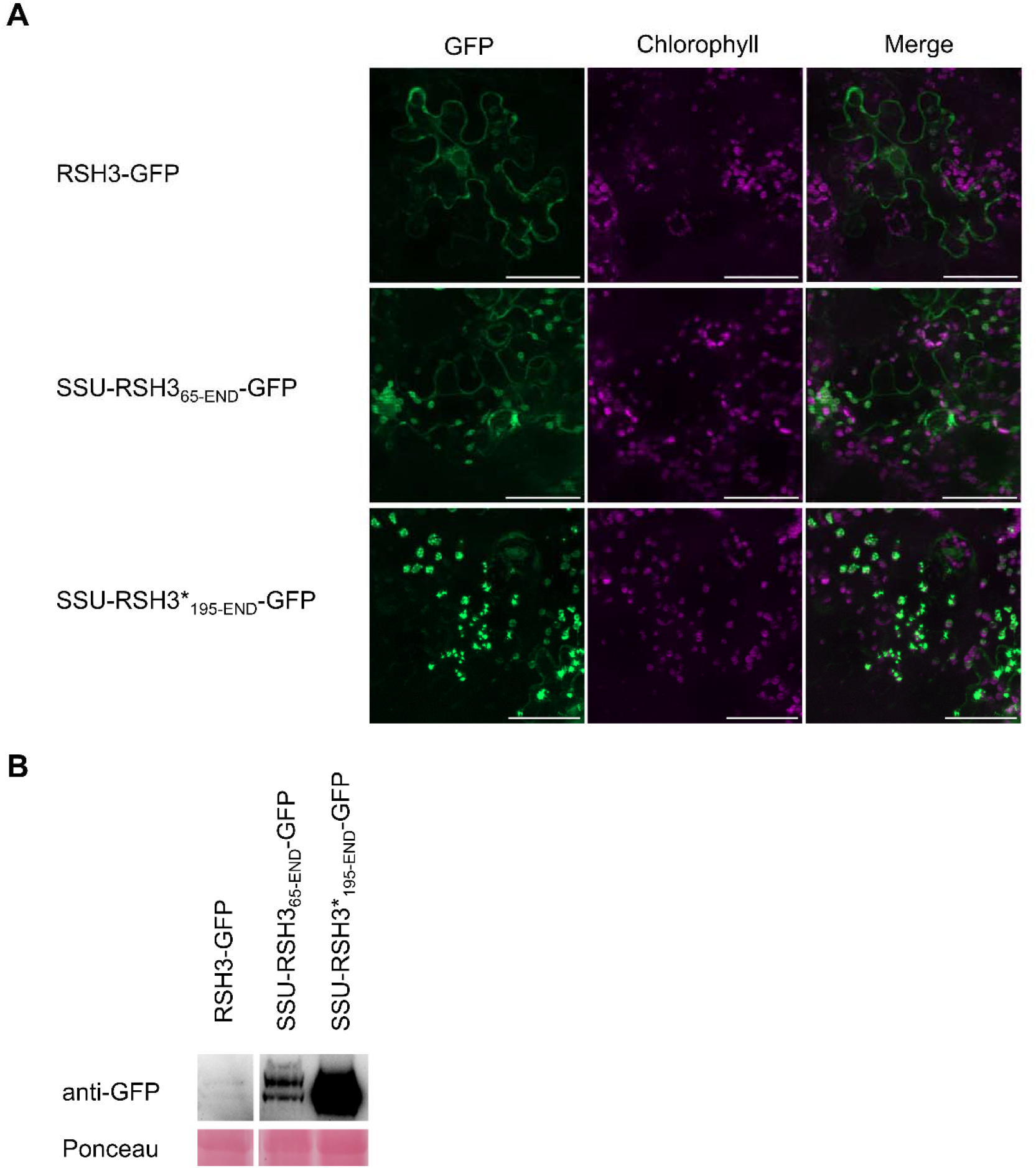
The RSH3 NTR restricts RSH3 accumulation. (A) Fluorescence microscopy images of *N. benthamiana* leaves expressing RSH3-GFP, SSU-RSH3_65-END_-GFP and SSU-RSH3*_195-END_-GFP. Scale bar, 50 μm. (B) Immunoblots of protein extracts from *N. benthamiana* expressing the same proteins.

**Fig. S4.**
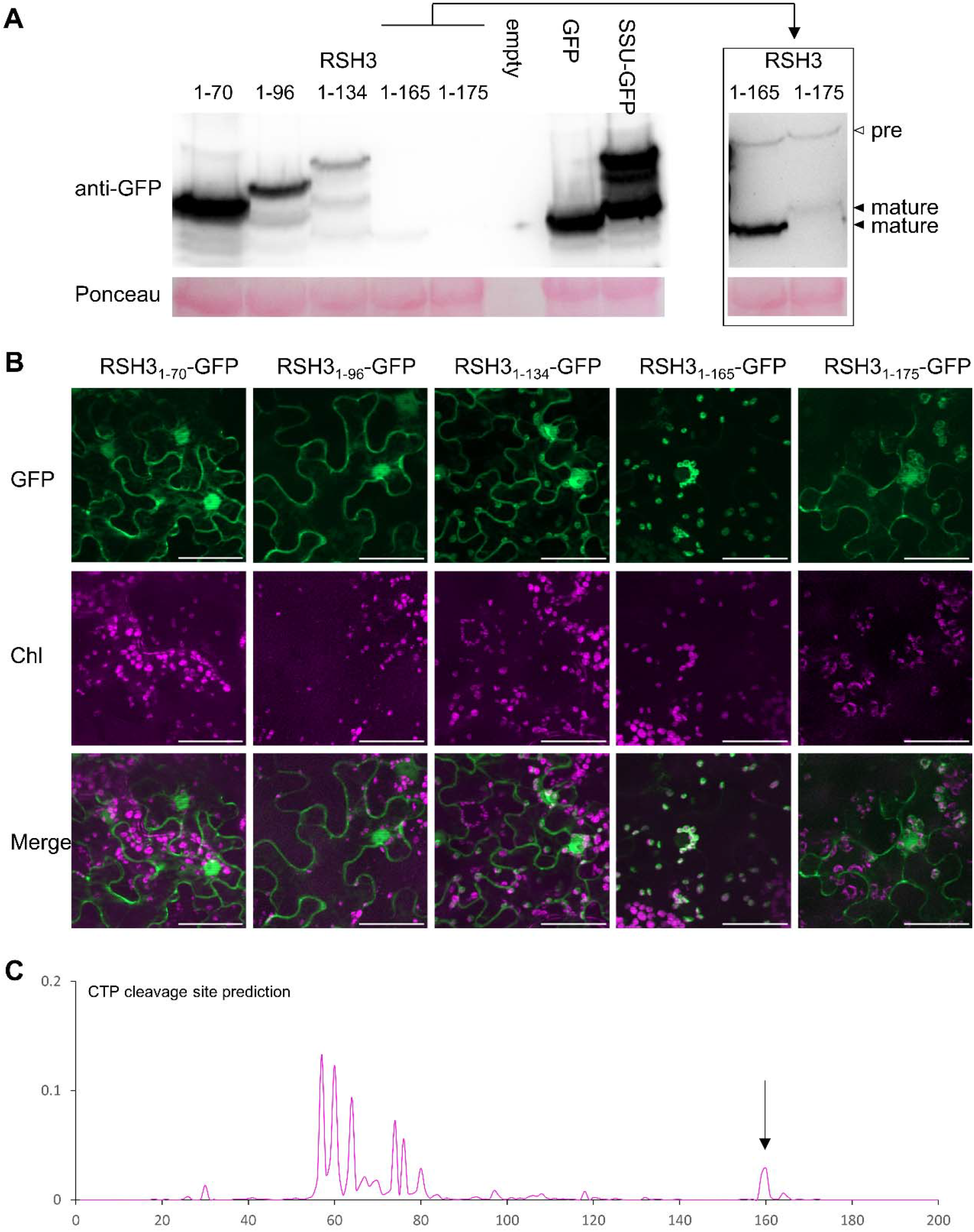
The RSH3 NTR contains an unusually long chloroplast transit peptide. A series of RSH3 N-terminal regions of increasing size were expressed in *N. benthamiana* and analyzed by (A) immunoblotting protein extracts and (B) fluorescence microscopy. Scale bar, 50 μm. (C) TargetP (Armenteros et al., 2019) prediction of RSH3 chloroplast cleavage sites (bottom).

We next tested whether the TOR complex is involved in phosphorylation of RSH3. We observed phosphoforms of the RSH3-GFP precursor in Arabidopsis OX:RSH3-GFP plants (Fig. 1E). Strikingly, the phosphoforms were lost rapidly upon TOR inhibition. Interestingly, we did not observe phosphoforms for mature RSH3-GFP (Fig. 1E, lower band), suggesting that TOR-dependent phosphorylation occurs only on the CTP which is subsequently cleaved and degraded after chloroplast import.

We then analyzed RSH3_39-221_ and found serine residues in TOR phosphorylation compatible contexts (Hsu et al., 2011; Van Leene et al., 2019) (Fig. S5A). When expressed in *N. benthamiana* several RSH3_39-221_-GFP phosphoforms accumulated under control conditions and disappeared upon TOR inhibition (Fig. 1F). Mutation of TOR compatible phosphosites in RSH3_39-221_-GFP almost completely abolished TOR-dependent phosphorylation. LC-MS analysis of immunoprecipitated RSH3_39-221_-GFP identified phosphorylation at 15 serine residues, including five putative TOR-dependent phosphosites (Fig. 1G). TOR-inhibition caused a dramatic reduction in the proportion of phosphorylated peptides, as expected. Together these results indicate that the RSH3 CTP is hyper-phosphorylated in a TOR-dependent manner, and that hyper-phosphorylation requires serine residues in canonical TOR-dependent contexts.

The region required for TOR interaction is found only in the NTR of RSH2/3 family members in plants and algae, and is absent from RSH1 and RSH4/CRSH enzymes, as well as from prokaryotic RSH enzymes (Fig. S5). The glaucophyte algae *Cyanophora paradoxa* RSH2/3 enzyme has an NTR-like region including predicted TOR dependent phosphosites, indicating possible origins prior to the divergence of green algae and land plants from glaucophytes, almost 2 billion years ago (Strassert et al., 2021). The RSH2/3 NTRs from Arabidopsis, tomato, and rice were all phosphorylated in a TOR-dependent manner (Fig. 1H). The *C. paradoxa* RSH2/3 NTR was not phosphorylated, suggesting either divergent TOR recognition or a later emergence of TOR-dependent regulation. Altogether, we show that the RSH3 NTR has ancient evolutionary origins, and phosphorylation by TOR is conserved at least among flowering plants.

**Fig. S5.**
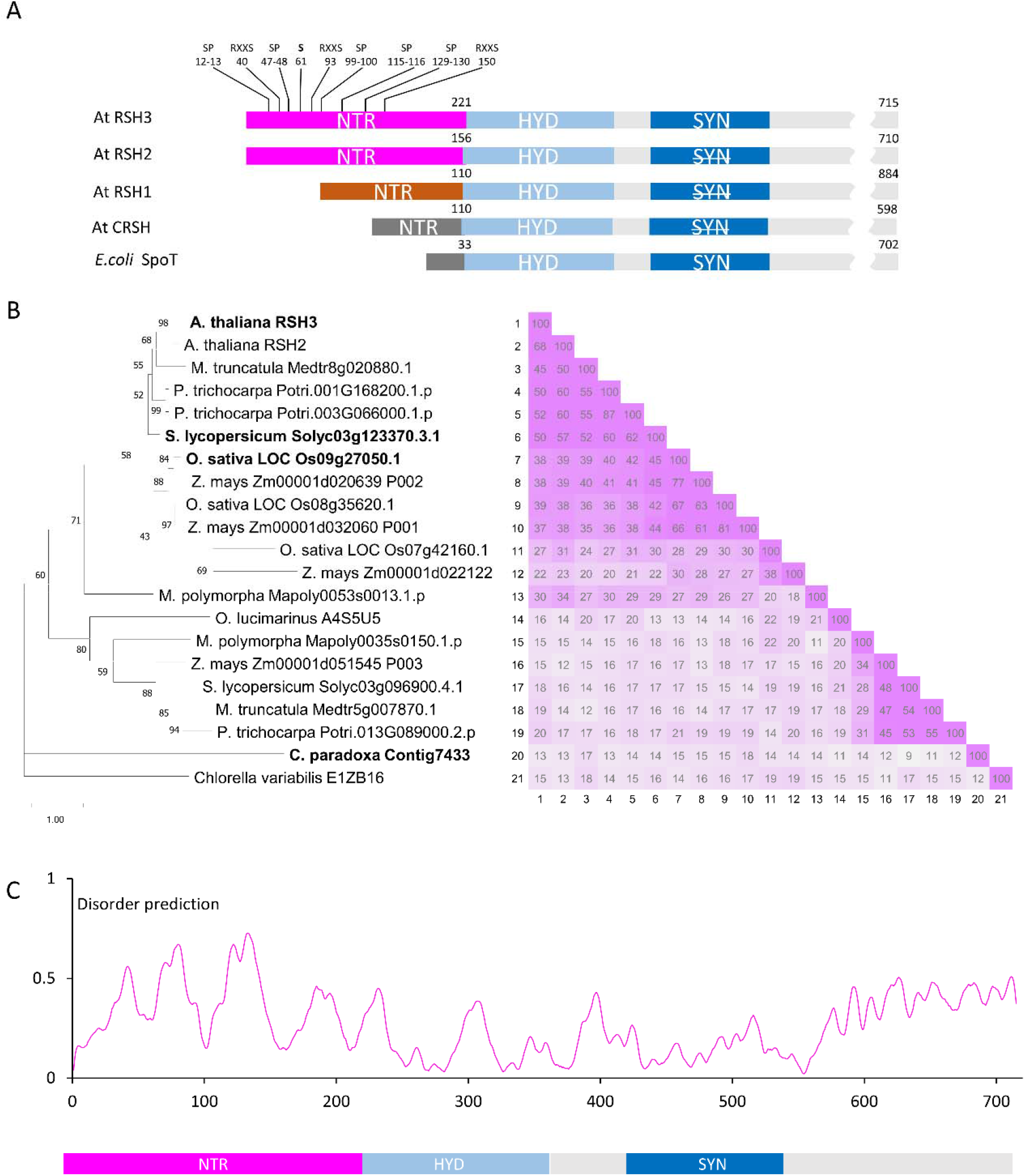
Features and evolution of the RSH NTR. (A) Domain structure of the four Arabidopsis RSH enzymes focused on the N-terminal and catalytic domains. Domains with the same colour show strong sequence homology. Serines in TOR-dependent phosphorylation contexts (SP, RXXS) are indicated above RSH3 (Van Leene et al., 2019). Prior experimental evidence existed for S61 phosphorylation (PhosPhAt 4.0)(Xi et al., 2021). (B) Maximum likelihood inference of evolutionary relationship between NTRs from RSH2/3 family enzymes found in plants, green algae and the glaucophyte *Cyanophora paradoxa*. NTRs tested for TOR-dependent phosphorylation are in bold. Boostrap support shown at branch nodes, and scale indicates substitutions per site. (C) Disorder prediction across RSH3, indicating higher levels of disorder within the NTR. IUPred3 (Erdős et al., 2021).

**Fig. S6.**
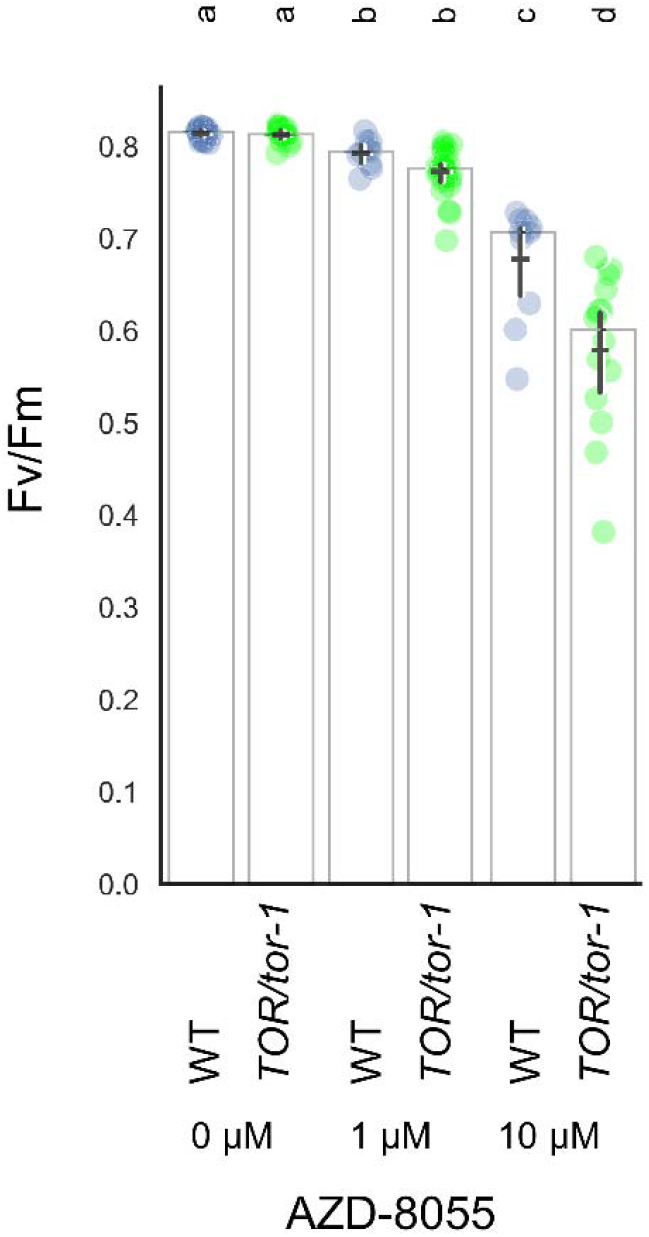
TOR-inhibition by AZD-8055 leads to a dose dependent decrease in Fv/Fm. Maximal efficiency of PSII (Fv/Fm) was measured in seedlings of wild type and TOR/*tor-1* heterozygous plants after 6 days growth on the indicated concentrations of AZD-8055. Graphs show mean (horizontal bar), median (column height) and 95% CI (vertical line). Lower-case letters indicate statistical groups.

We next sought to determine whether TOR regulates RSH-dependent ppGpp homeostasis by monitoring photosynthesis, a conserved target of ppGpp (Mehrez et al., 2022). Treatment of Arabidopsis seedlings with AZD caused a dose dependent drop in photosynthetic efficiency. TOR-haploinsufficient seedlings were more sensitive to AZD, indicating that this effect is TOR dependent (Fig. S6)(Montané and Menand, 2013). We then inhibited TOR in seedlings lacking the RSH1 ppGpp hydrolase (*rsh1-1*), or the main ppGpp synthetases RSH2 and RSH3 (*rsh2-1, rsh3-1, rsh*_*2,3*_)(Fig. 2A). TOR inhibition caused a sharp drop in the maximal efficacity of photosystem II (Fv/Fm) in the wild type, and this drop was more pronounced in *rsh1-1* seedlings that lack ppGpp hydrolase activity. In contrast, we observed progressive resistance to the effect of TOR inhibition in *rsh2-1, rsh3-1*, and finally *rsh*_*2,3*_, where the drop in Fv/Fm was strongly curtailed. We further confirmed the effect of TOR inhibition on ppGpp homeostasis by testing plants overexpressing the RSH1 ppGpp hydrolase (OX:RSH1) and OX:RSH3 plants that accumulate high ppGpp levels (Sugliani et al., 2016). OX:RSH3 seedlings were hypersensitive to TOR inhibition, while OX:RSH1 seedlings were resistant (Fig. 2B). OX:RSH3 adult plants were also hypersensitive to TOR inhibition (Fig. S7A,B), and, importantly, this was accompanied by a marked increase in ppGpp levels (Fig. 2C). The Fv/Fm of OX:RSH3 was also hypersensitive to nitrogen deprivation (Fig. S7C), a physiologically-relevant stress known to inhibit TOR activity (Liu et al., 2021). We therefore show that ppGpp accumulation mediated by the activities of RSH2, RSH3 and RSH1 is the main driver of photosynthesis repression following the inhibition of TOR.

**Fig. 2.**
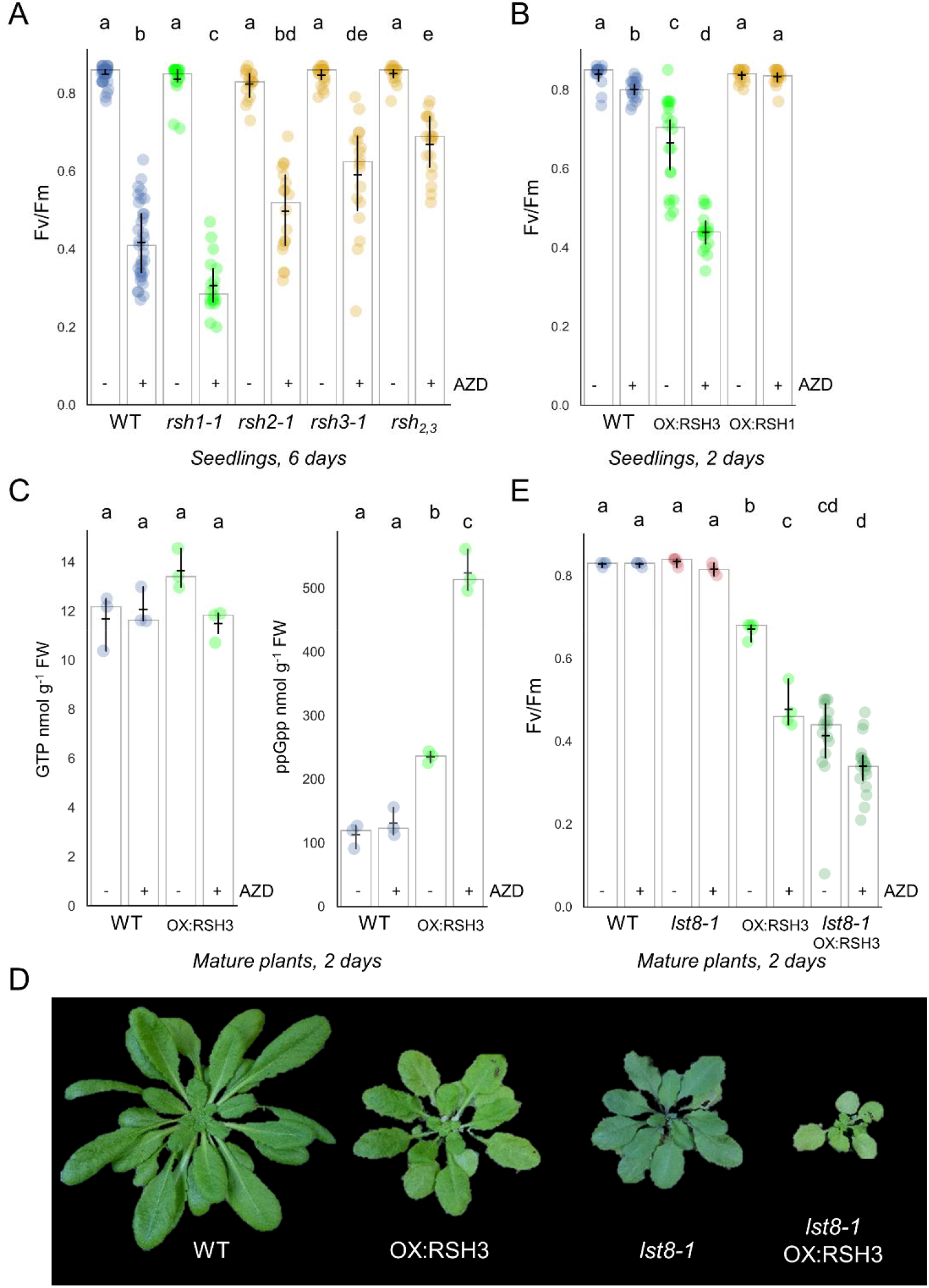
TOR activity regulates photosynthesis via RSH-dependent ppGpp synthesis. Maximal efficiency of PSII (Fv/Fm) was measured in seedlings of the indicated Arabidopsis lines treated ± 10 μM AZD for (A) 6 days (n=17-35 plants), and (B) 48 hours (n=17-20 plants). (C) Nucleotide quantification in adult plants treated ± 10 μM AZD for 48 hours (n=3 biological replicates). (D) Images of 5-week-old Arabidopsis wild-type and mutant plants. (E) Fv/Fm measurements of mature plants treated ± 10 μM AZD for 48 hrs (n=4-20). Graphs show mean (horizontal bar), median (column height) and 95% CI (vertical line). Lower-case letters indicate statistical groups.

Next, we investigated whether loss of LST8 in the *lst8-1* mutant affected the hypersensitivity of OX:RSH3 plants to AZD. Strikingly, *lst8-1* OX:RSH3 plants showed a severe growth and development phenotype that was stronger than in the parental lines (Fig. 2D). Despite phenotypic differences, mature wild type and *lst8-1* plants showed similar Fv/Fm ratios with or without exposure to AZD for 2 days (Fig. 2E). However, the Fv/Fm of untreated *lst8-1* OX:RSH3 dropped to the same level as in AZD-treated OX:RSH3, strongly suggesting that the absence of LST8 leads to constitutive activation of RSH3 via a reduction in either TOR activity or the quantity of interaction partner. (Fig. 2E). In agreement, *lst8-1* OX:RSH3 appeared less sensitive to AZD treatment.

**Fig. S7:**
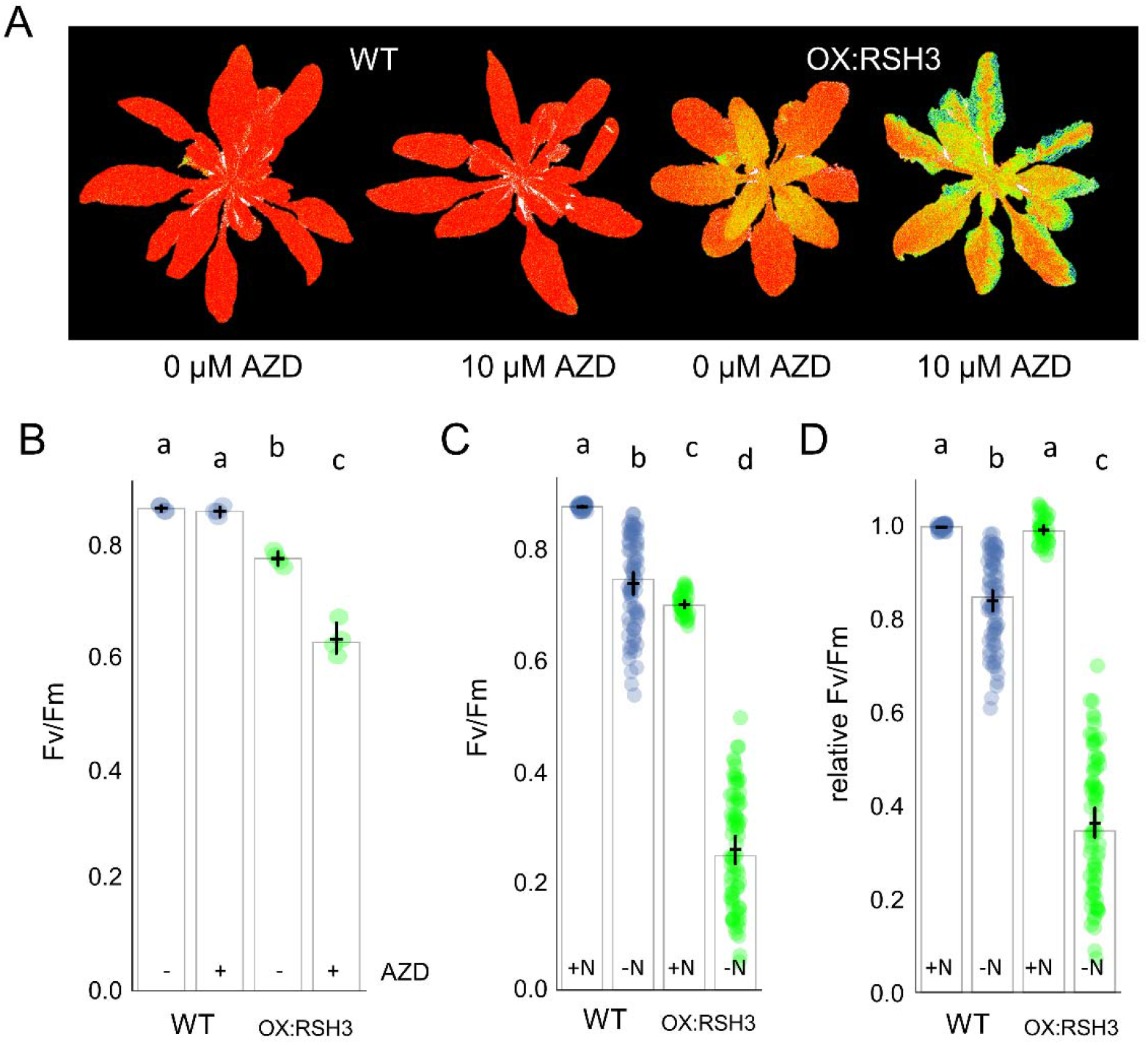
OX:RSH3 plants are hypersensitive to TOR inhibition and nitrogen starvation. Maximal efficiency of PSII (Fv/Fm) was (A) imaged and (B) quantified in 5-week-old wild type and OX:RSH3-GFP plants treated with ± 10 μM AZD for 48 hours (n= 4 plants). Fv/Fm was measured in wild type and OX:RSH3-GFP seedlings following transfer to a nitrogen-replete (+N) or nitrogen-limited medium (-N) for 16 days and is presented as either unmodified Fv/Fm (C), or as relative Fv/Fm (D) where values were normalized to the corresponding +N control (+N, n=36 plants; -N n=72 plants). Graphs show mean (horizontal bar), median (column height) and 95% CI (vertical line). Lower-case letters indicate statistical groups.

The previous experiments lead us to ask whether the RSH3 NTR is required for the regulation of ppGpp signaling by TOR. We therefore generated plants overexpressing RSH3-GFP (OX:RSH3-GFP), or RSH3-GFP with a truncated NTR that is no longer able to interact with LST8 (OX:SSU-RSH3_65-END_-GFP)(Fig. 1D, S2). These lines were created in the *rsh*_*2,3*_ background to prevent interference by the upregulation of *RSH2/3* transcription that can occur under stress conditions (Romand et al., 2022). As expected, *rsh*_*2,3*_ OX:RSH3-GFP was hypersensitive to TOR inhibition (Fig. 3A). Strikingly, however, *rsh*_*2,3*_ seedlings expressing SSU-RSH3_65-END_-GFP, which cannot interact with LST8 (Fig. 1D) were completely insensitive to TOR inhibition. These results, together with the constitutive activation of RSH3 in the OX:RSH3-GFP *lst8* plants (Fig. 2E), demonstrate that the interaction between LST8 and the NTR is required for repression of RSH3 activity *in planta*.

**Fig. 3.**
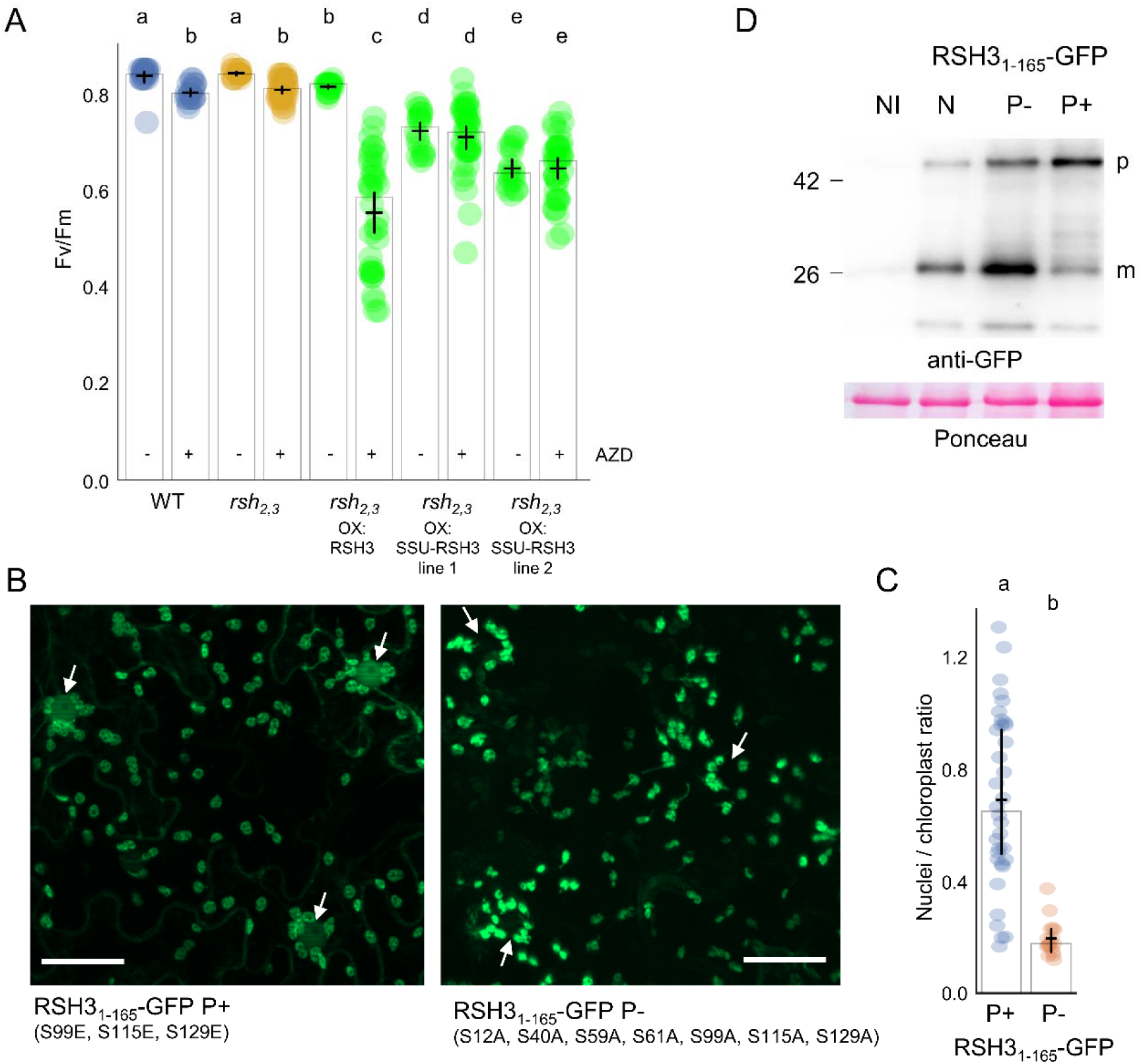
TOR regulates RSH3 activity and cellular distribution via the RSH3 NTR. (A) Maximal efficiency of PSII (Fv/Fm) was measured in Arabidopsis seedlings of the indicated lines treated ± 10 μM AZD for 48 hours (n=18-36 plants). RSH3, RSH3-GFP; SSU-RSH3, SSU-RSH3_65-END_-GFP. (B) Fluorescence microscopy images of *N. benthamiana* leaves expressing RSH3_1-165_-GFP P+ and RSH3_1-165_-GFP P-. White arrows indicate nuclei surrounded by chloroplasts. Scale bar, 50 μm. (C) Quantification of GFP fluorescence in nuclei against GFP fluorescence in nuclei-proximal chloroplasts, n=50 nuclei. (D) Immunoblots of protein extracts from *N. benthamiana* expressing the indicated proteins or the non-inoculated control (NI). p, precursor protein; m, mature protein after chloroplast import. Graphs show mean (horizontal bar), median (column height) and 95% CI (vertical line). Lower-case letters indicate statistical groups.

We reasoned that phosphorylation of the RSH3 CTP might affect RSH3 activity by reducing accumulation in the chloroplast. Indeed, phosphorylation of chloroplast precursors is known to impede import (Waegemann and Soll, 1996). We therefore analyzed the localization of phosphomimic and phospho-null mutants of RSH3_1-165_-GFP. Phosphomimic RSH3_1-165_-GFP showed both chloroplastic and nucleocytosolic localization, while the native and phospho-null form showed only chloroplastic localization (Fig. 3C, S4B). Furthermore, a greater proportion of mature protein compared to precursor was observed for phospho-null RSH3_1-165_-GFP, while the phosphomimic accumulated a lower proportion of mature protein (Fig. 3D). Full-length phosphomimic RSH3-GFP also showed a reduced chloroplast localization (Fig. S8A). Altogether, these results support a model whereby TOR-dependent phosphorylation attenuates the chloroplast localization of RSH3. Inactivation of TOR, either artificially or via nitrogen starvation, leads to rapid RSH3 dephosphorylation and increased chloroplast localization to allow more ppGpp synthesis, which in turn inhibits photosynthesis. The inherent instability of RSH3 (Fig. S3B) somewhat masks the increased level of mature RSH3 in the chloroplast (Fig. 1E). The small and transient pool of mature RSH3 might be highly sensitive to fluctuations in import of the precursor protein.

**Fig. S8.**
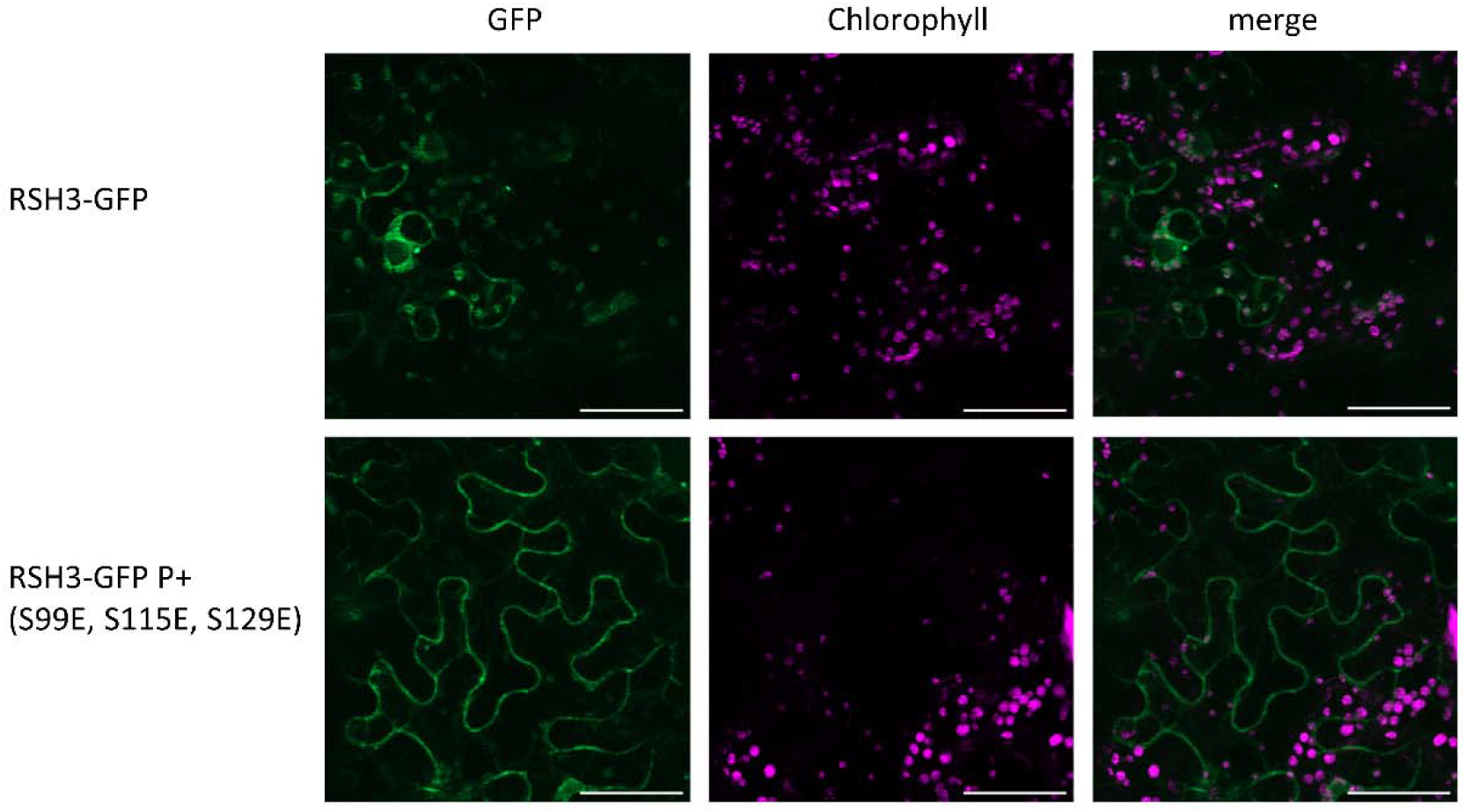
The phosphostatus of full length RSH3-GFP influences cellular distribution. Fluorescence microscopy images of *N. benthamiana* expressing RSH3-GFP or RSH3-GFP P+. Scale bar, 50 μm.

In conclusion, we reveal a conserved post-translational mechanism for the regulation of chloroplast function by the TOR kinase in plants. This mechanism, which is independent and distinct from transcriptional pathways of chloroplast (Dong et al., 2015; Sun et al., 2016; Imamura et al., 2018; Han et al., 2022), may allow rapid co-ordination of the nucleocytosolic and chloroplast compartments. Such co-ordination may be particularly important during episodes of stress where growth and photosynthesis must be downregulated in lockstep to prevent photooxidative damage (Romand et al., 2022). Together with a recent report that TOR promotes accumulation of the chloroplast β-AMYLASE1 in stomatal chloroplasts (Han et al., 2022) our work sets a new precedent for the regulation of organellar function by TOR, and provides a molecular mechanism for explaining the TOR-dependent regulation of photosynthesis.

## Acknowledgements

We thank Marina Siponen from the BIAM ProteinTec platform for providing α-GFP nanobody:Halo:His6 and Julia Bartoli for providing nucleotide standards. Nucleotide measurements were performed on the IJPB Plant Observatory technological platform that is supported by Saclay Plant Sciences-SPS (ANR-17-EUR-0007), microscopy experiments were performed on theLJPiCSL-FBI core faculty member of the France-BioImaging national research infrastructure (ANR-10-INBS-04). We acknowledge funding form the Erasmus+ International Credit Mobility for MM and funding from the Agence National de la Recherche (ANR-22-CE20-0033, ANR-17-CE13-0005).

## Contributions

Stefano D’Alessandro Conception and design, Method development, Acquisition, analysis, interpretation and visualisation of data, Writing original draft, Review and editing

Florent Velay Conception and design, Method development, Acquisition, analysis, validation and interpretation of data, Review and editing

Regine Lebrun Method development, Acquisition and analysis of LC-MS data, Review and editing Marwa Mehrez Acquisition and analysis of data from plant physiology assays.

Shanna Romand Acquisition and analysis of data from plant physiology assays, nucleotide quantification

Rim Saadouni Method development, Acquisition and analysis of LC-MS data Celine Forzani Acquisition and analysis of Y2H data

Sylvie Citerne Acquisition and analysis of nucleotide quantification data.

Marie-Helene Montane Conception and design, Review and editing

Christophe Robaglia Conception and design, Review and editing

Benoit Menand Conception and design, Review and editing

Christian Meyer Conception and design, Acquisition and analysis of Y2H data, Review and editing

Ben Field Conception and design, Method development, Acquisition, analysis, interpretation and visualization of data, Writing original draft, Review and editing

## References

Almagro Armenteros, J.J., Salvatore, M., Emanuelsson, O., Winther, O., von Heijne, G., Elofsson, A., and Nielsen, H. (2019). Detecting sequence signals in targeting peptides using deep learning. Life Sci Alliance 2: e201900429.

Bange, G., Brodersen, D.E., Liuzzi, A., and Steinchen, W. (2021). Two P or Not Two P: Understanding Regulation by the Bacterial Second Messengers (p)ppGpp. Annu Rev Microbiol.

Bienvenut, W.V., Sumpton, D., Martinez, A., Lilla, S., Espagne, C., Meinnel, T., and Giglione, C. (2012). Comparative Large Scale Characterization of Plant versus Mammal Proteins Reveals Similar and Idiosyncratic N-α-Acetylation Features *. Molecular & Cellular Proteomics 11.

Burkart, G.M. and Brandizzi, F. (2021). A Tour of TOR Complex Signaling in Plants. Trends in Biochemical Sciences 46: 417–428.

D’Alessandro, S. (2022). Coordination of Chloroplast Activity with Plant Growth: Clues Point to TOR. Plants 11: 803.

Dong, P., Xiong, F., Que, Y., Wang, K., Yu, L., Li, Z., and Maozhi, R. (2015). Expression profiling and functional analysis reveals that TOR is a key player in regulating photosynthesis and phytohormone signaling pathways in Arabidopsis. Front. Plant Sci. 6.

Han, C., Hua, W., Li, J., Qiao, Y., Yao, L., Hao, W., Li, R., Fan, M., De Jaeger, G., Yang, W., and Bai, M.-Y. (2022). TOR promotes guard cell starch degradation by regulating the activity of β-AMYLASE1 in Arabidopsis. The Plant Cell 34: 1038–1053.

Hsu, P.P., Kang, S.A., Rameseder, J., Zhang, Y., Ottina, K.A., Lim, D., Peterson, T.R., Choi, Y., Gray, N.S., Yaffe, M.B., Marto, J.A., and Sabatini, D.M. (2011). The mTOR-regulated phosphoproteome reveals a mechanism of mTORC1-mediated inhibition of growth factor signaling. Science 332: 1317–1322.

Imamura, S., Nomura, Y., Takemura, T., Pancha, I., Taki, K., Toguchi, K., Tozawa, Y., and Tanaka, K. (2018). The checkpoint kinase TOR (target of rapamycin) regulates expression of a nuclear-encoded chloroplast RelA-SpoT homolog (RSH) and modulates chloroplast ribosomal RNA synthesis in a unicellular red alga. The Plant Journal 94: 327–339.

Liu, Y., Duan, X., Zhao, X., Ding, W., Wang, Y., and Xiong, Y. (2021). Diverse nitrogen signals activate convergent ROP2-TOR signaling in Arabidopsis. Developmental Cell 56: 1283–1295.e5.

Maekawa, M., Honoki, R., Ihara, Y., Sato, R., Oikawa, A., Kanno, Y., Ohta, H., Seo, M., Saito, K., and Masuda, S. (2015). Impact of the plastidial stringent response in plant growth and stress responses. Nature Plants 1: 1–7.

Mehrez, M., Romand, S., and Field, B. (2022). New perspectives on the molecular mechanisms of stress signalling by the nucleotide guanosine tetraphosphate (ppGpp), an emerging regulator of photosynthesis in plants and algae. New Phytologist.

Montané, M.-H. and Menand, B. (2013). ATP-competitive mTOR kinase inhibitors delay plant growth by triggering early differentiation of meristematic cells but no developmental patterning change. J Exp Bot 64: 4361–4374.

Pacheco, J.M., Canal, M.V., Pereyra, C.M., Welchen, E., Martínez-Noël, G.M.A., and Estevez, J.M. (2021). The tip of the iceberg: emerging roles of TORC1, and its regulatory functions in plant cells. Journal of Experimental Botany 72: 4085–4101.

Romand, S. et al. (2022). A guanosine tetraphosphate (ppGpp) mediated brake on photosynthesis is required for acclimation to nitrogen limitation in Arabidopsis. eLife 11: e75041.

Strassert, J.F.H., Irisarri, I., Williams, T.A., and Burki, F. (2021). A molecular timescale for eukaryote evolution with implications for the origin of red algal-derived plastids. Nat Commun 12: 1879.

Sugliani, M., Abdelkefi, H., Ke, H., Bouveret, E., Robaglia, C., Caffarri, S., and Field, B. (2016). An Ancient Bacterial Signaling Pathway Regulates Chloroplast Function to Influence Growth and Development in Arabidopsis. The Plant Cell 28: 661–679.

Sun, L., Yu, Y., Hu, W., Min, Q., Kang, H., Li, Y., Hong, Y., Wang, X., and Hong, Y. (2016). Ribosomal protein S6 kinase1 coordinates with TOR-Raptor2 to regulate thylakoid membrane biosynthesis in rice. Biochimica et Biophysica Acta (BBA) - Molecular and Cell Biology of Lipids 1861: 639–649.

Upadhyaya, S. and Rao, B.J. (2019). Reciprocal regulation of photosynthesis and mitochondrial respiration by TOR kinase in Chlamydomonas reinhardtii. Plant Direct 3: e00184–e00184.

Van Leene, J. et al. (2019). Capturing the phosphorylation and protein interaction landscape of the plant TOR kinase. Nature Plants 5: 316–327.

Waegemann, K. and Soll, J. (1996). Phosphorylation of the Transit Sequence of Chloroplast Precursor Proteins (*). Journal of Biological Chemistry 271: 6545–6554.

Xi, L., Zhang, Z., and Schulze, W.X. (2021). PhosPhAt 4.0: An Updated ArabidopsisArabidopsis Database for Searching Phosphorylation Sites and Kinase-Target Interactions. In Plant Phosphoproteomics: Methods and Protocols, X.N. Wu, ed, Methods in Molecular Biology. (Springer US: New York, NY), pp. 189–202.

Zhang, Y., Song, G., Lal, N.K., Nagalakshmi, U., Li, Y., Zheng, W., Huang, P., Branon, T.C., Ting, A.Y., Walley, J.W., and Dinesh-Kumar, S.P. (2019). TurboID-based proximity labeling reveals that UBR7 is a regulator of N NLR immune receptor-mediated immunity. Nat Commun 10: 1–17.

