## Supplementary Data Files for "Post-translational regulation of photosynthetic activity via the TOR kinase in plants": Fig1B.pptx

### Slide 1
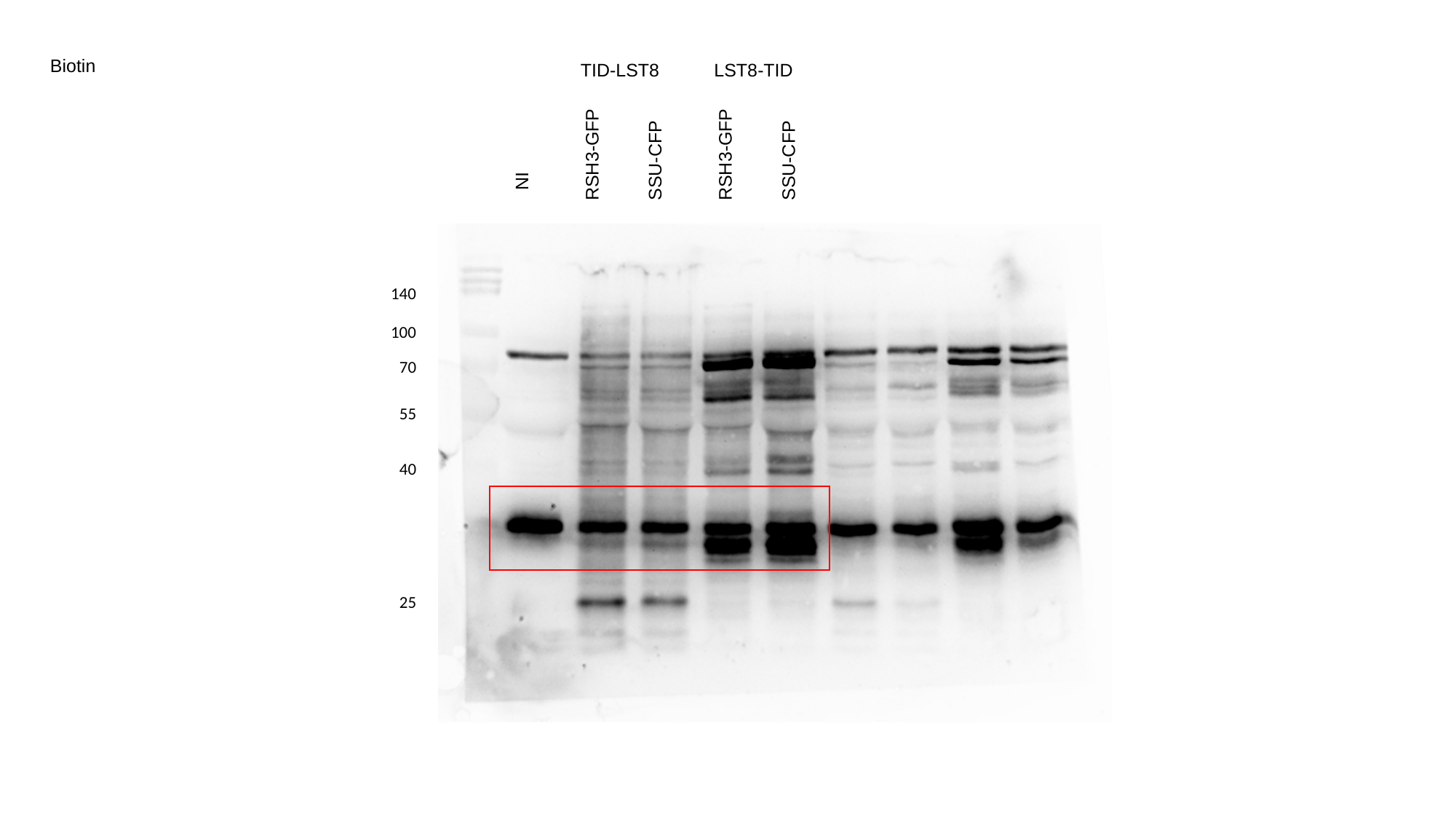

Biotin
TID-LST8
LST8-TID
RSH3-GFP
RSH3-GFP
SSU-CFP
SSU-CFP
NI
140
100
70
55
40
25

### Slide 2
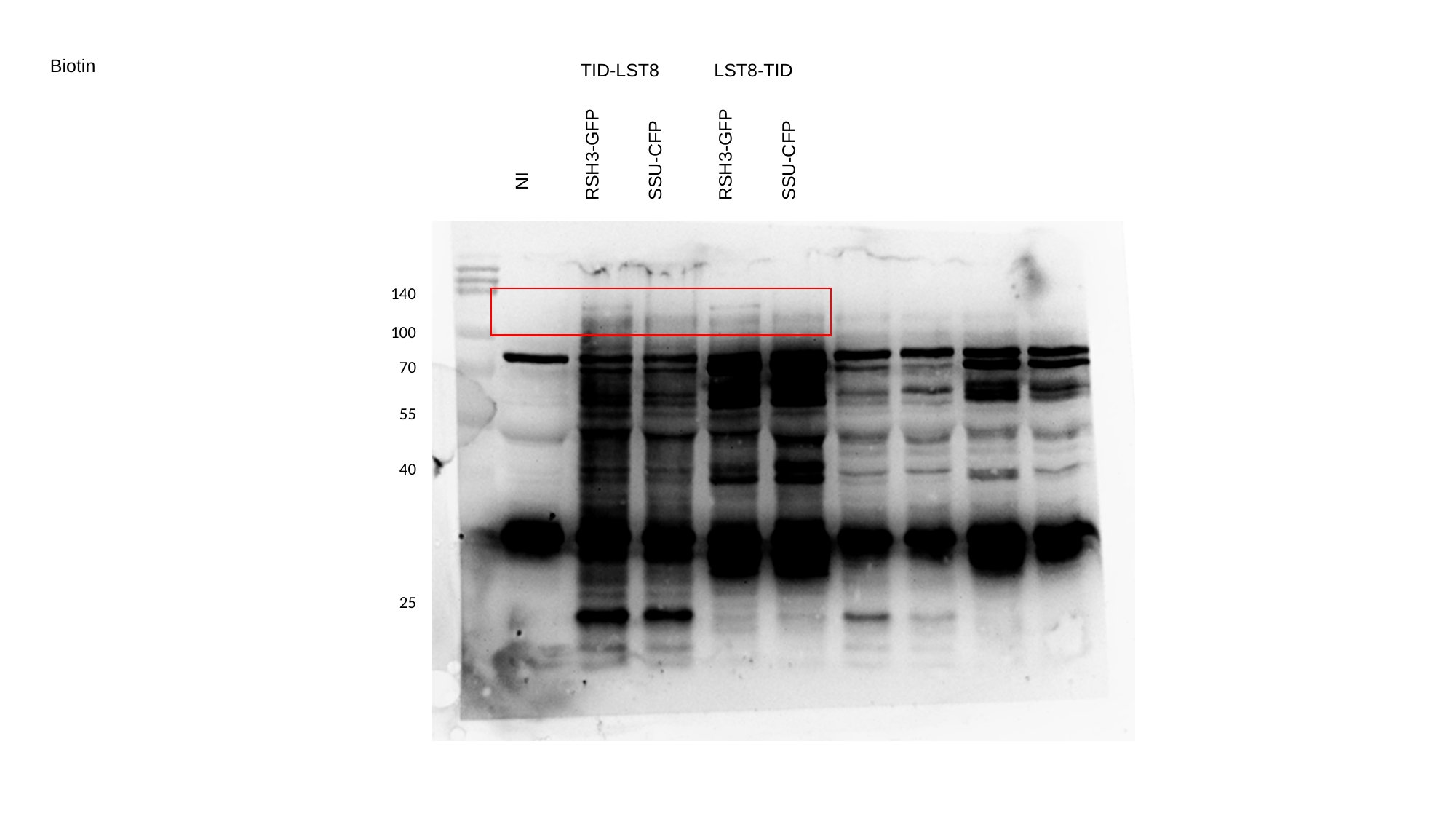

Biotin
TID-LST8
LST8-TID
RSH3-GFP
RSH3-GFP
SSU-CFP
SSU-CFP
NI
140
100
70
55
40
25

### Slide 3
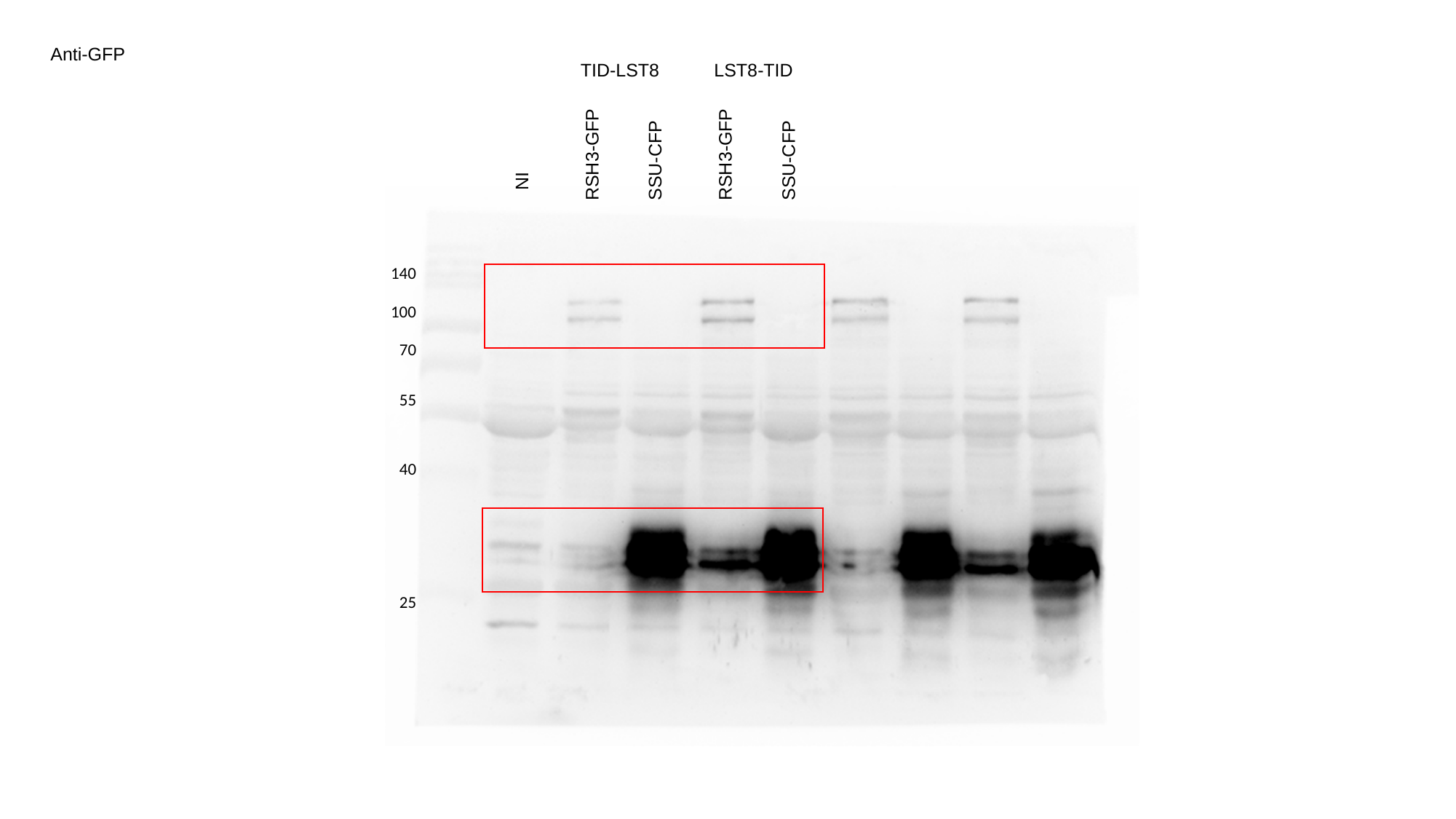

Anti-GFP
TID-LST8
LST8-TID
RSH3-GFP
RSH3-GFP
SSU-CFP
SSU-CFP
NI
140
100
70
55
40
25

### Slide 4
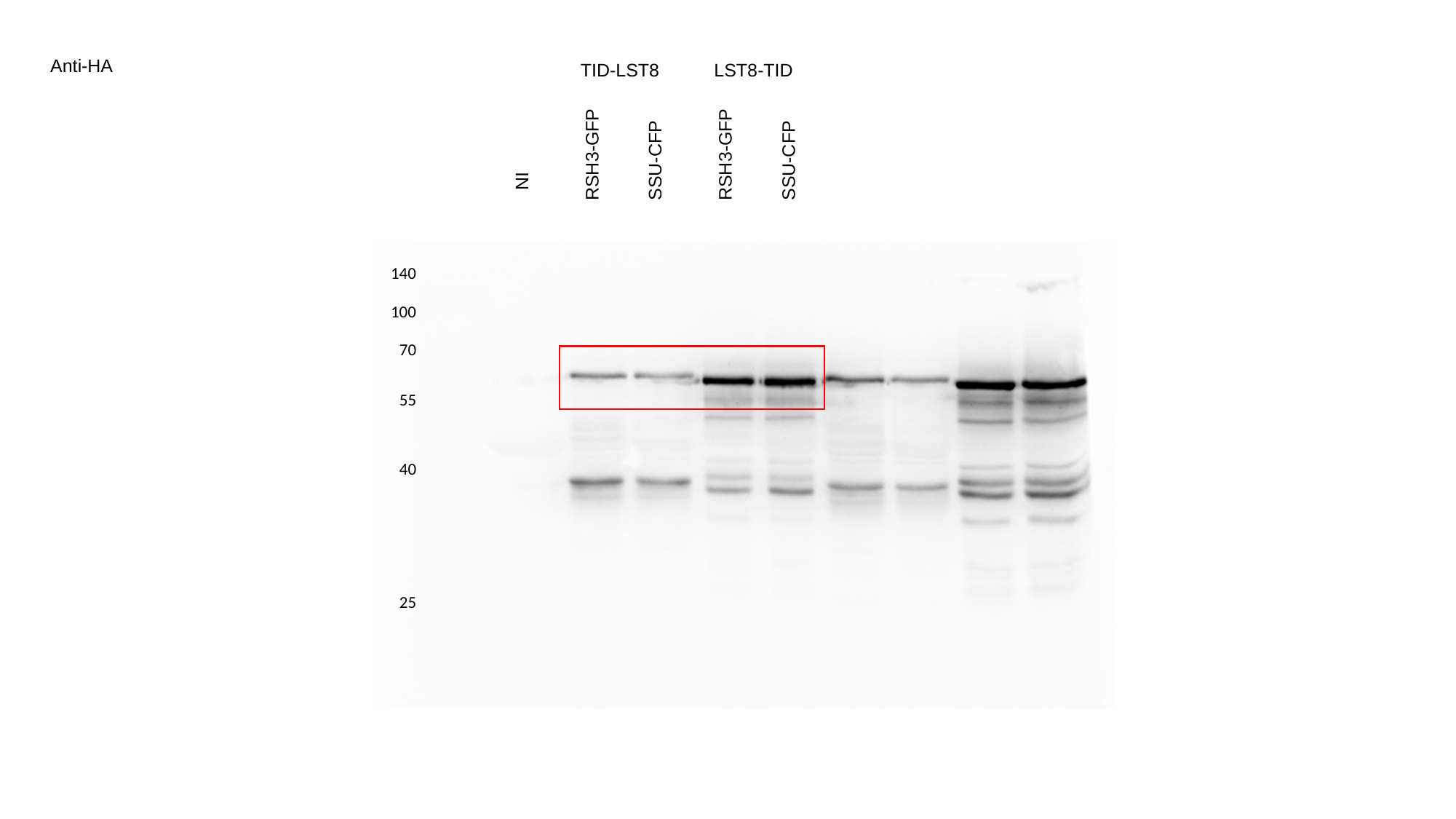

Anti-HA
TID-LST8
LST8-TID
RSH3-GFP
RSH3-GFP
SSU-CFP
SSU-CFP
NI
140
100
70
55
40
25
