## Supplementary Data Files for "Post-translational regulation of photosynthetic activity via the TOR kinase in plants": Fig1D.pptx

### Slide 1
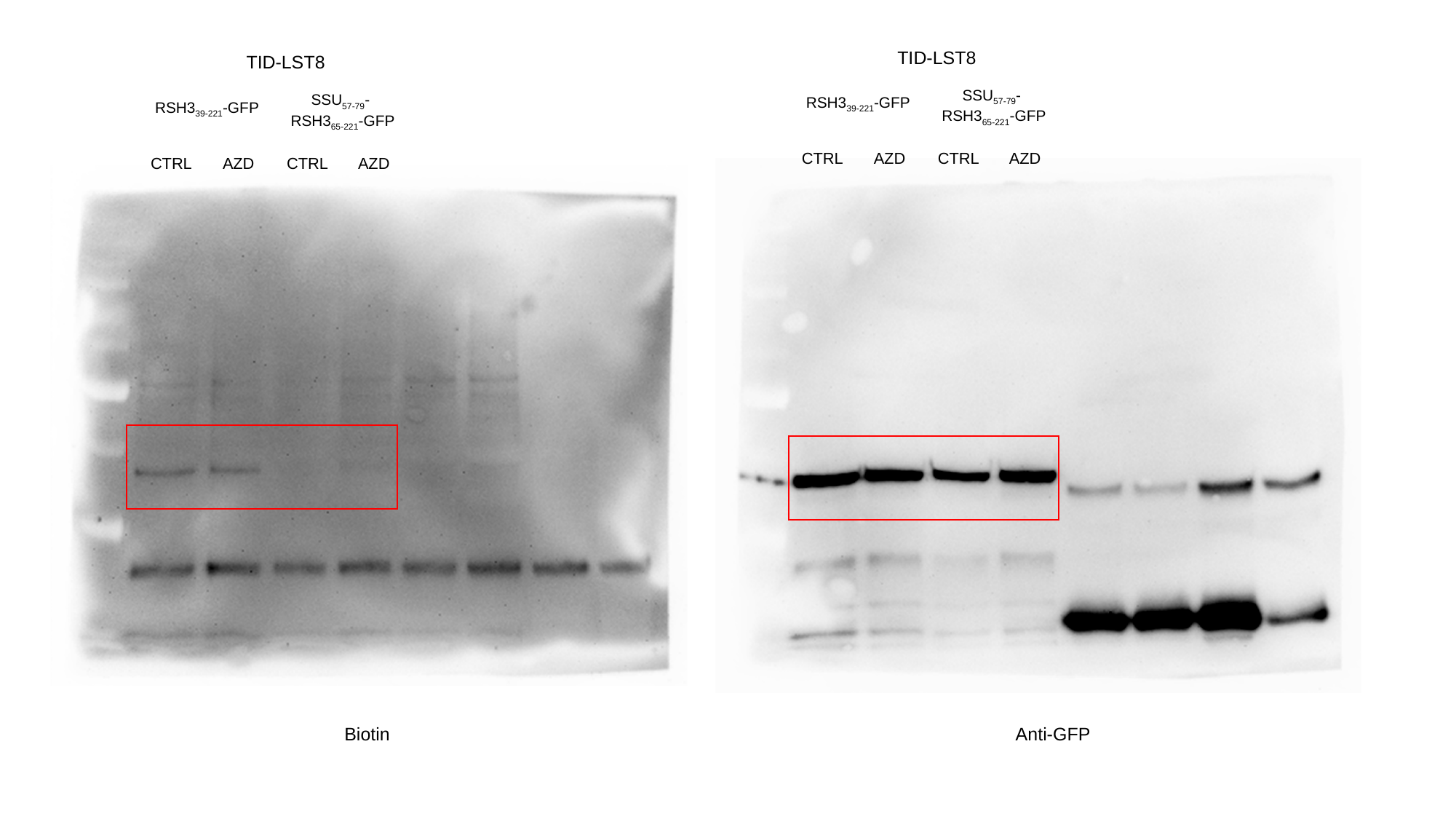

TID-LST8
SSU_57-79_-
RSH3_65-221_-GFP
RSH3_39-221_-GFP
CTRL
AZD
CTRL
AZD
TID-LST8
SSU_57-79_-
RSH3_65-221_-GFP
RSH3_39-221_-GFP
CTRL
AZD
CTRL
AZD
Anti-GFP
Biotin
