## Supplementary Data Files for "Post-translational regulation of photosynthetic activity via the TOR kinase in plants": Fig1F.pptx

### Slide 1
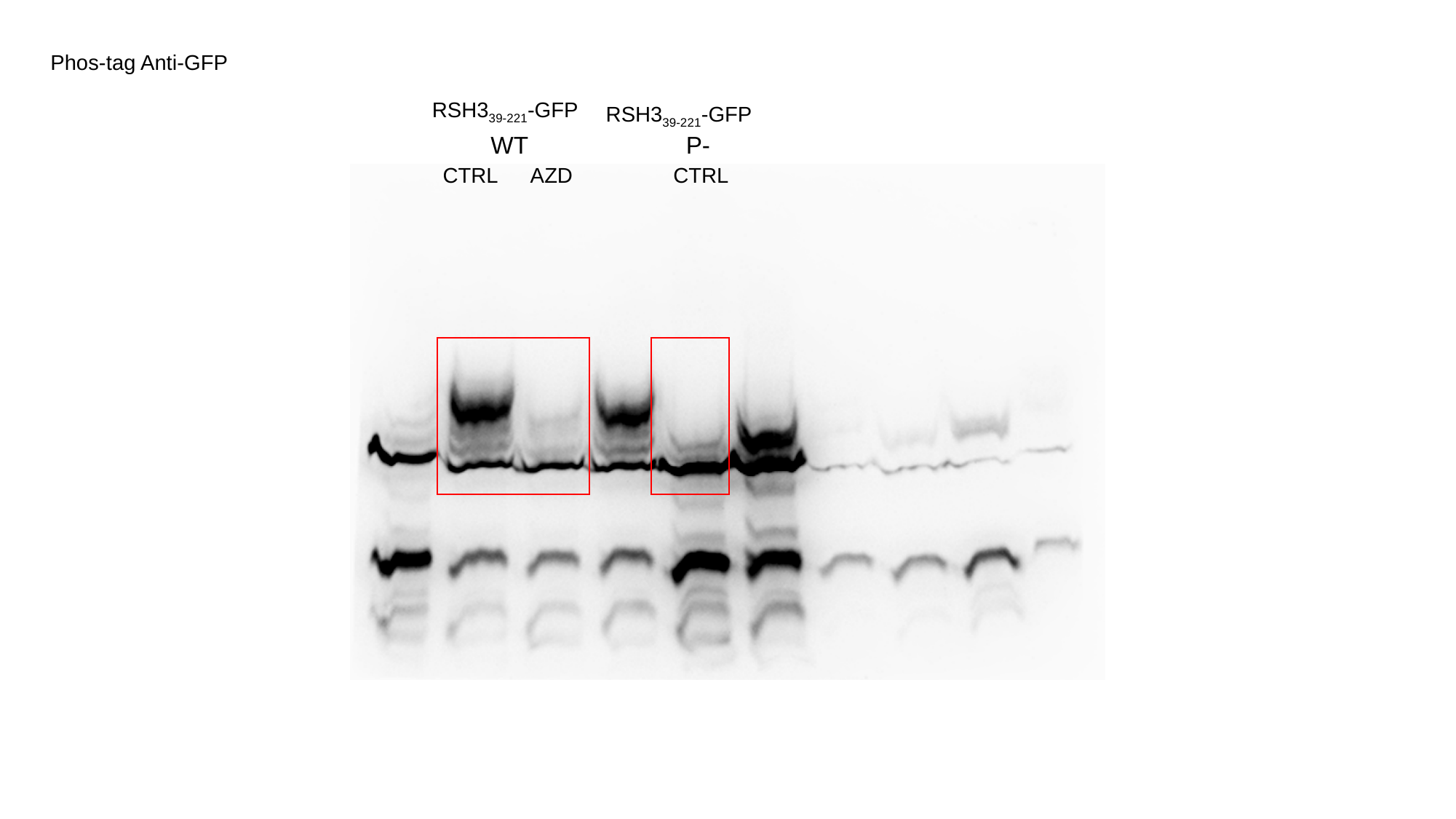

Phos-tag Anti-GFP
RSH3_39-221_-GFP
RSH3_39-221_-GFP
WT
P-
CTRL
CTRL
AZD

### Slide 2
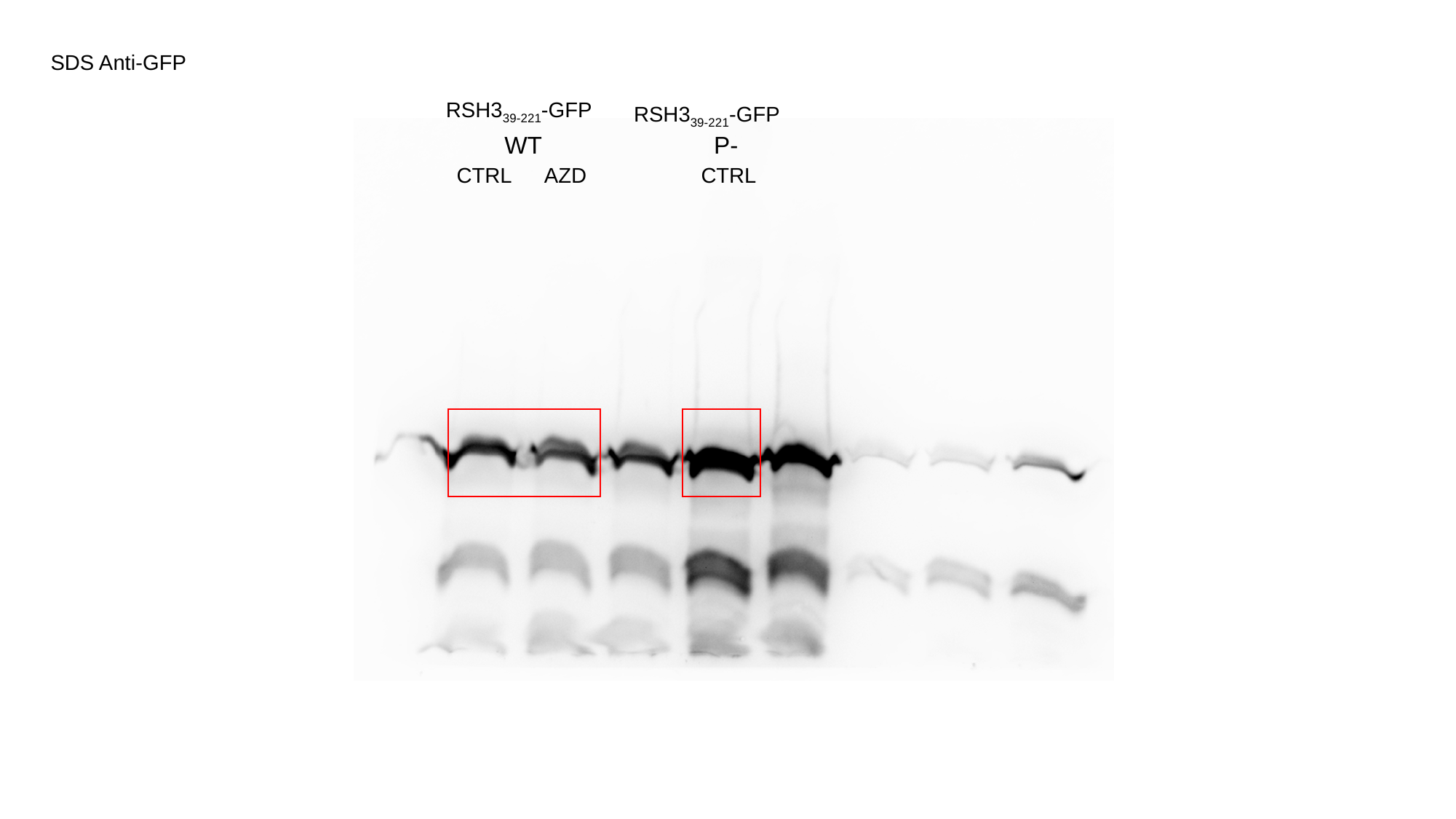

SDS Anti-GFP
RSH3_39-221_-GFP
RSH3_39-221_-GFP
WT
P-
CTRL
CTRL
AZD
