## Supplementary Data Files for "Post-translational regulation of photosynthetic activity via the TOR kinase in plants": RSH3(39-221) P positions.pptx

### Slide 1
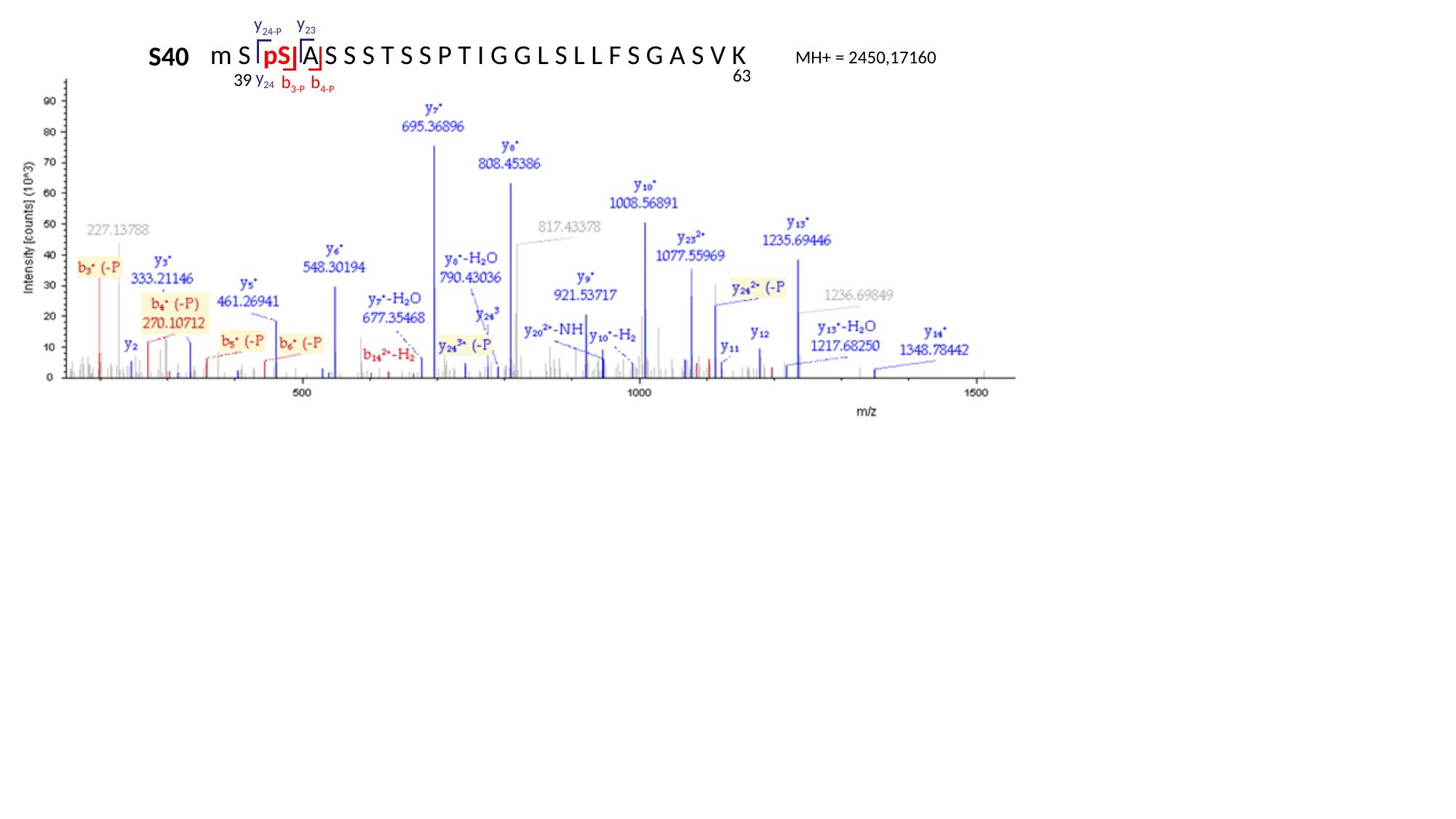

y_23_
y_24-P_
m S pS A S S S T S S P T I G G L S L L F S G A S V K
S40
MH+ = 2450,17160
63
y_24_
39
b_4-P_
b_3-P_

### Slide 2
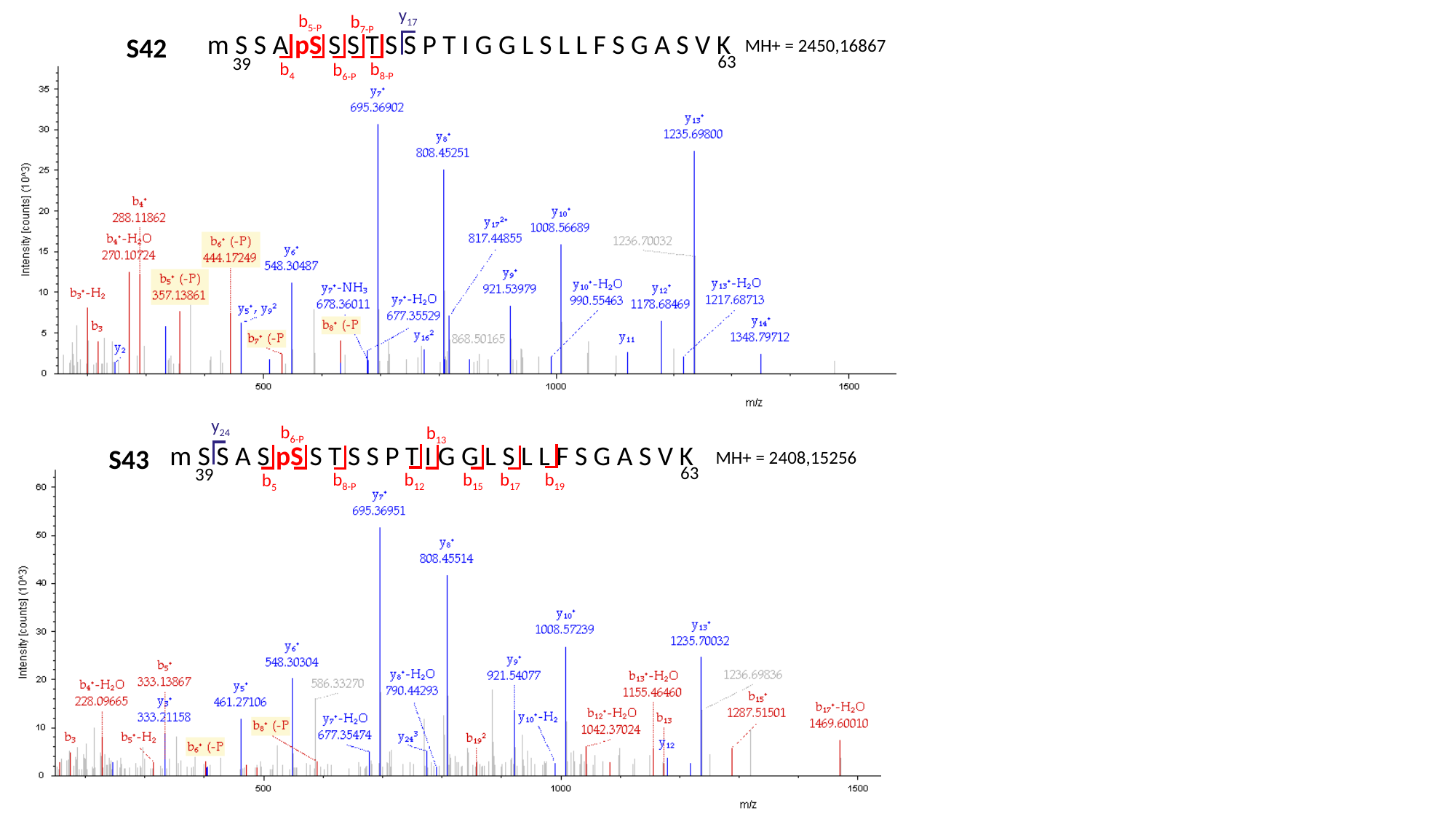

y_17_
b_5-P_
b_7-P_
m S S A pS S S T S S P T I G G L S L L F S G A S V K
S42
MH+ = 2450,16867
63
39
b_4_
b_8-P_
b_6-P_
y_24_
b_6-P_
b_13_
m S S A S pS S T S S P T I G G L S L L F S G A S V K
S43
MH+ = 2408,15256
63
39
b_8-P_
b_19_
b_15_
b_12_
b_17_
b_5_

### Slide 3
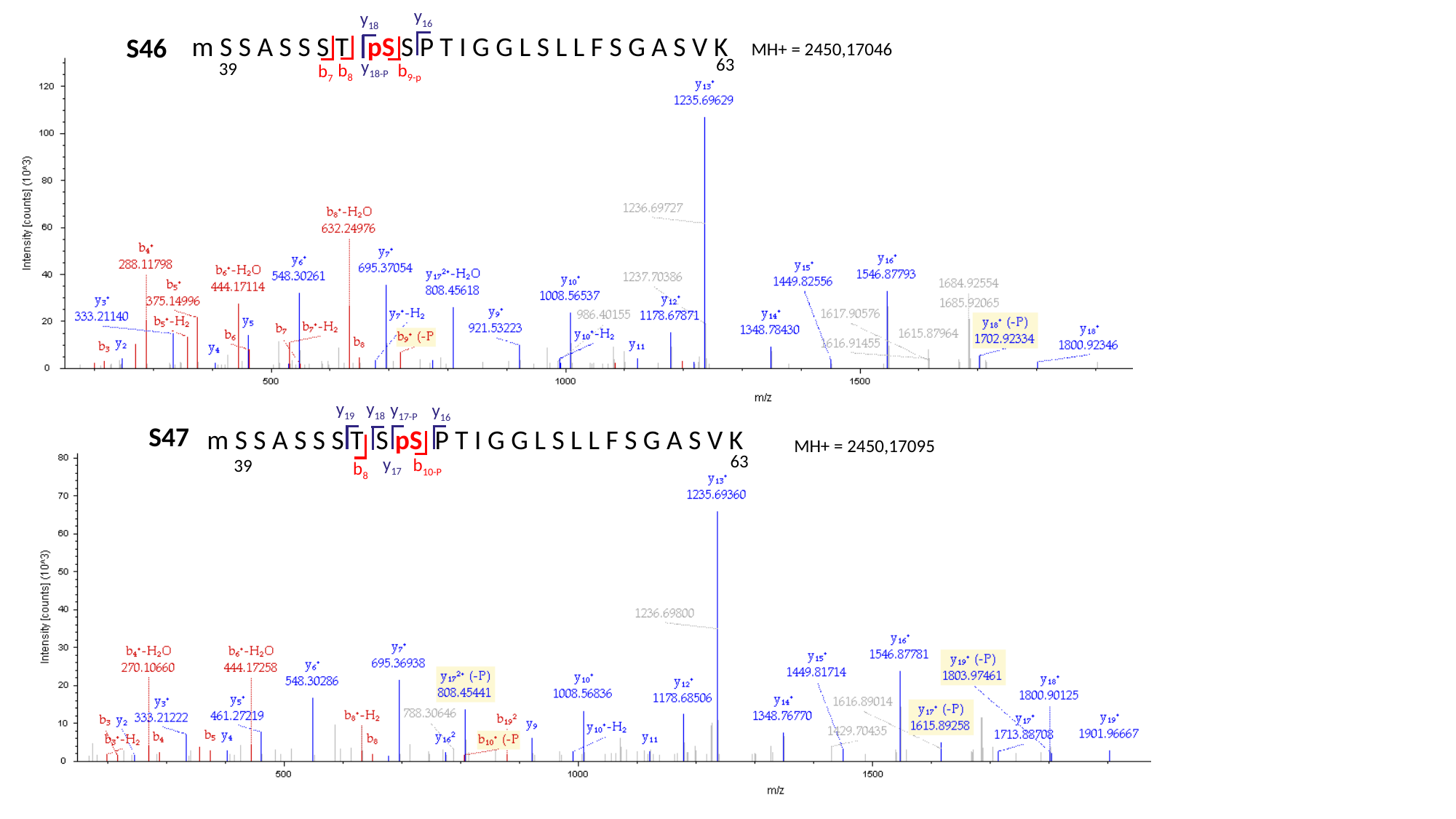

y_16_
y_18_
m S S A S S S T pS S P T I G G L S L L F S G A S V K
S46
MH+ = 2450,17046
63
y_18-P_
39
b_8_
b_9-p_
b_7_
y_19_
y_18_
y_17-P_
y_16_
S47
m S S A S S S T S pS P T I G G L S L L F S G A S V K
MH+ = 2450,17095
63
y_17_
b_10-P_
39
b_8_

### Slide 4
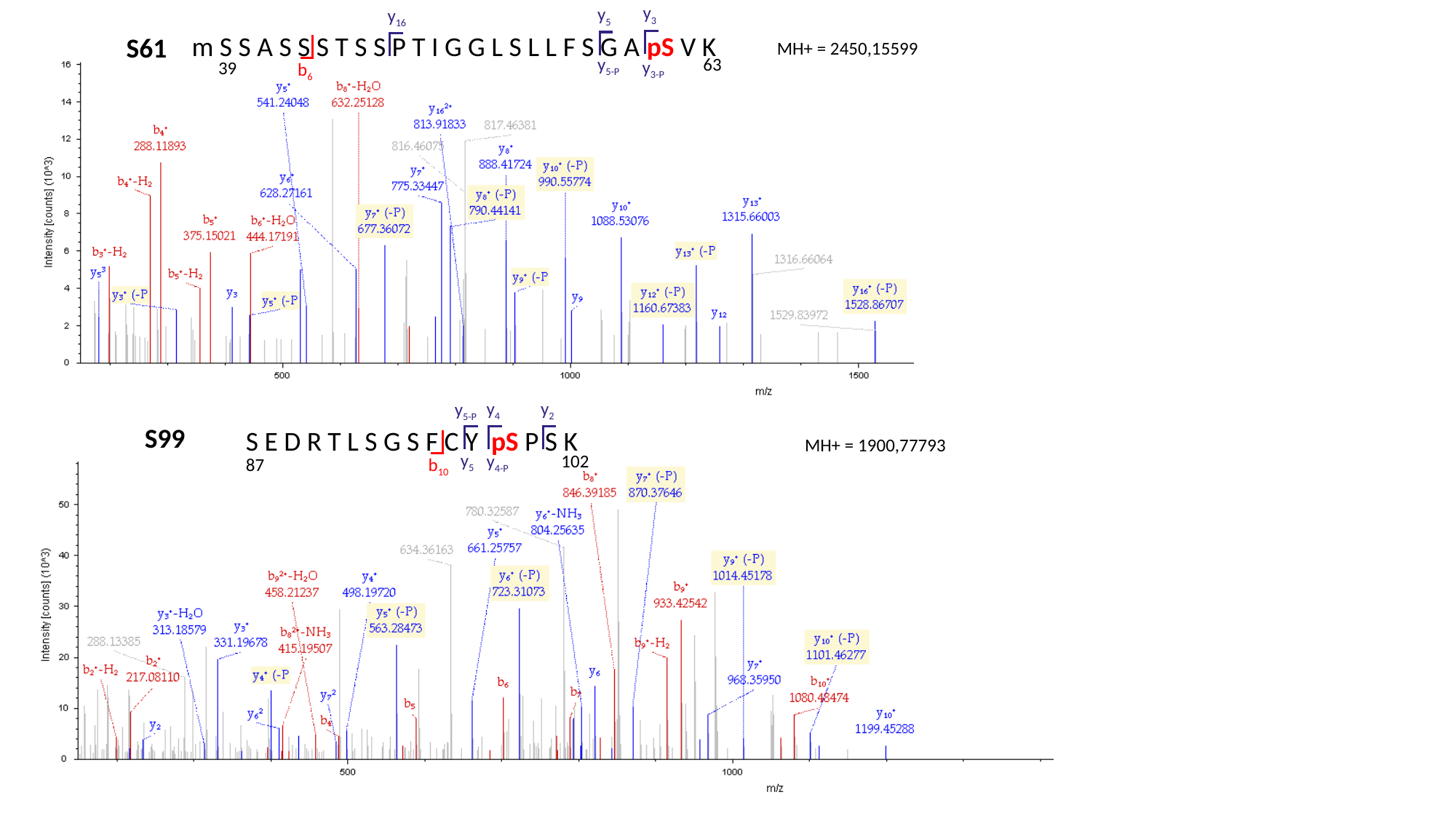

y_3_
y_5_
y_16_
m S S A S S S T S S P T I G G L S L L F S G A pS V K
S61
MH+ = 2450,15599
y_5-P_
63
y_3-P_
39
b_6_
y_2_
y_4_
y_5-P_
S99
S E D R T L S G S F C Y pS P S K
MH+ = 1900,77793
y_5_
y_4-P_
102
87
b_10_

### Slide 5
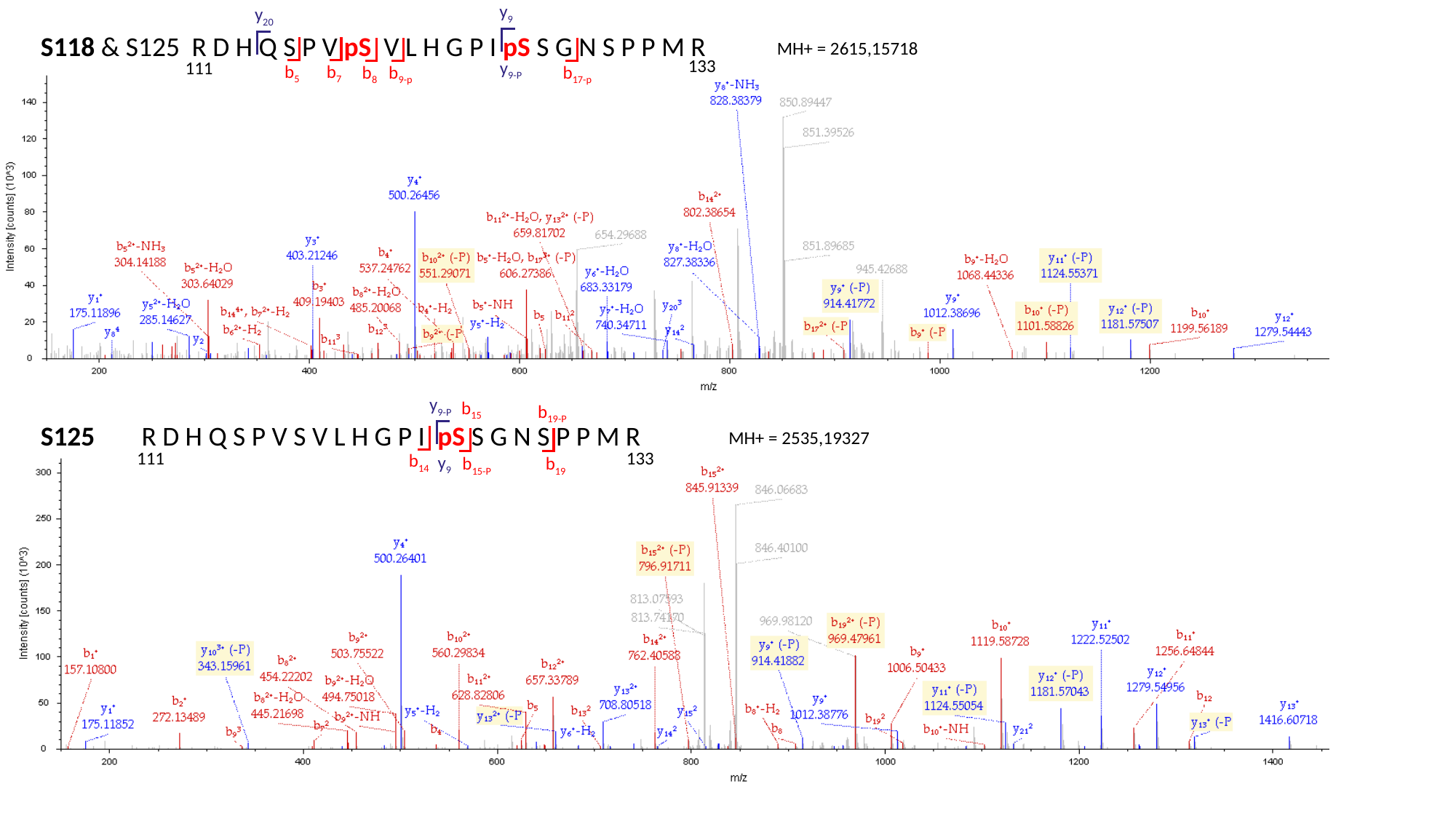

y_9_
y_20_
S118 & S125
R D H Q S P V pS V L H G P I pS S G N S P P M R
MH+ = 2615,15718
133
111
y_9-P_
b_5_
b_7_
b_8_
b_17-p_
b_9-p_
y_9-P_
b_15_
b_19-P_
S125
R D H Q S P V S V L H G P I pS S G N S P P M R
MH+ = 2535,19327
111
133
b_14_
y_9_
b_15-P_
b_19_

### Slide 6
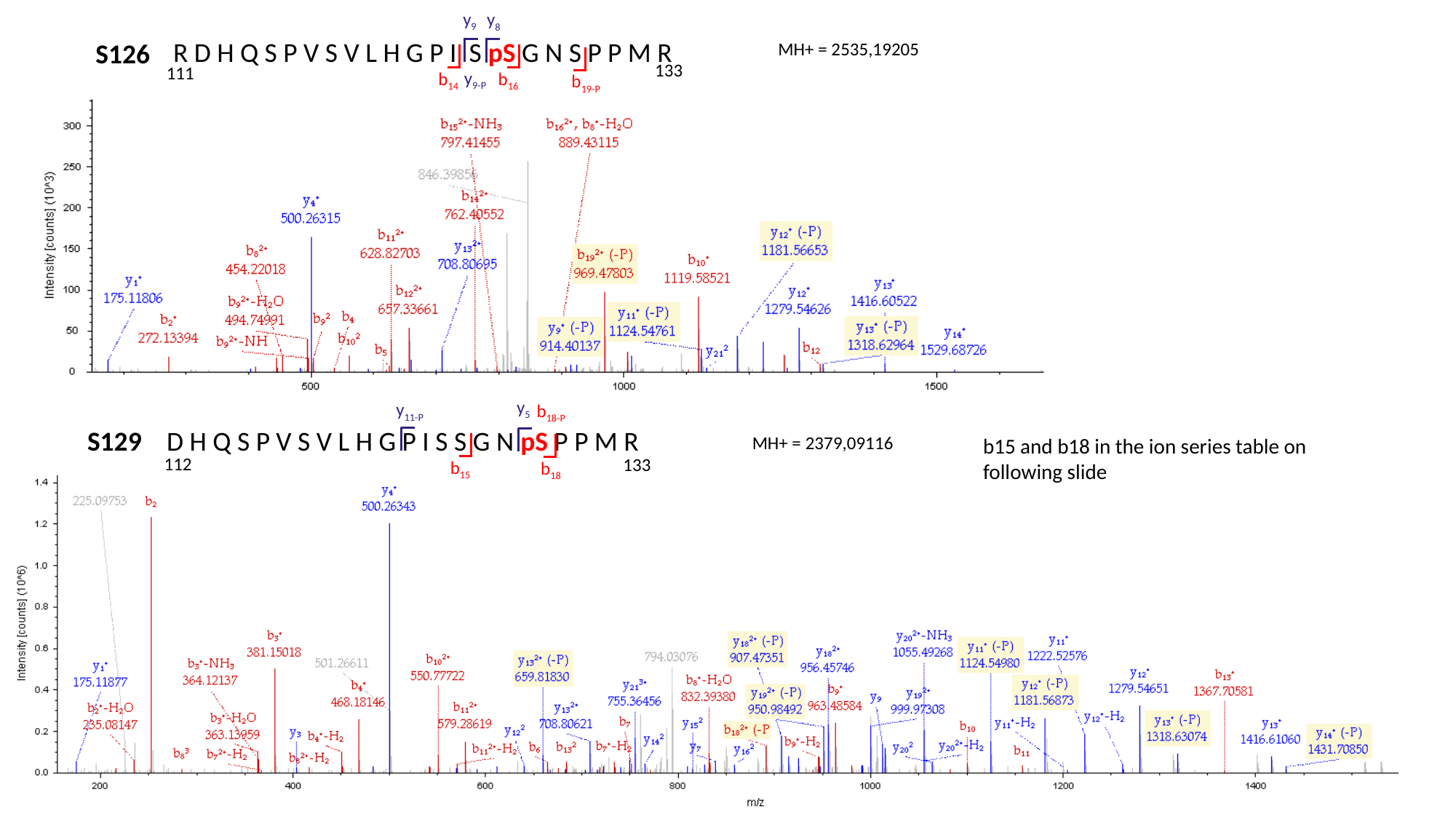

y_9_
y_8_
R D H Q S P V S V L H G P I S pS G N S P P M R
S126
MH+ = 2535,19205
133
111
y_9-P_
b_14_
b_16_
b_19-P_
y_5_
y_11-P_
b_18-P_
S129
D H Q S P V S V L H G P I S S G N pS P P M R
MH+ = 2379,09116
b15 and b18 in the ion series table on following slide
112
133
b_15_
b_18_

### Slide 7
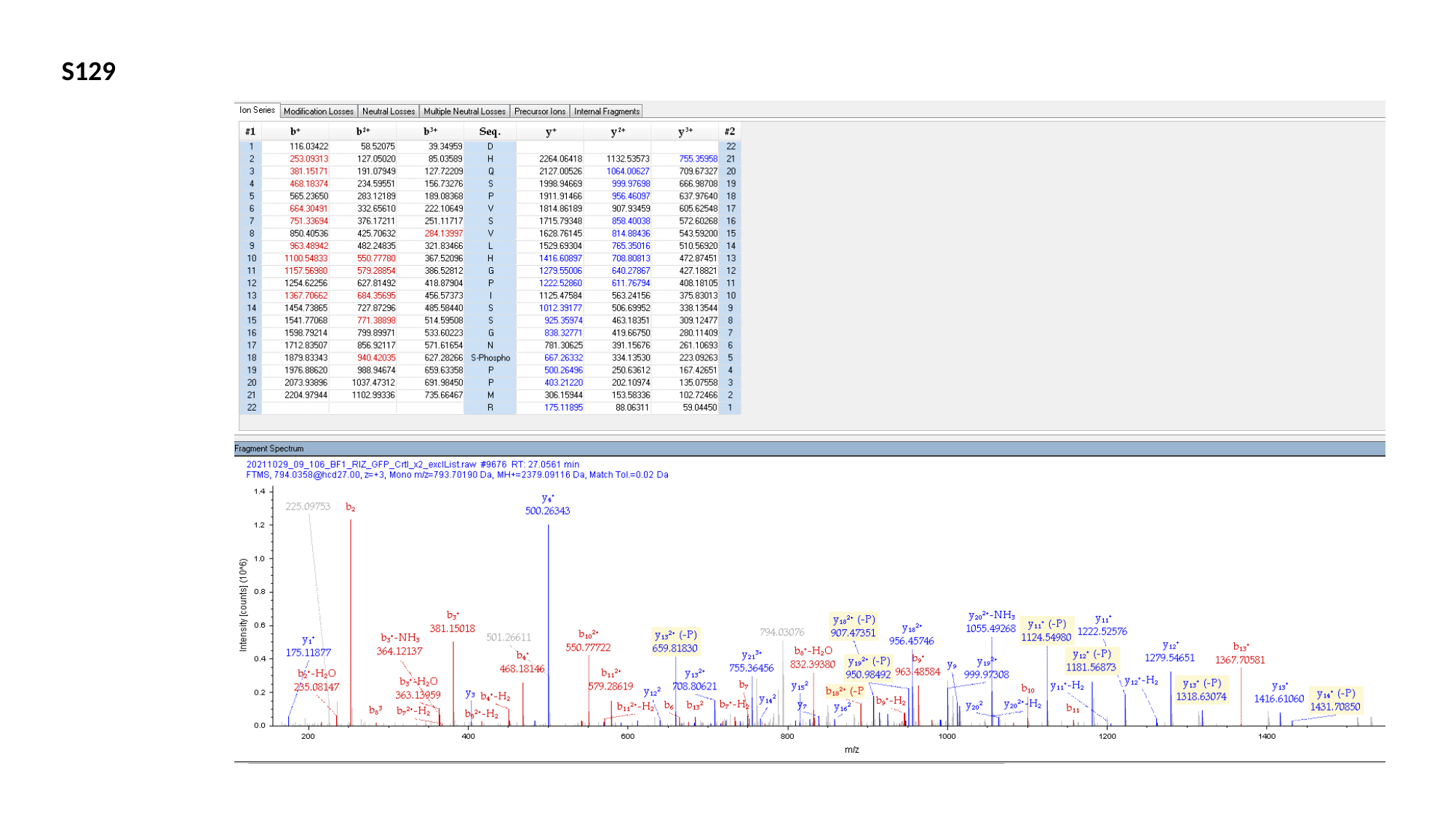

S129

### Slide 8
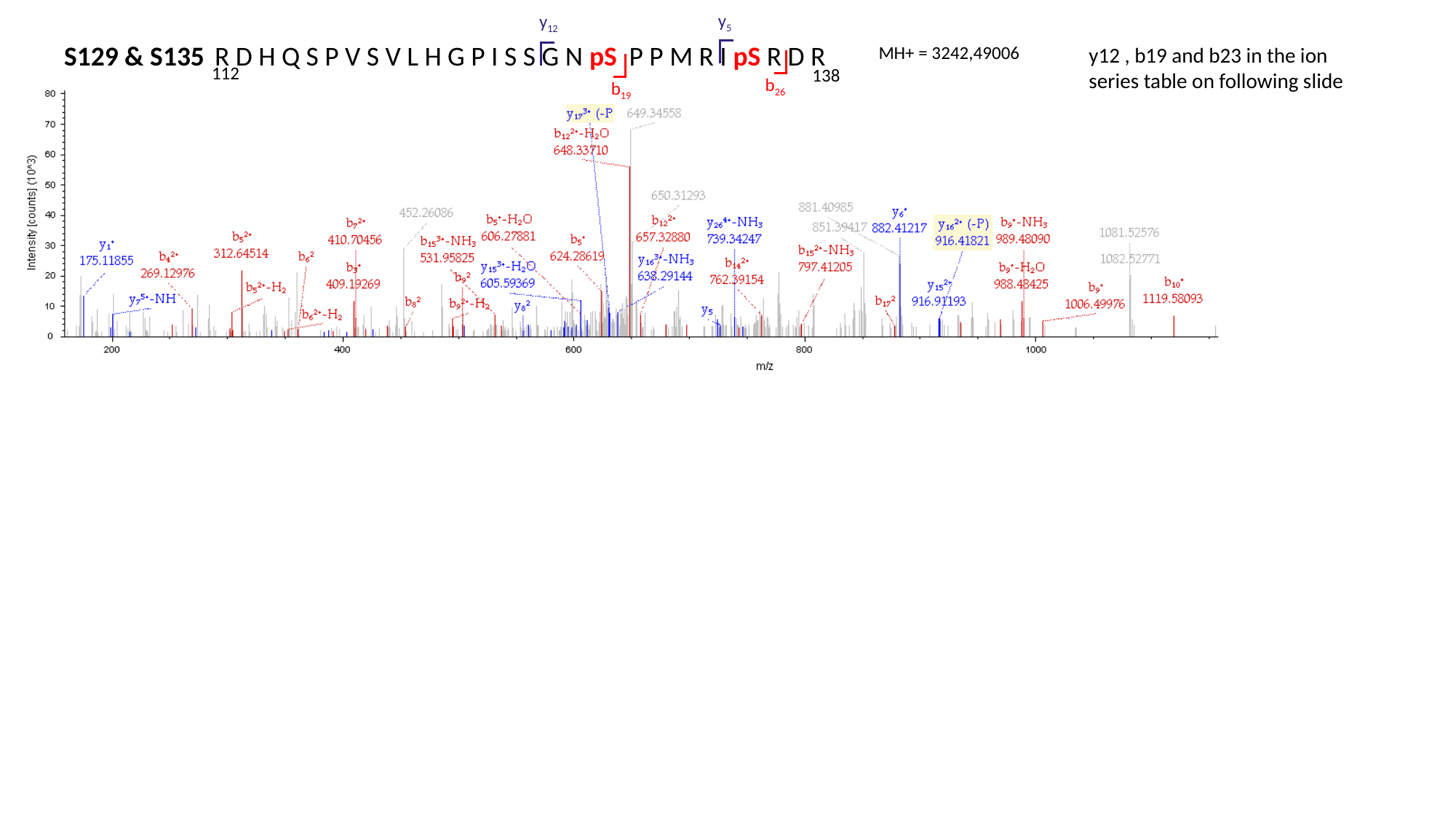

y_5_
y_12_
S129 & S135
R D H Q S P V S V L H G P I S S G N pS P P M R I pS R D R
MH+ = 3242,49006
y12 , b19 and b23 in the ion series table on following slide
112
138
b_26_
b_19_

### Slide 9
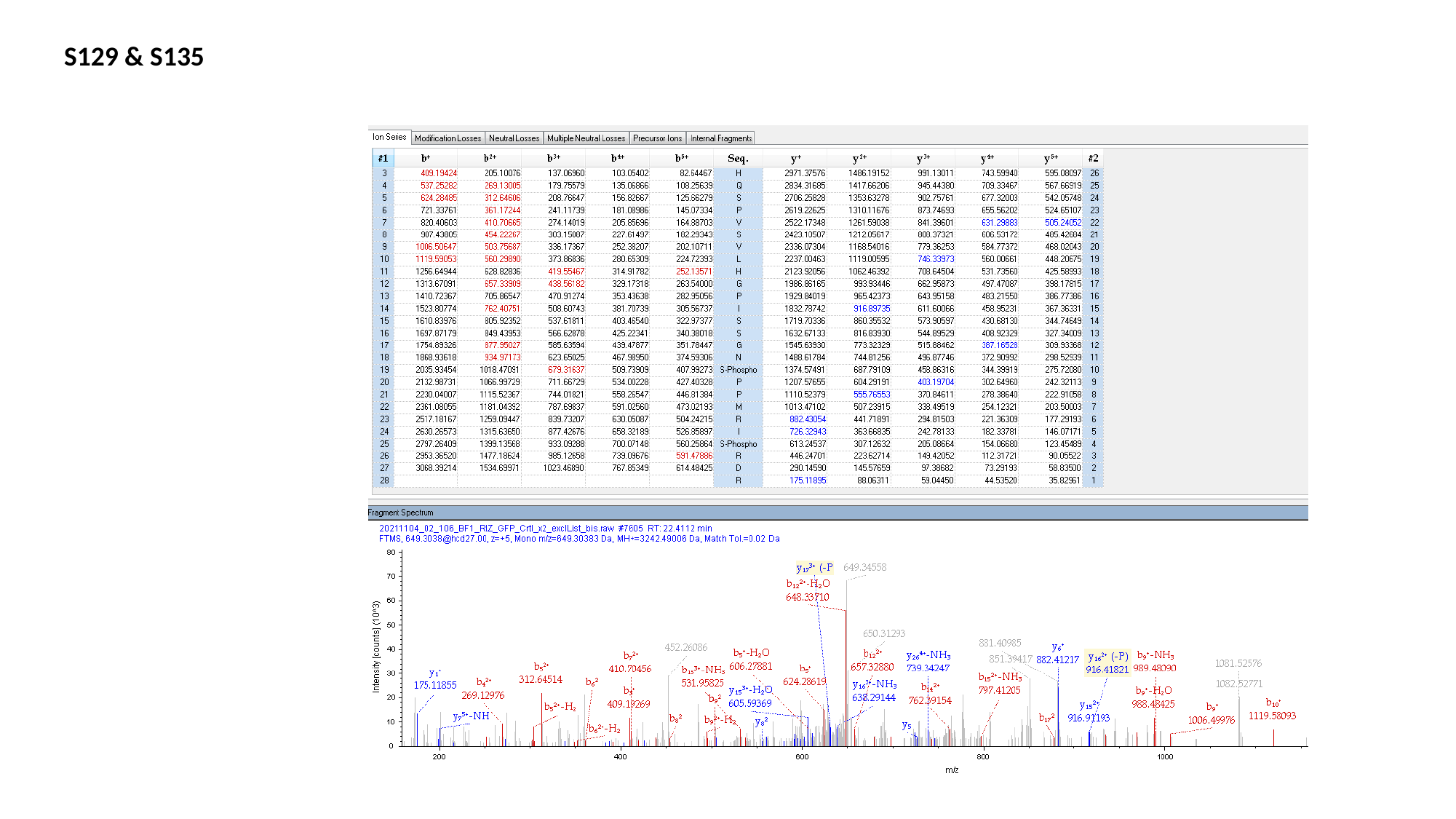

S129 & S135

### Slide 10
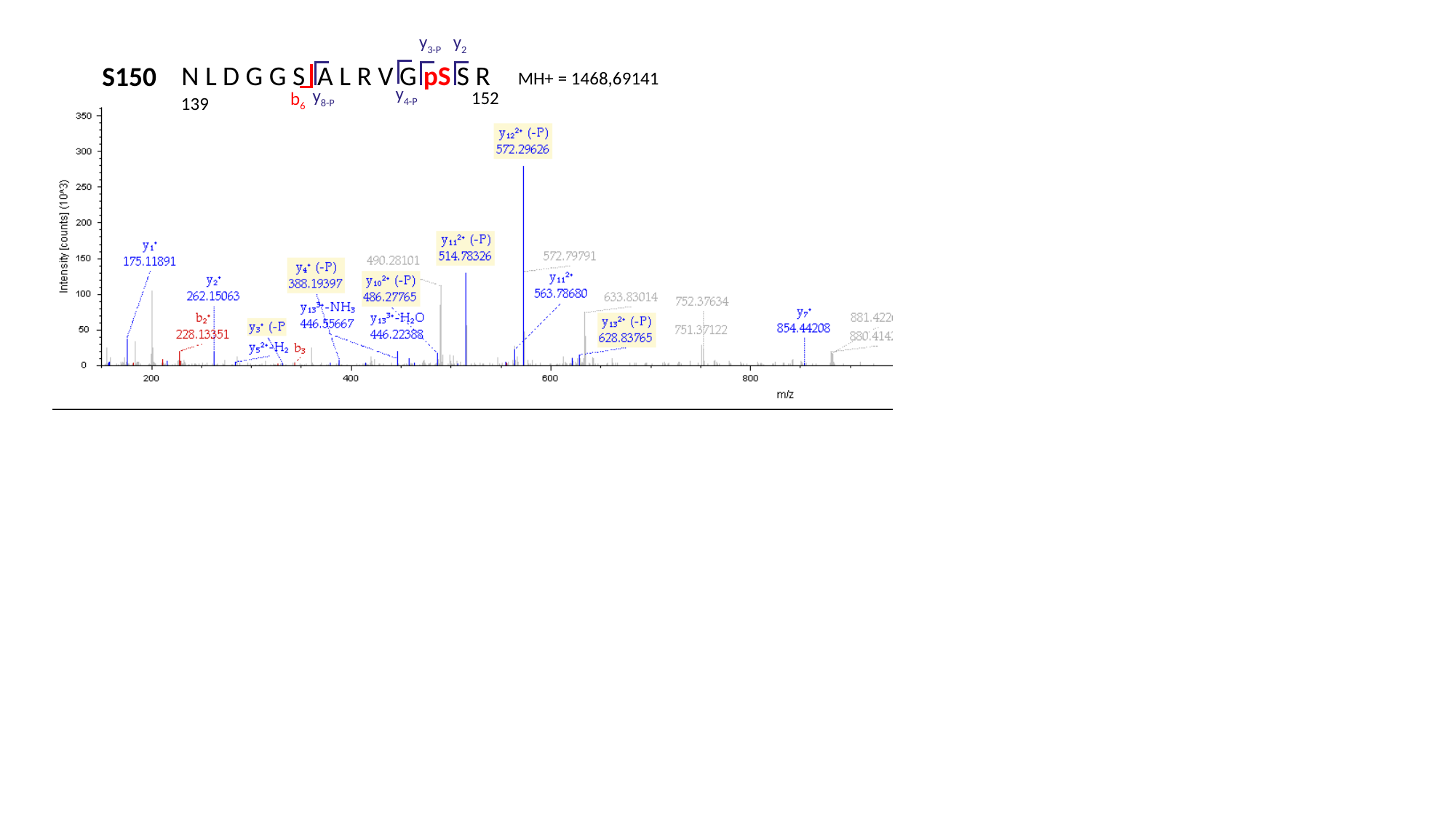

y_3-P_
y_2_
N L D G G S A L R V G pS S R
S150
MH+ = 1468,69141
y_4-P_
y_8-P_
152
b_6_
139
