## Supplementary Data Files for "Post-translational regulation of photosynthetic activity via the TOR kinase in plants": Fig1H.pptx

### Slide 1
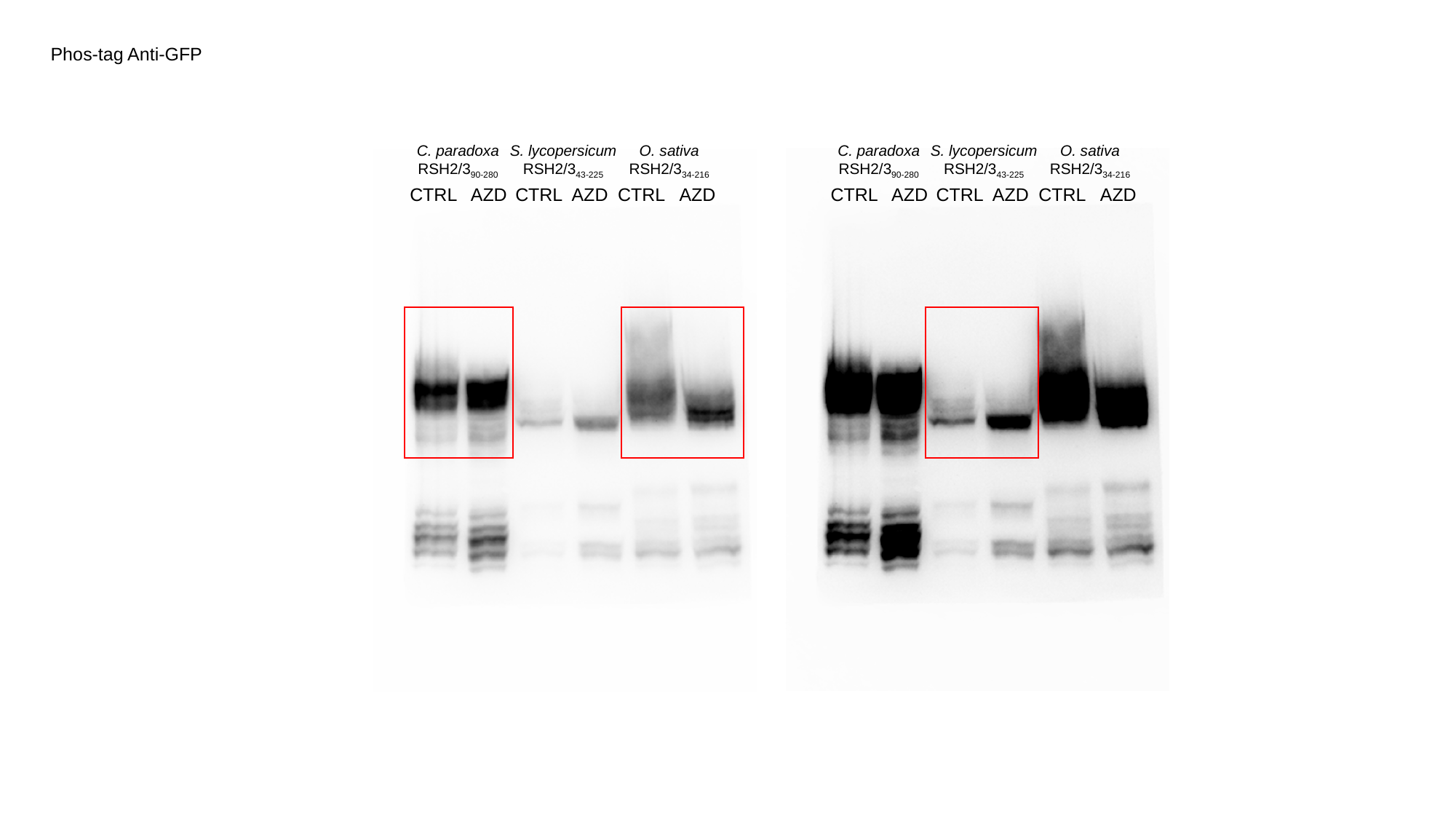

Phos-tag Anti-GFP
C. paradoxa
RSH2/3_90-280_
S. lycopersicum
RSH2/3_43-225_
O. sativa
RSH2/3_34-216_
C. paradoxa
RSH2/3_90-280_
S. lycopersicum
RSH2/3_43-225_
O. sativa
RSH2/3_34-216_
CTRL
AZD
CTRL
AZD
CTRL
AZD
CTRL
AZD
CTRL
AZD
CTRL
AZD

### Slide 2
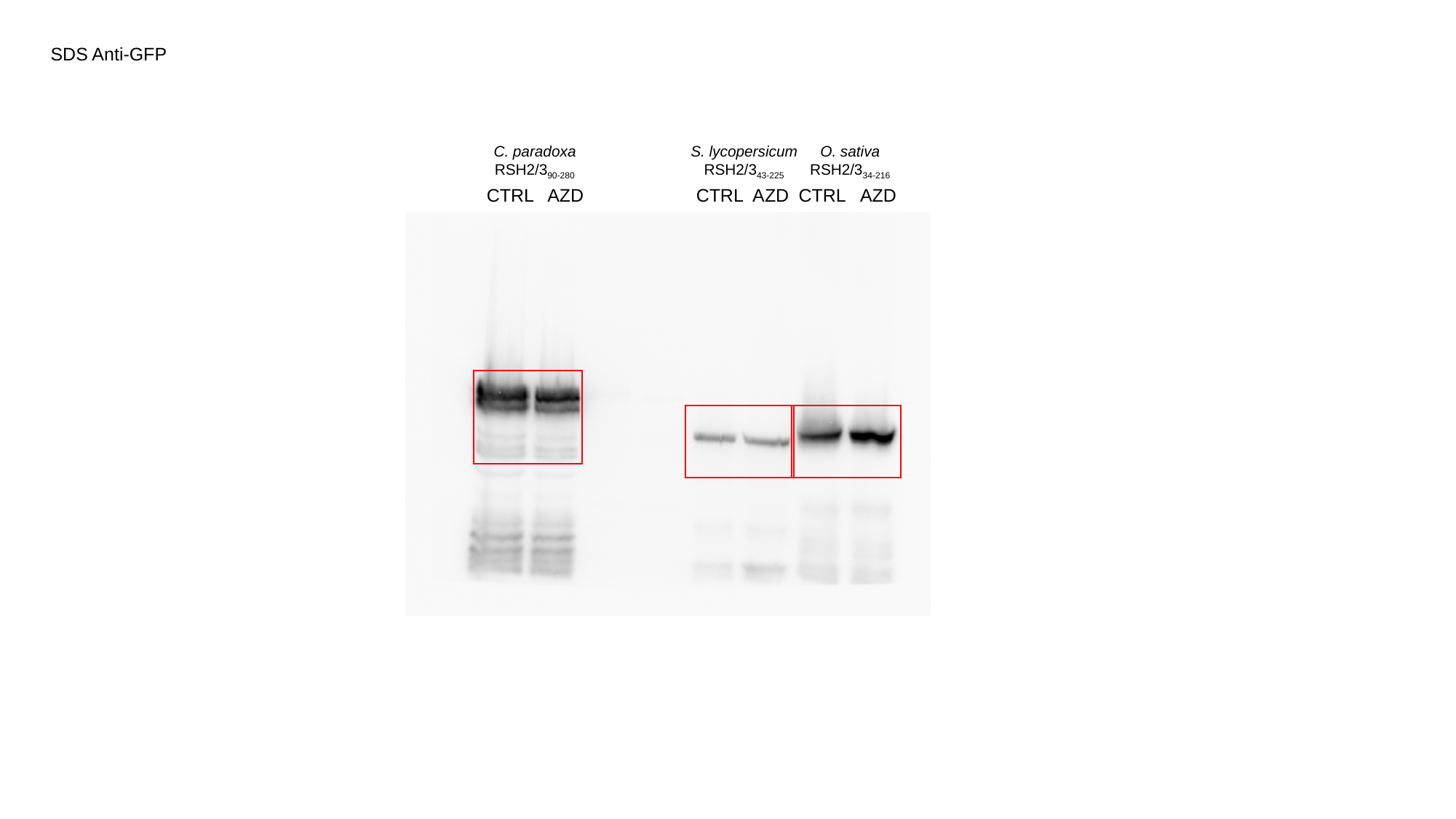

SDS Anti-GFP
C. paradoxa
RSH2/3_90-280_
S. lycopersicum
RSH2/3_43-225_
O. sativa
RSH2/3_34-216_
CTRL
AZD
CTRL
AZD
CTRL
AZD
