## Supplementary Data Files for "Post-translational regulation of photosynthetic activity via the TOR kinase in plants": FigS3B.pptx

### Slide 1
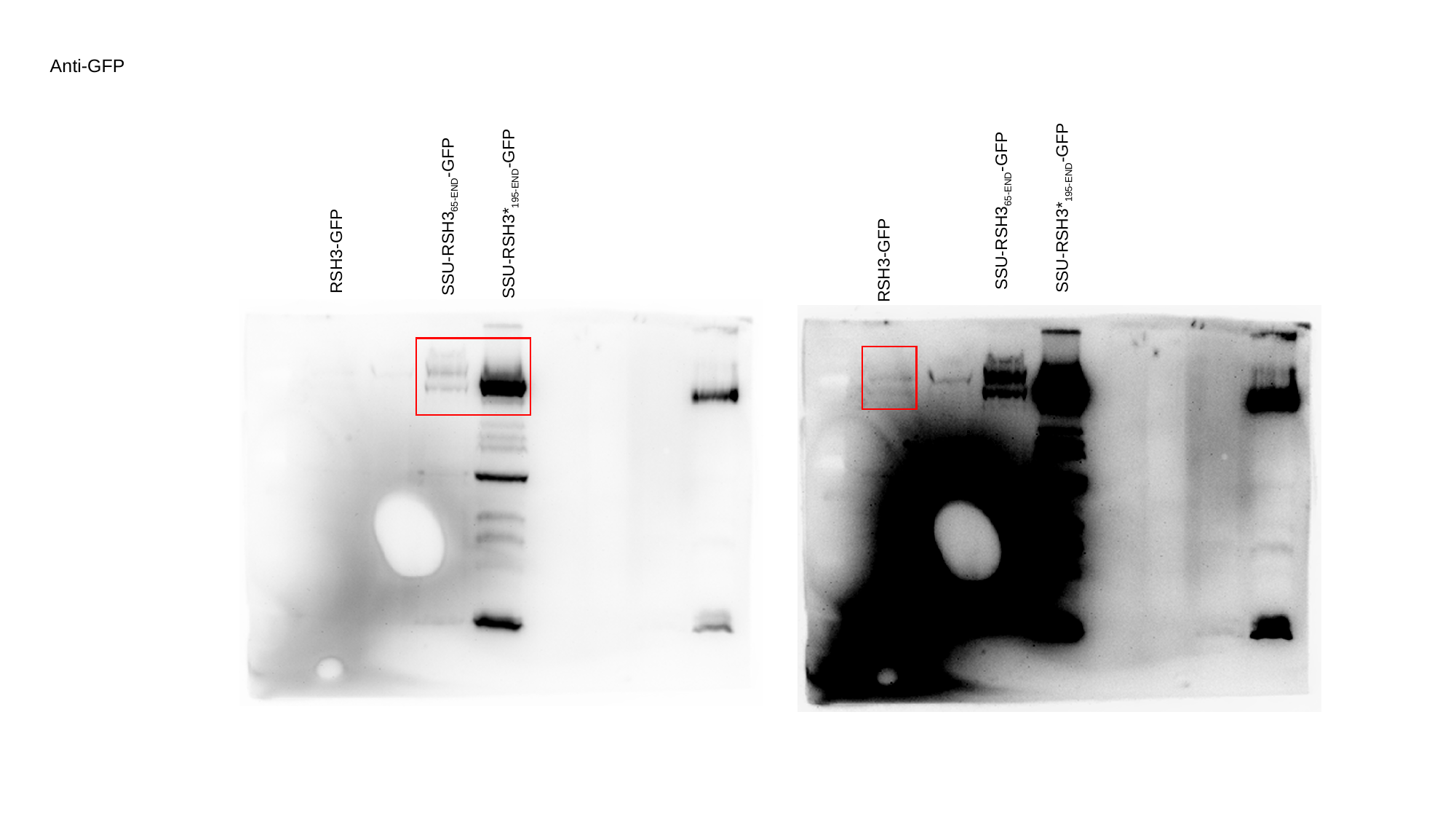

Anti-GFP
SSU-RSH3*_195-END_-GFP
SSU-RSH3_65-END_-GFP
SSU-RSH3*_195-END_-GFP
SSU-RSH3_65-END_-GFP
RSH3-GFP
RSH3-GFP

### Slide 2
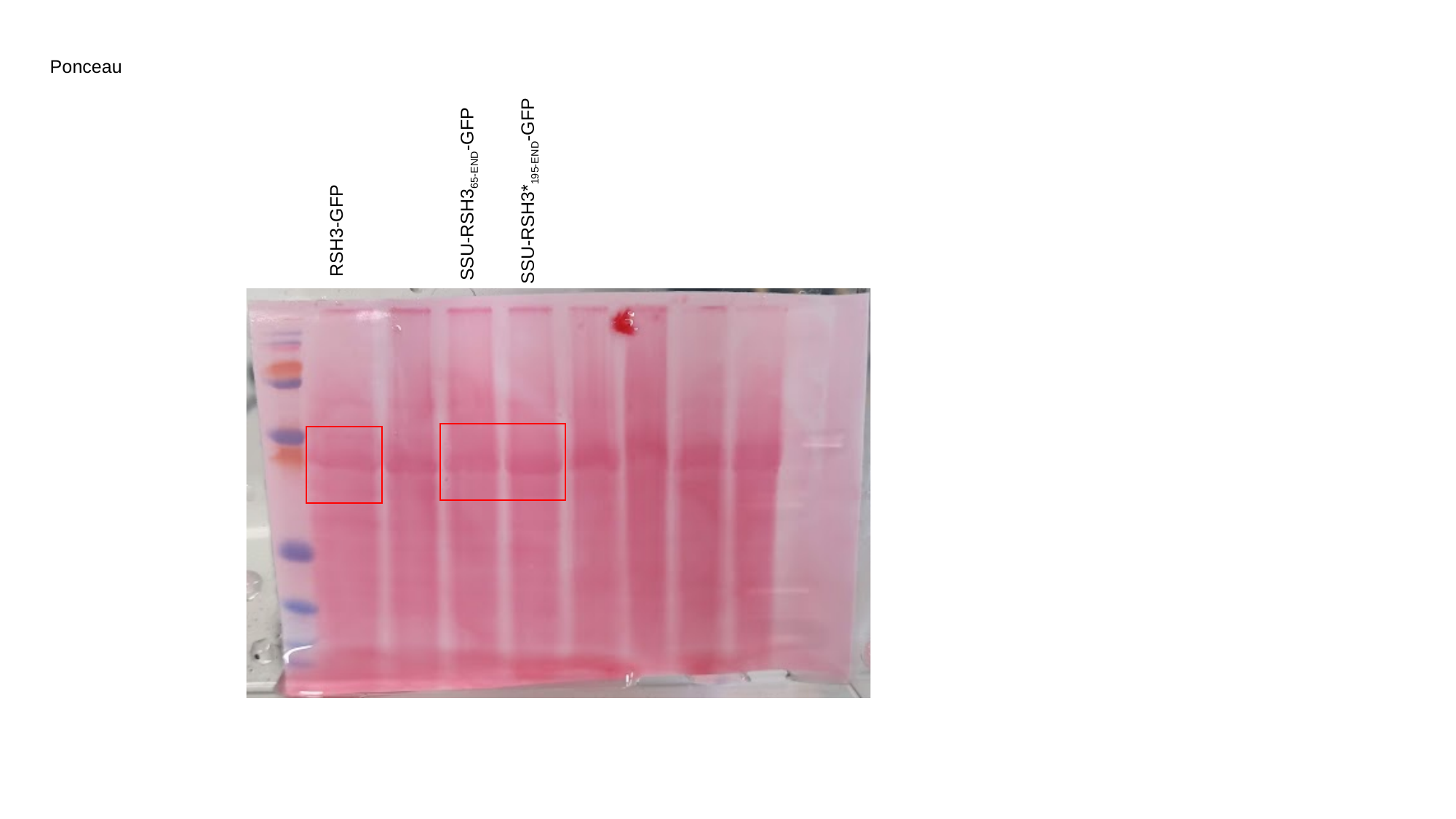

Ponceau
SSU-RSH3*_195-END_-GFP
SSU-RSH3_65-END_-GFP
RSH3-GFP
