## Supplementary Data Files for "Post-translational regulation of photosynthetic activity via the TOR kinase in plants": FigS4A.pptx

### Slide 1
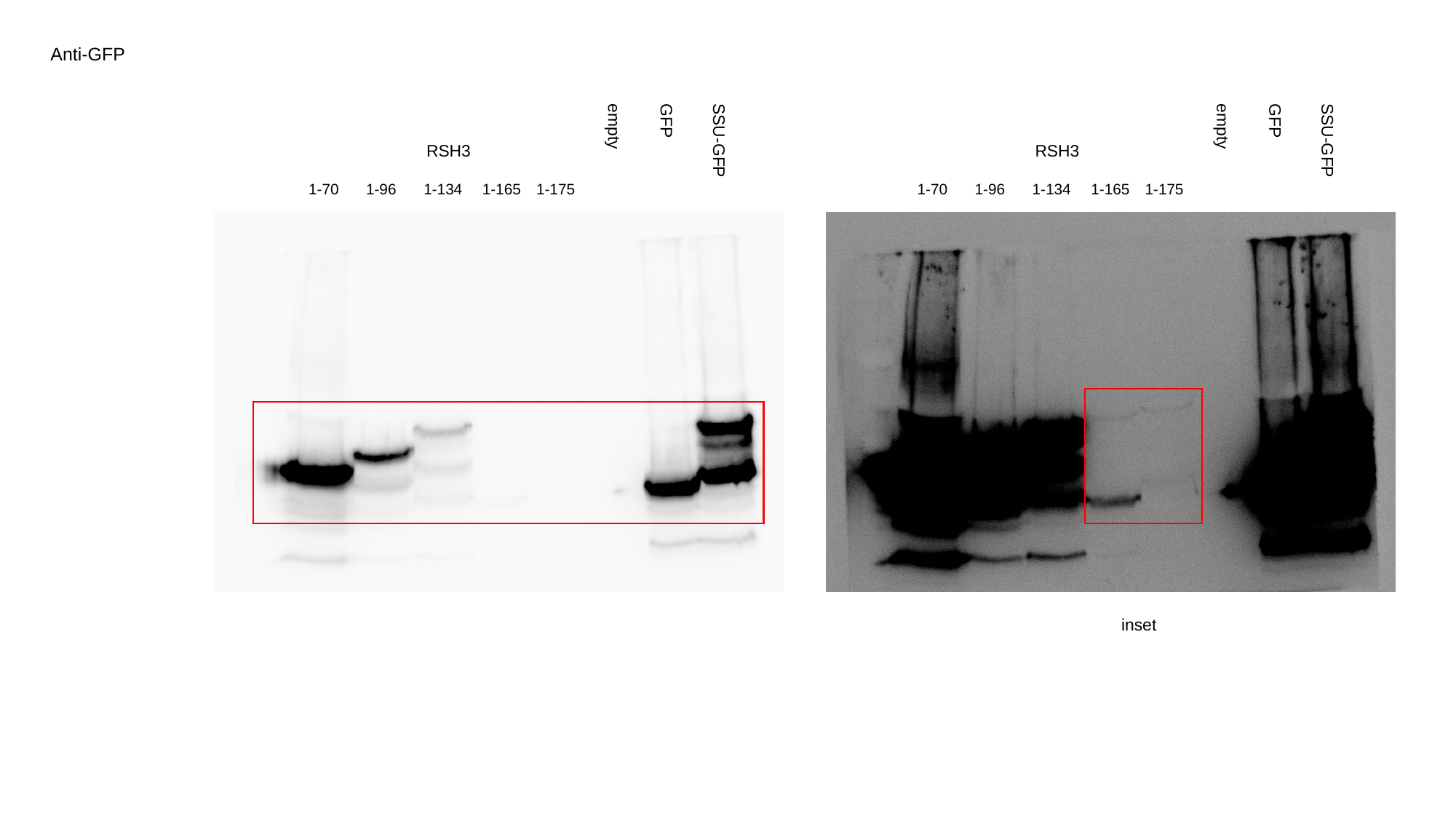

Anti-GFP
GFP
empty
RSH3
SSU-GFP
1-70
1-96
1-134
1-165
1-175
GFP
empty
RSH3
SSU-GFP
1-70
1-96
1-134
1-165
1-175
inset

### Slide 2
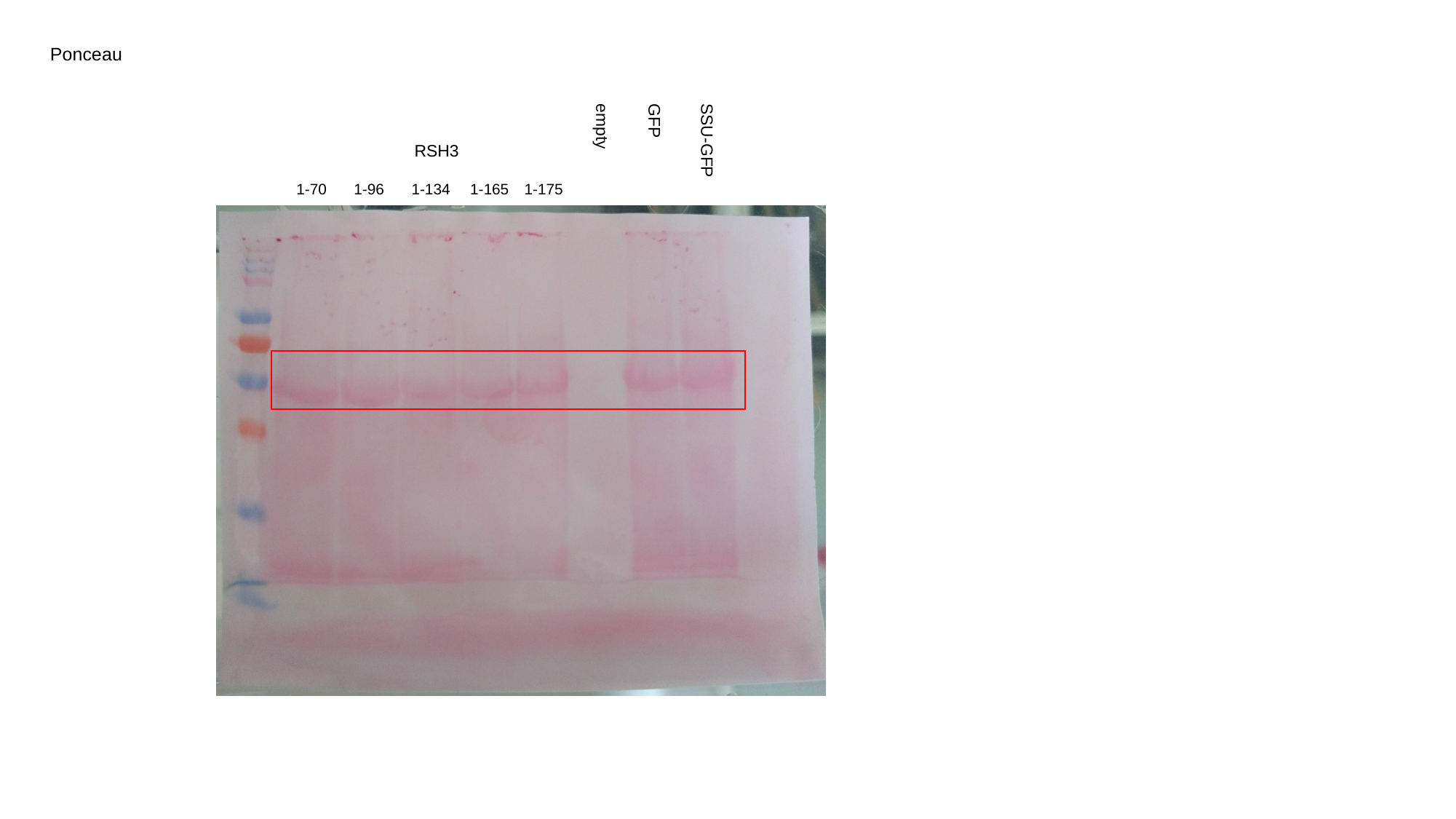

Ponceau
GFP
empty
RSH3
SSU-GFP
1-70
1-96
1-134
1-165
1-175
