## Supplementary Data Files for "Post-translational regulation of photosynthetic activity via the TOR kinase in plants": Stats_and_Plot.ipynb

Jupyter Notebook


Jupyter Notebook requires JavaScript.  
Please enable it to proceed.

Logout

Menu

Kernel

 


- File
  - New NotebookDropdown
  - Open...
  - Make a Copy...
  - Save as...
  - Rename...
  - Save and Checkpoint
  - Revert to CheckpointDropdown
  - Print Preview
  - Download asDropdown
    - AsciiDoc (.asciidoc)
    - HTML (.html)
    - HTML + ToC (.html)
    - LaTeX (.tex)
    - Markdown (.md)
    - Notebook (.ipynb)
    - PDF via LaTeX (.pdf)
    - PDF via HTML (.html)
    - PNG via HTML (.html)
    - reST (.rst)
    - Script ()
    - Reveal.js slides (.slides.html)
    - PDF via HTML (.html)
  - Deploy as
  - Trust Notebook
  - Close and Halt
- Edit
  - Cut Cells
  - Copy Cells
  - Paste Cells Above
  - Paste Cells Below
  - Paste Cells & Replace
  - Delete Cells
  - Undo Delete Cells
  - Split Cell
  - Merge Cell Above
  - Merge Cell Below
  - Move Cell Up
  - Move Cell Down
  - Edit Notebook Metadata
  - Find and Replace
  - Cut Cell Attachments
  - Copy Cell Attachments
  - Paste Cell Attachments
  - Insert Image
- View
  - Toggle Header
  - Toggle Toolbar
  - Toggle Line Numbers
  - Cell Toolbar
- Insert
  - Insert Cell Above
  - Insert Cell Below
- Cell
  - Run Cells
  - Run Cells and Select Below
  - Run Cells and Insert Below
  - Run All
  - Run All Above
  - Run All Below
  - Cell Type
    - Code
    - Markdown
    - Raw NBConvert
  - Current Outputs
    - Toggle
    - Toggle Scrolling
    - Clear
  - All Output
    - Toggle
    - Toggle Scrolling
    - Clear
- Kernel
  - Interrupt
  - Restart
  - Restart & Clear Output
  - Restart & Run All
  - Reconnect
  - Shutdown
  - Change kernel
- Help
  - User Interface Tour
  - Keyboard Shortcuts
  - Edit Keyboard Shortcuts
  - Notebook Help
  - Markdown
  - About
