## Supplementary figures and images for "Post-translational regulation of photosynthetic activity via the TOR kinase in plants"

### 3D.pptx

#### Slide 1
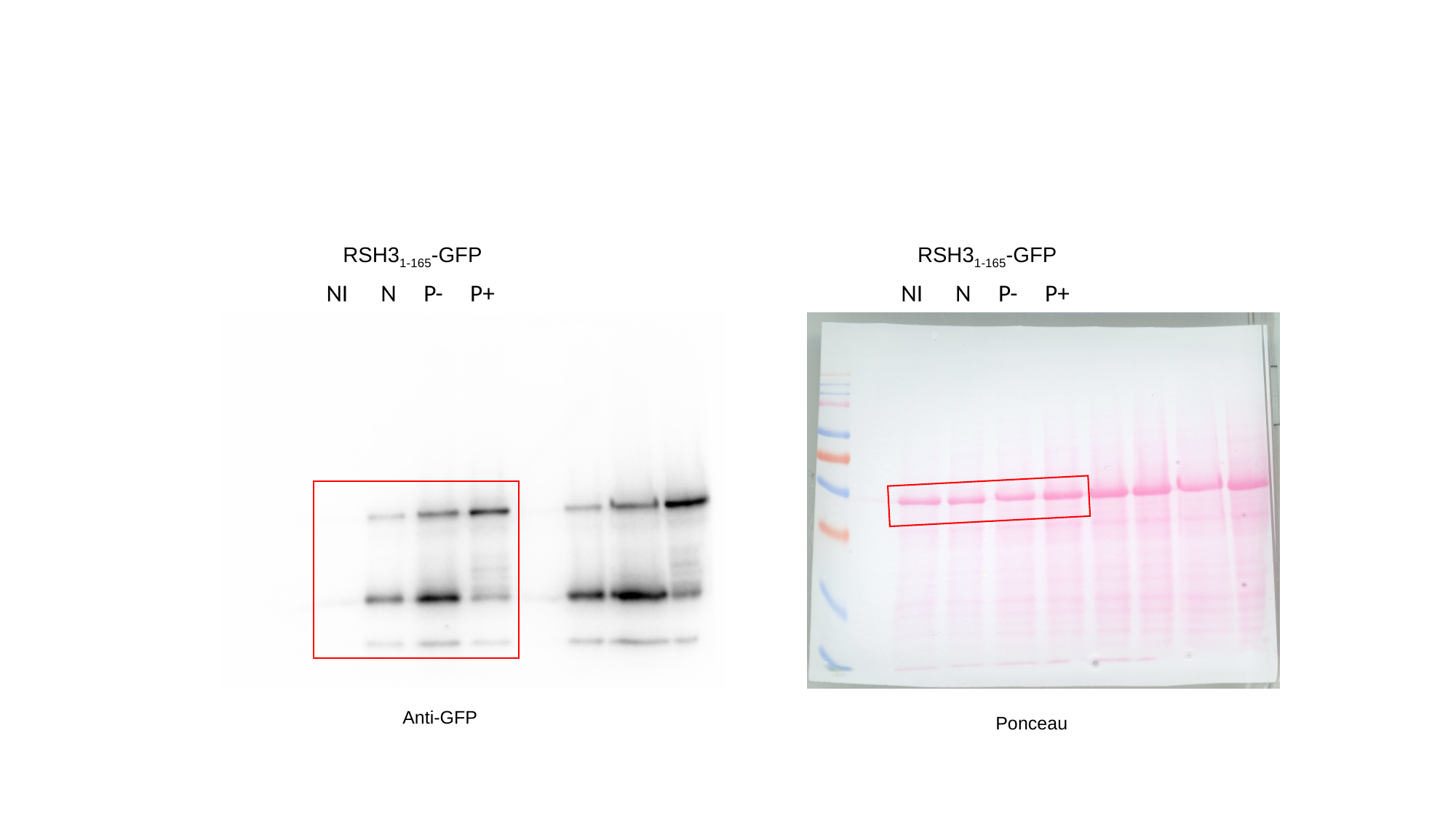

RSH3_1-165_-GFP
RSH3_1-165_-GFP
NI N P- P+
NI N P- P+
Anti-GFP
Ponceau

### anti-GFP blot.Tif

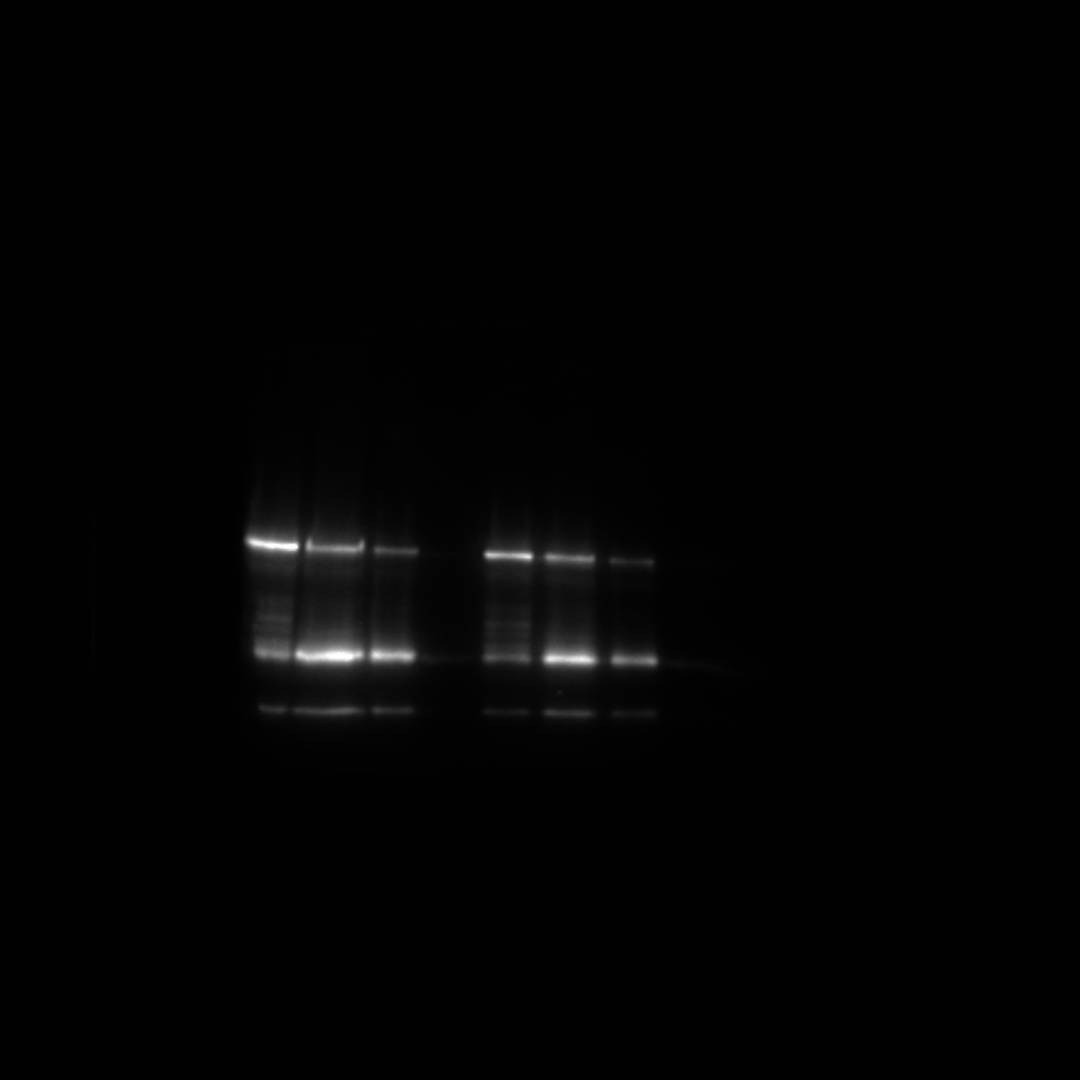

### anti-GFP.Tif

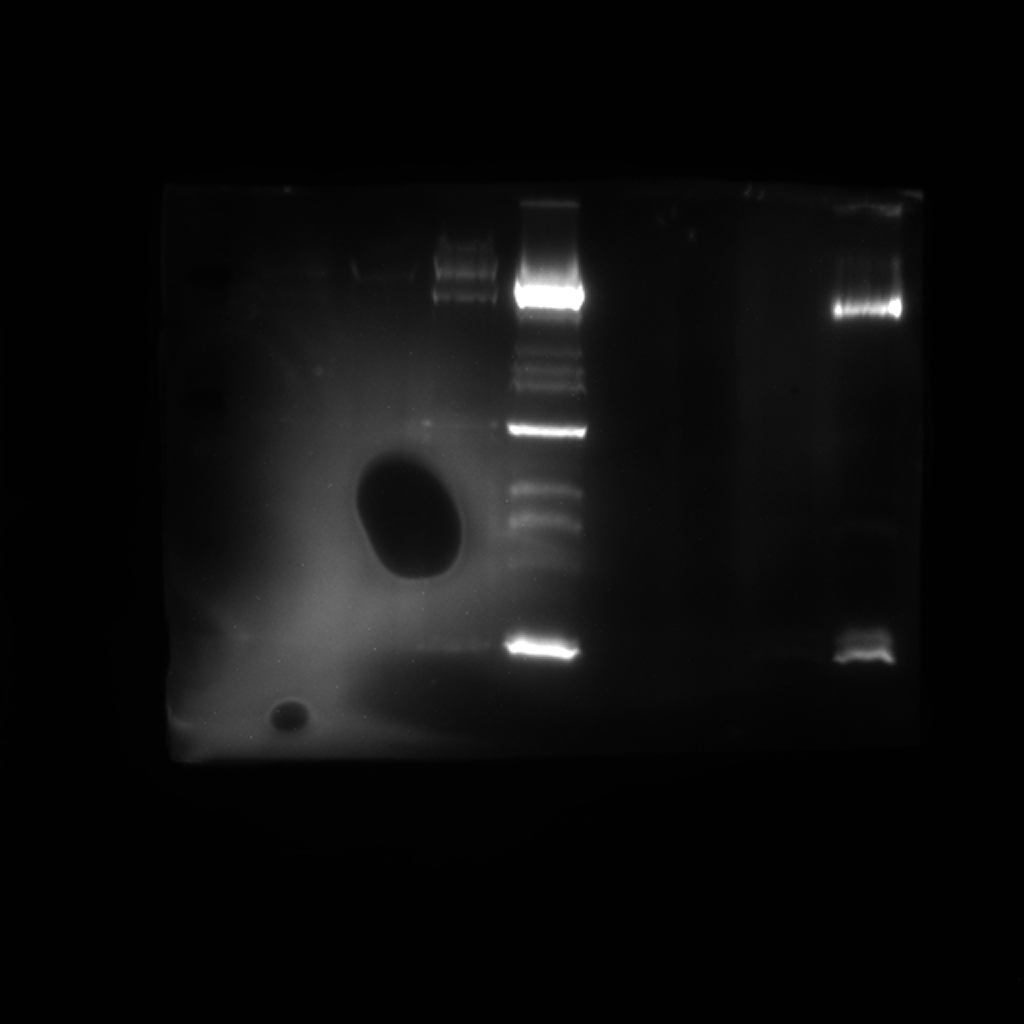

### anti-GFP.Tif

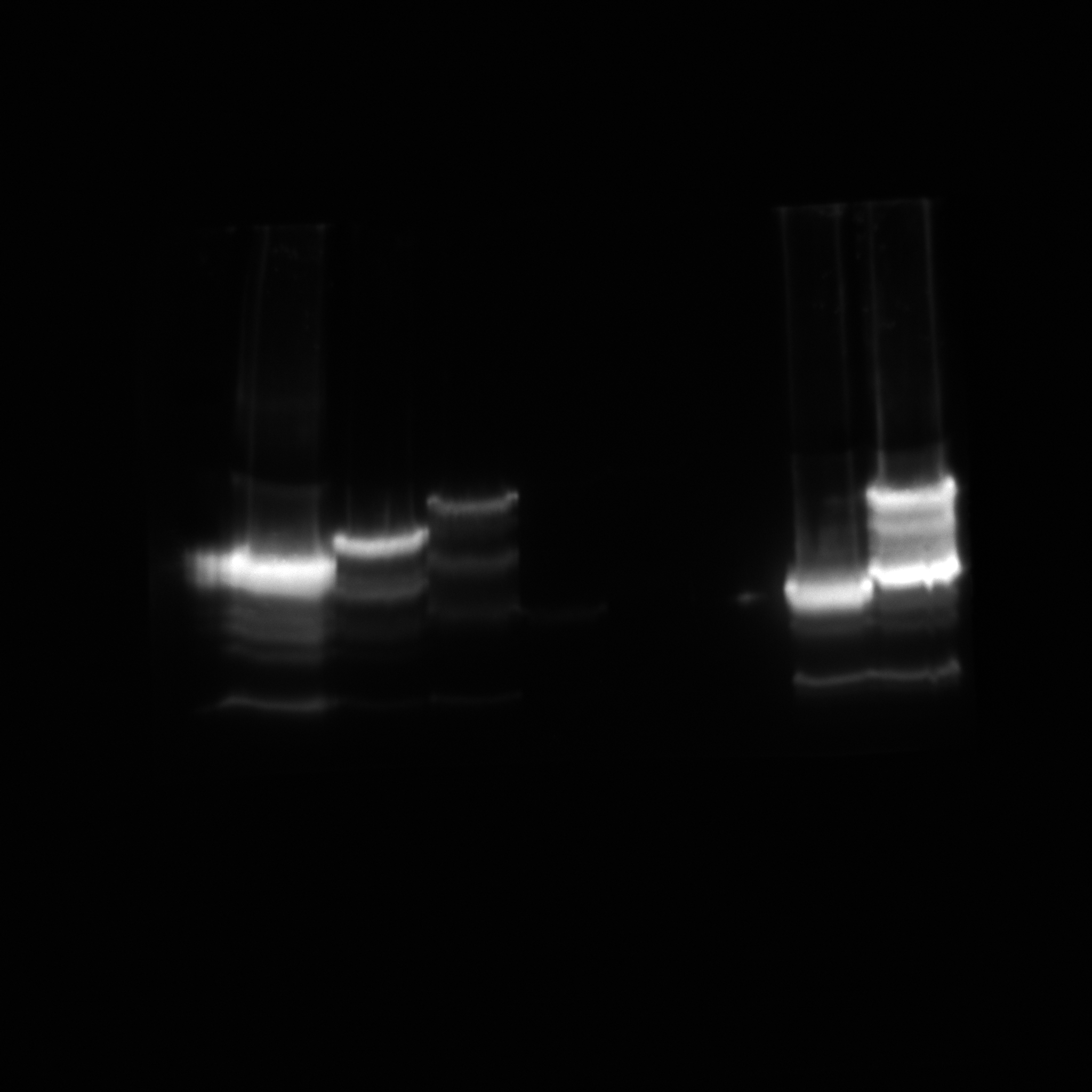

### biotin.Tif

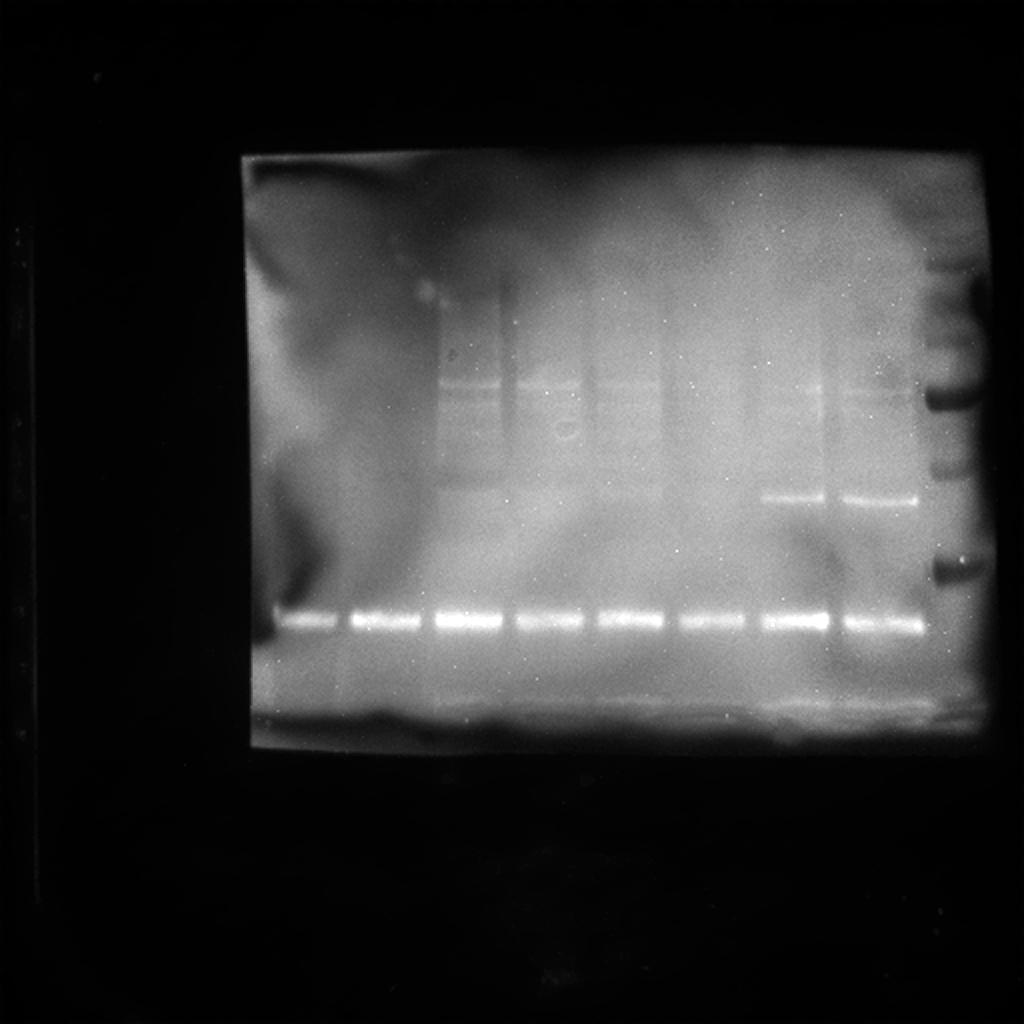

### epi.Tif

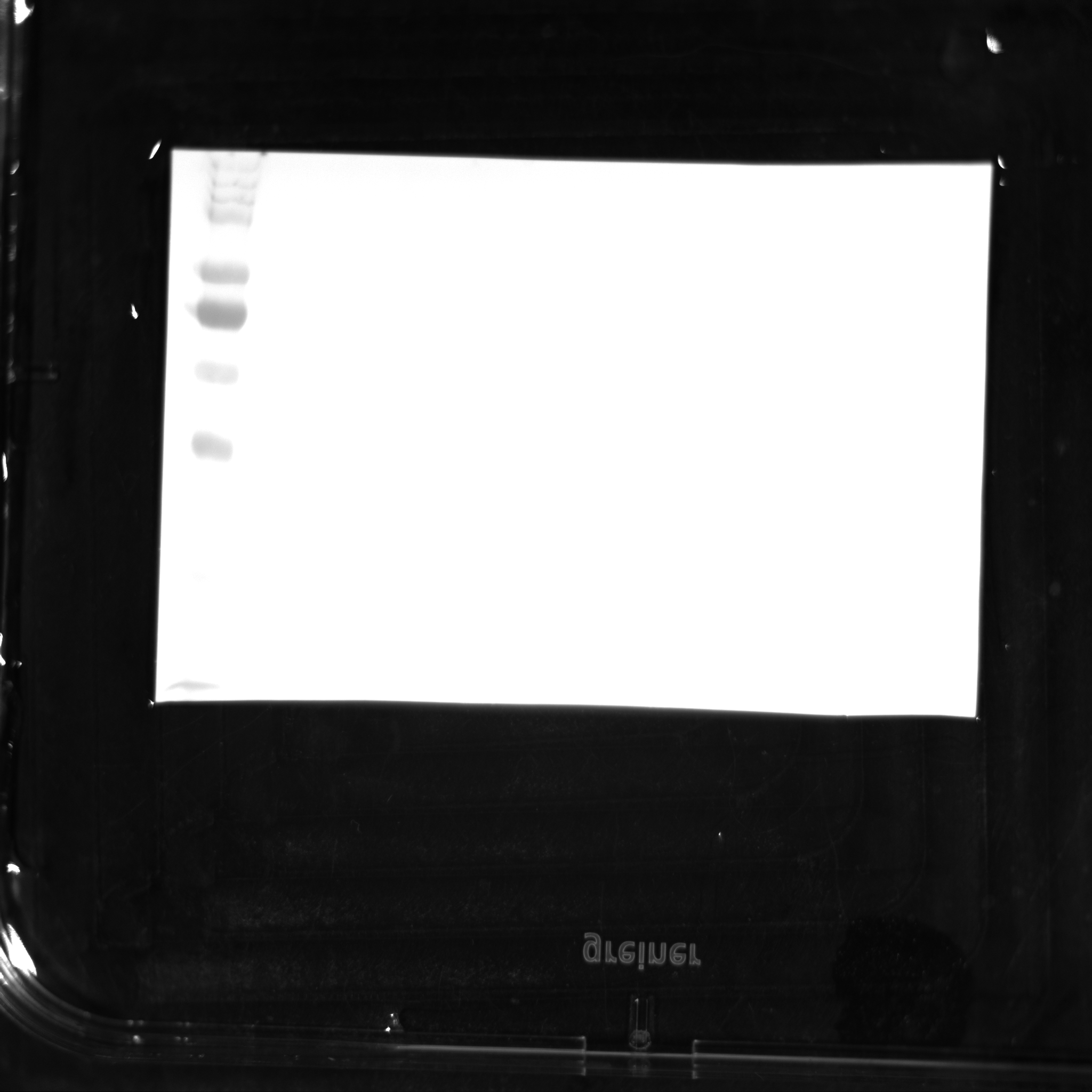

### Fig1E.pptx

#### Slide 1
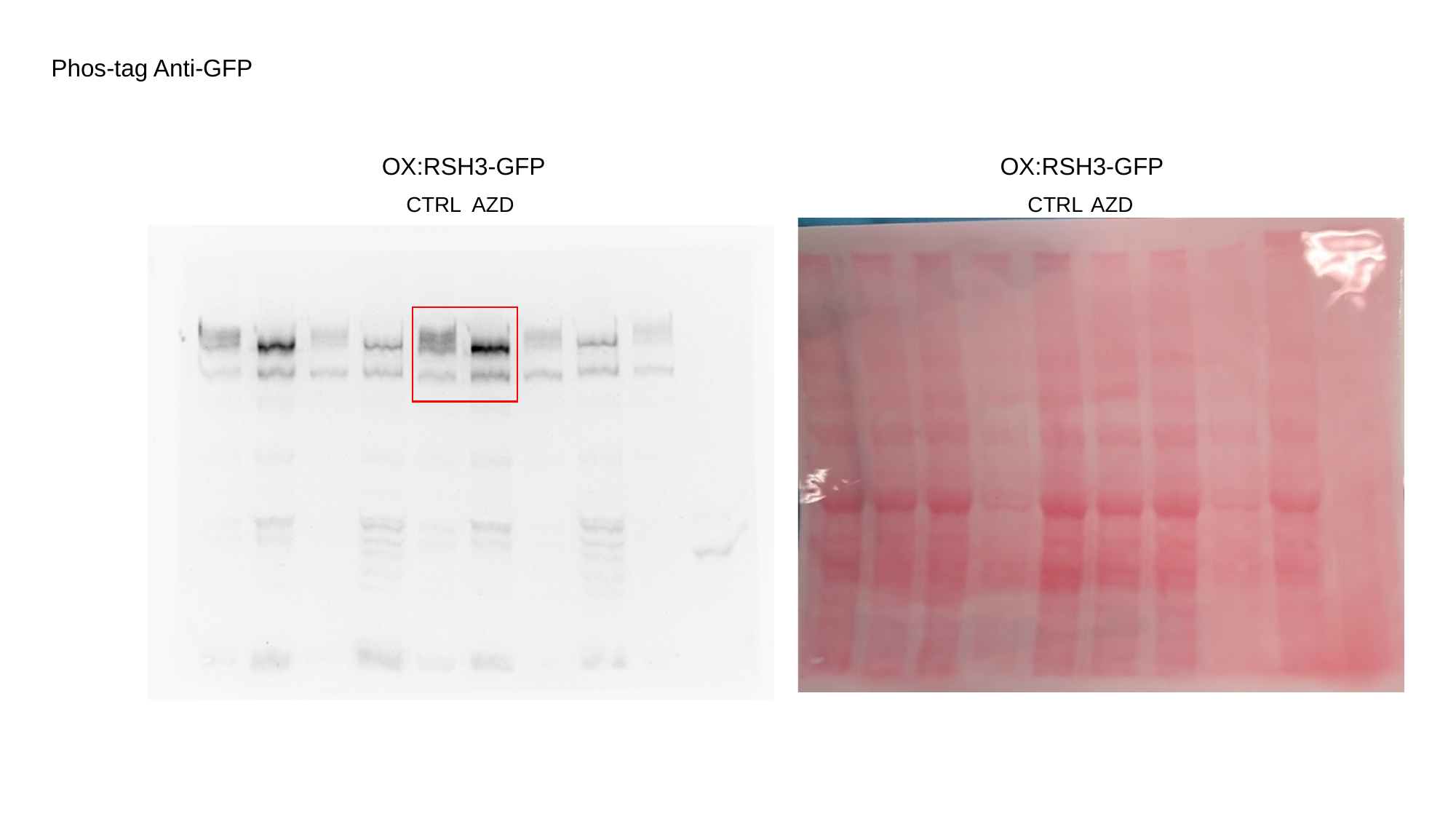

Phos-tag Anti-GFP
OX:RSH3-GFP
OX:RSH3-GFP
CTRL
AZD
CTRL
AZD

#### Slide 2
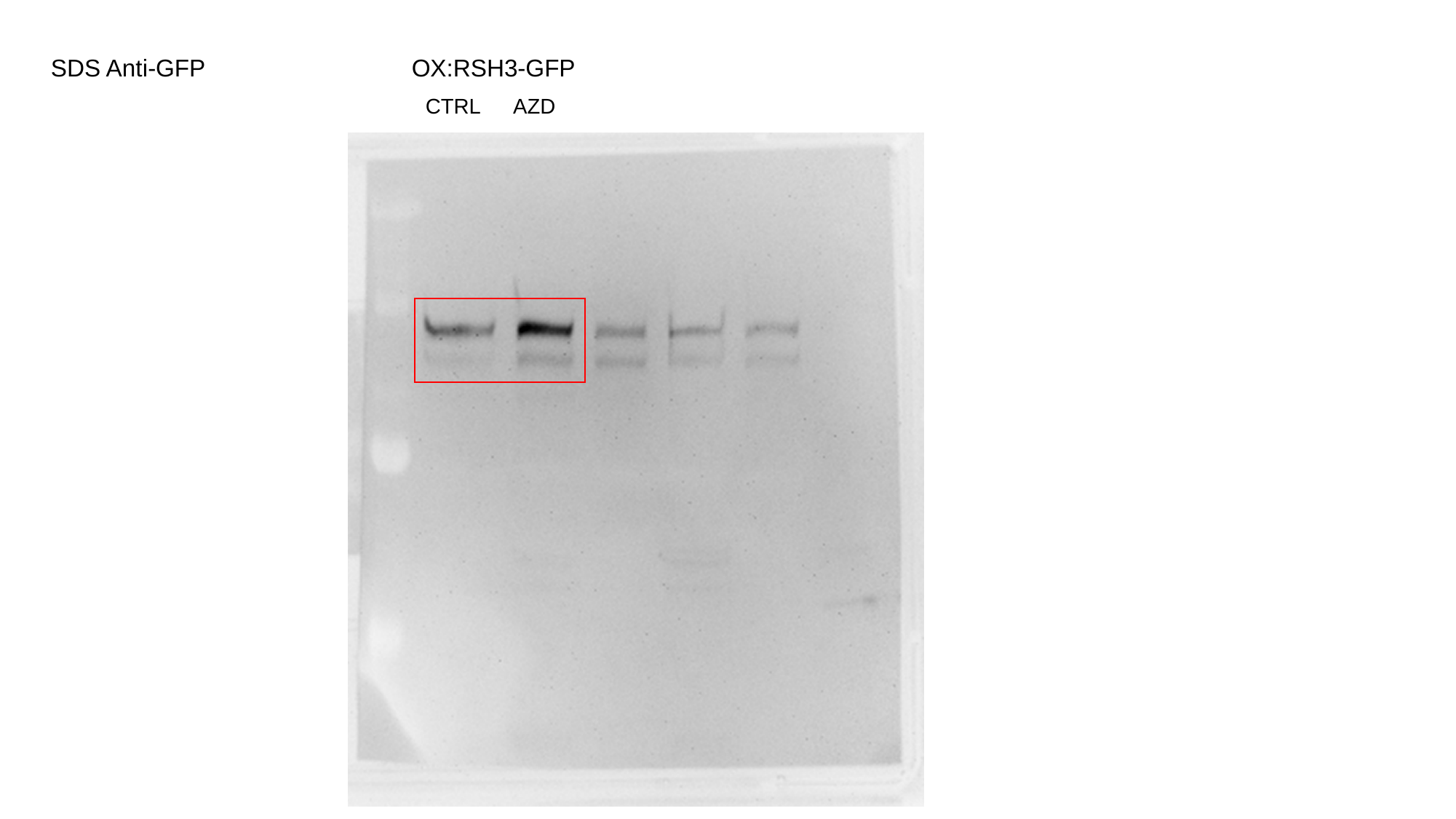

SDS Anti-GFP
OX:RSH3-GFP
CTRL
AZD

### GFP.Tif

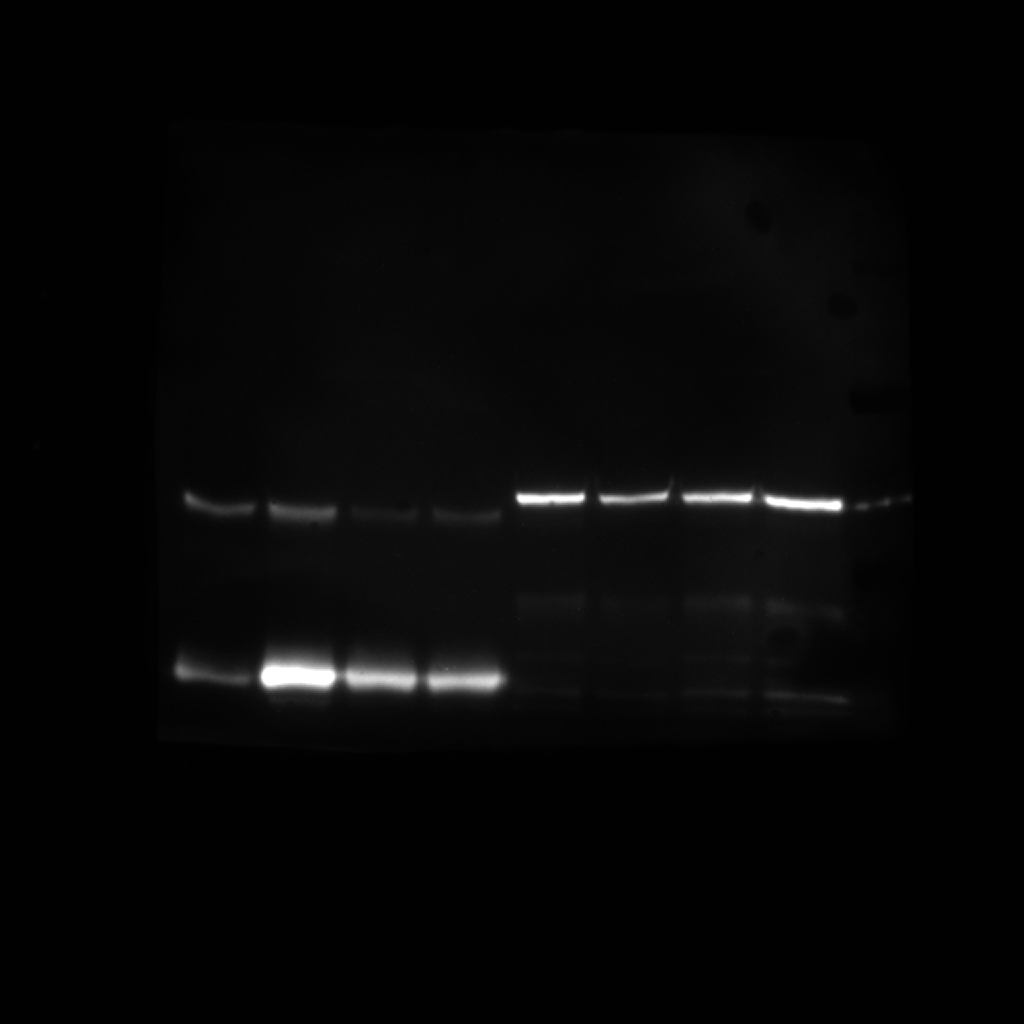

### IM000005 30.Tif

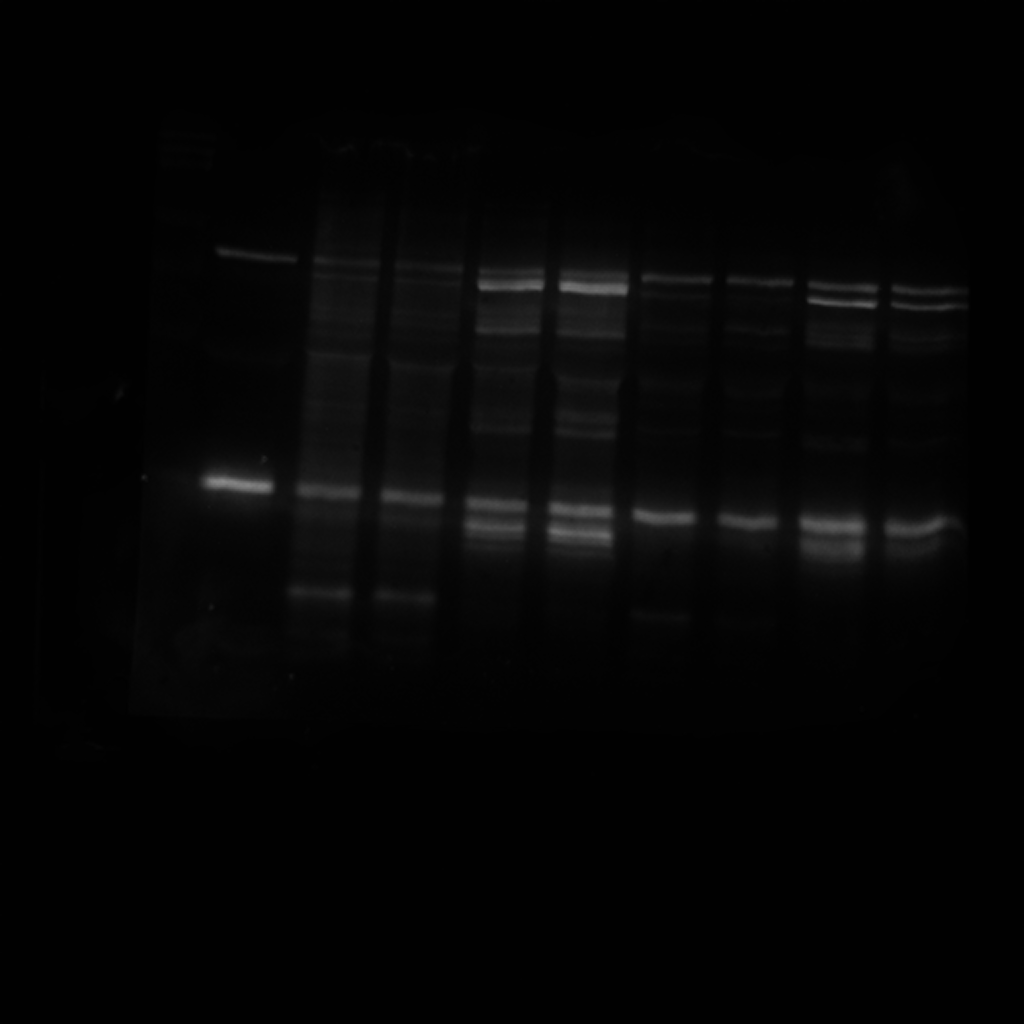

### IM000007 2min.Tif

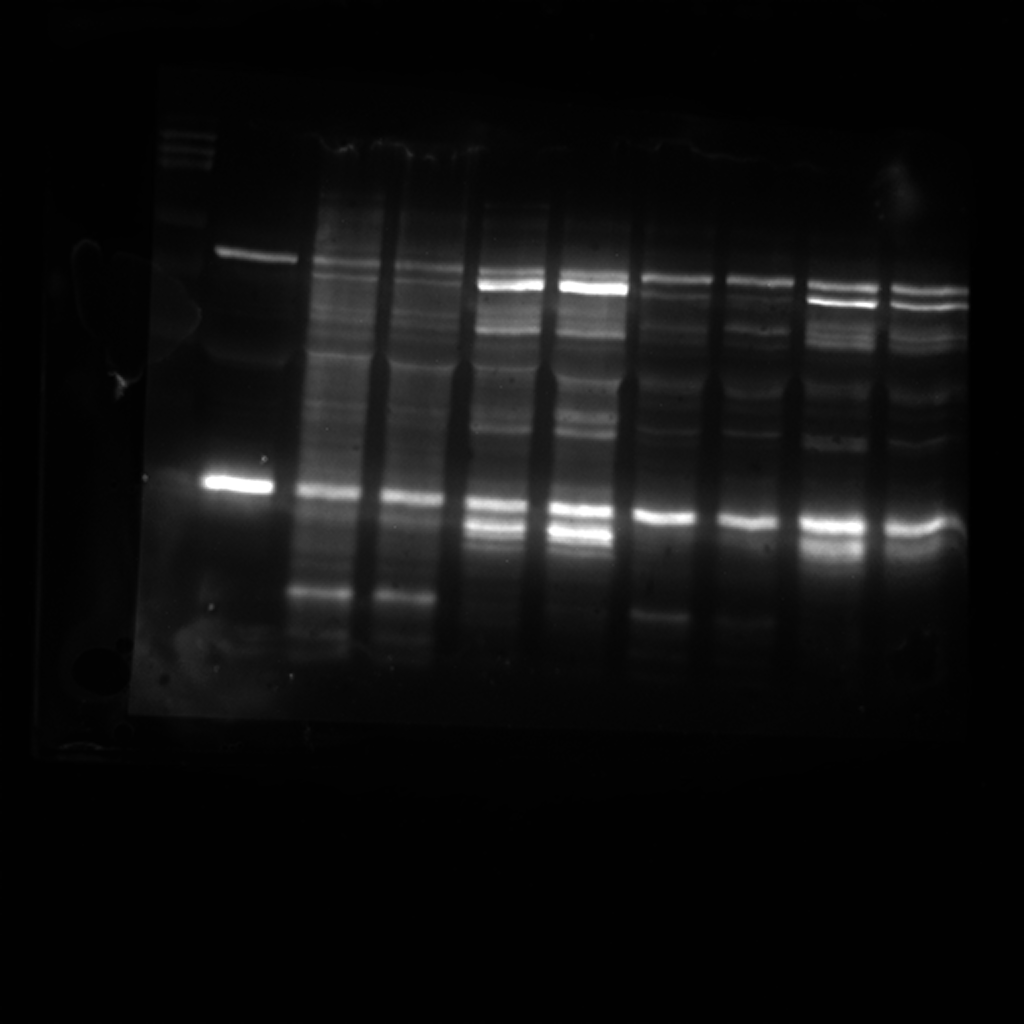

### IM000008 full.Tif

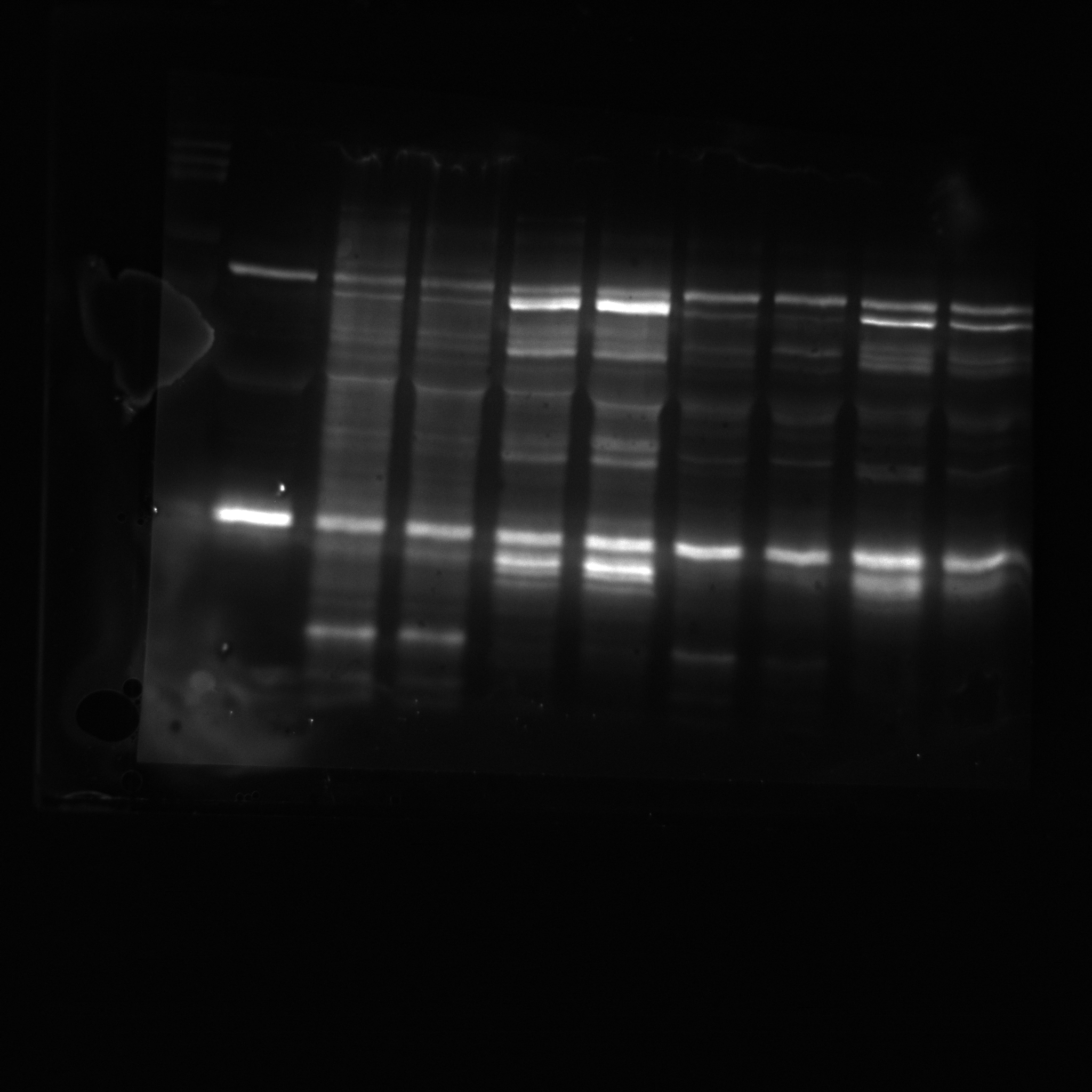

### IM000009 30.Tif

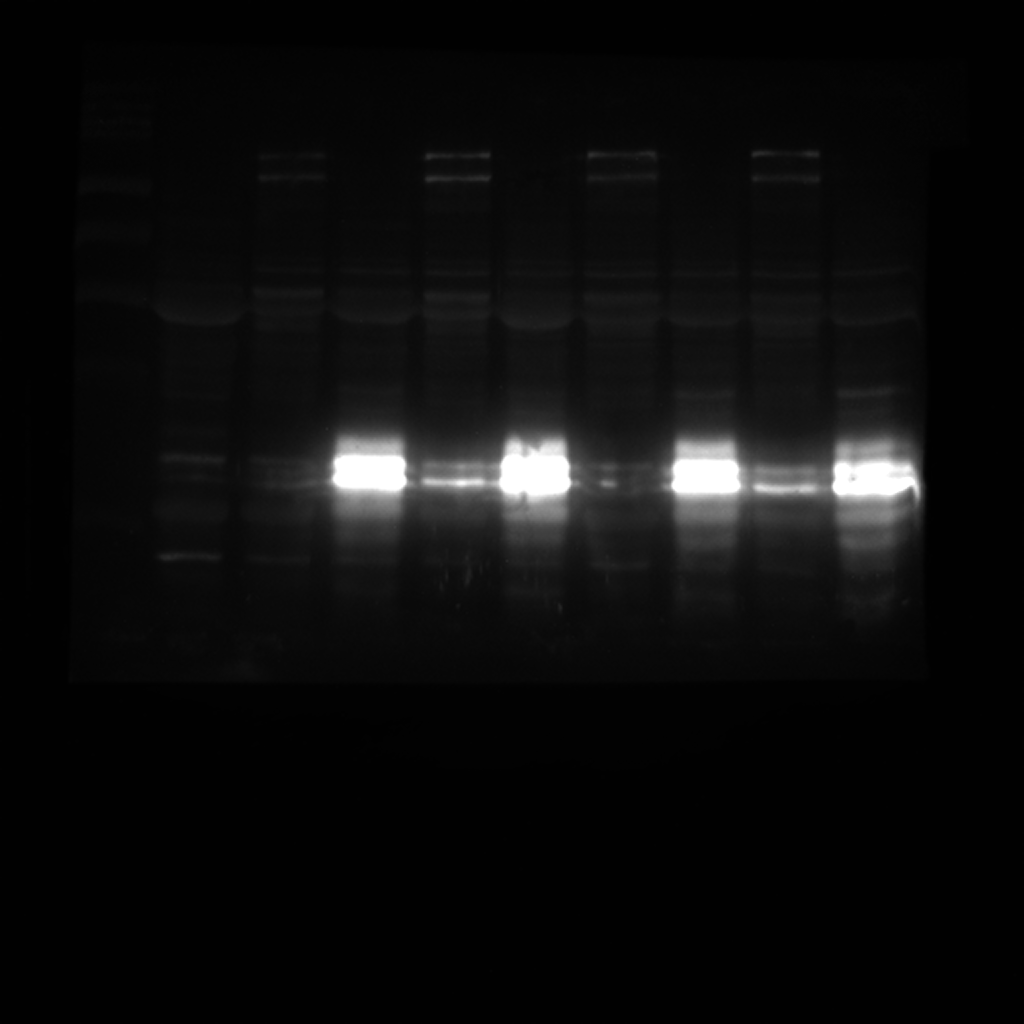

### IM000010 1min.Tif

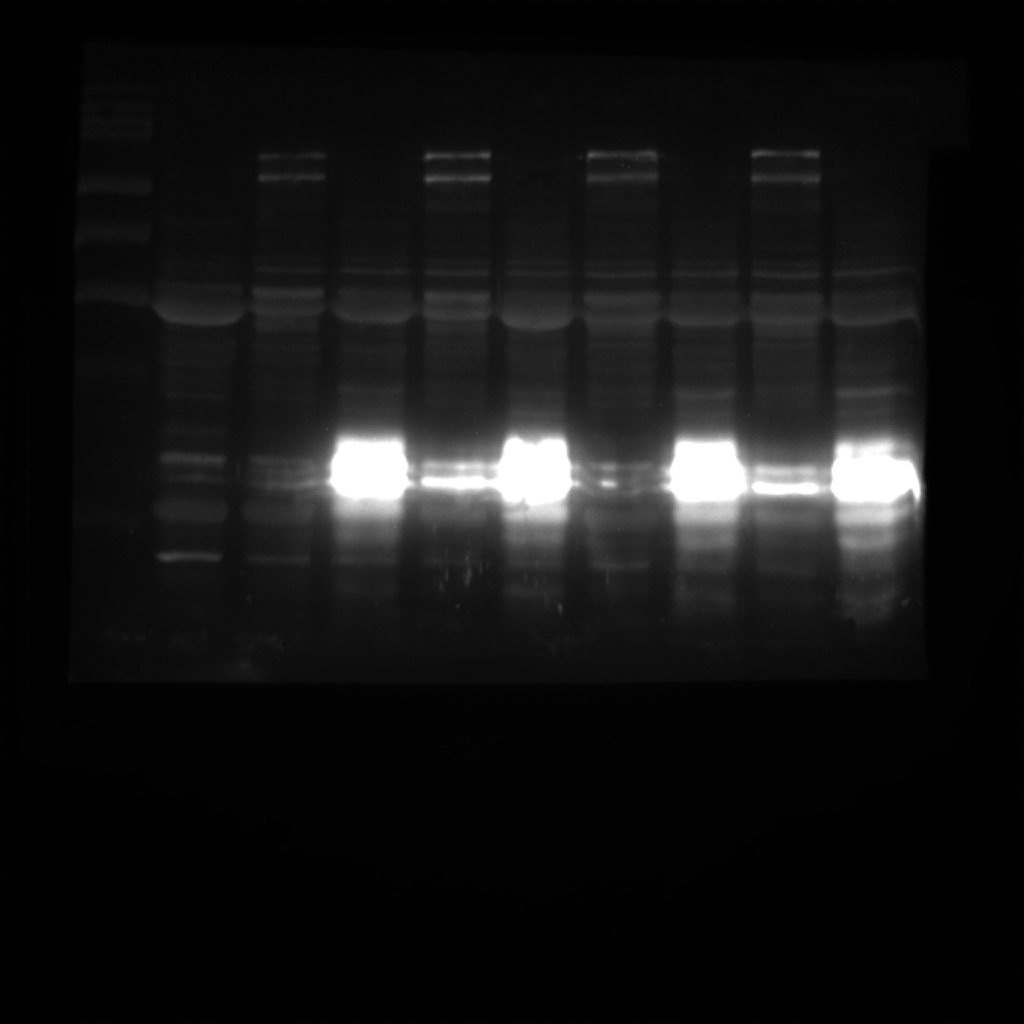

### IM000013 1min.Tif

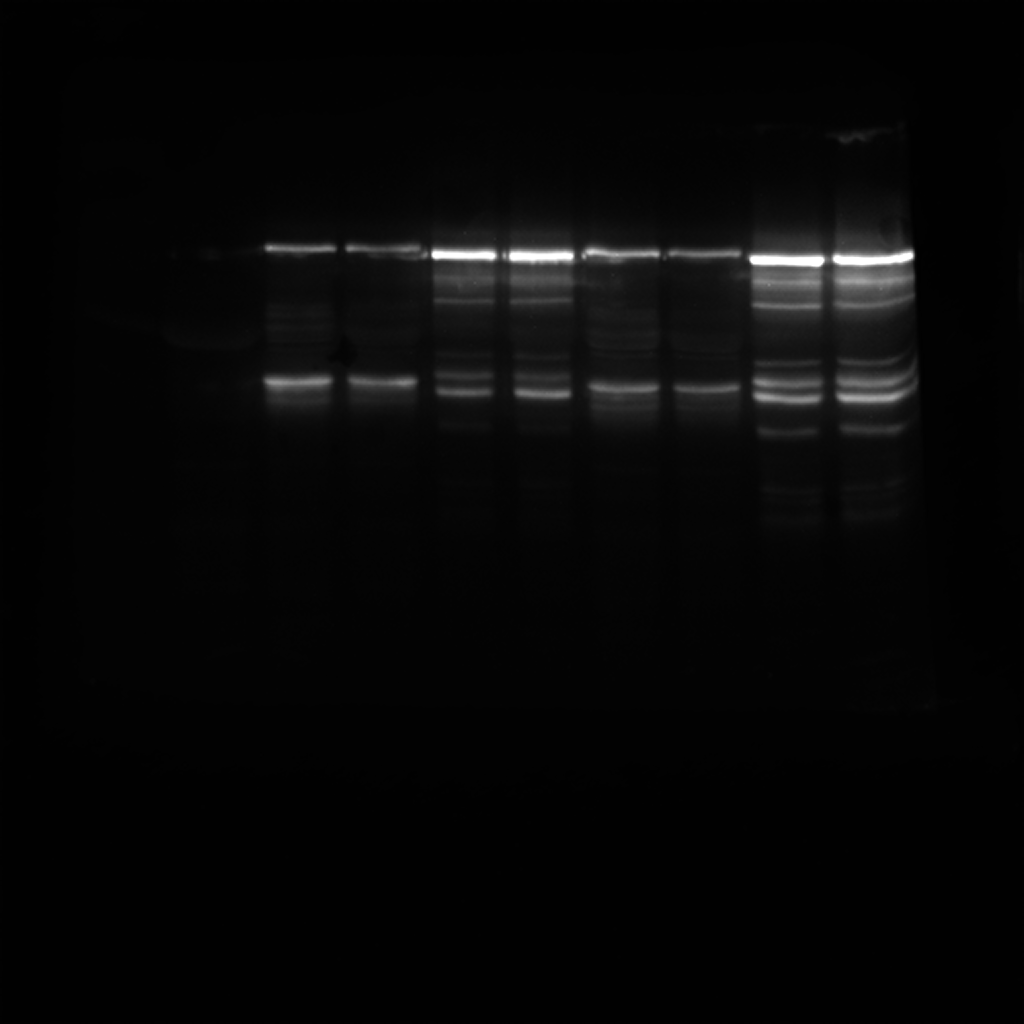

### Phost anti-GFP.Tif

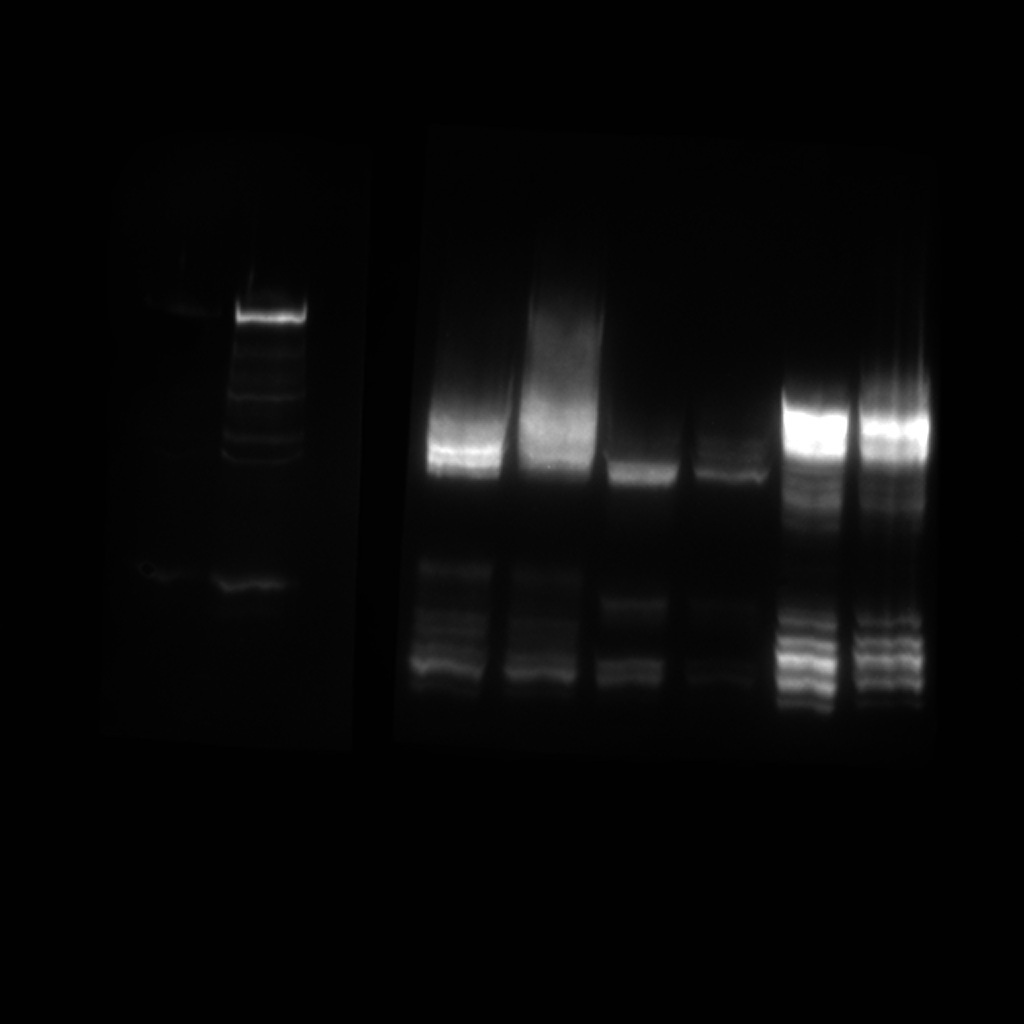

### Phostag antiGFP.Tif

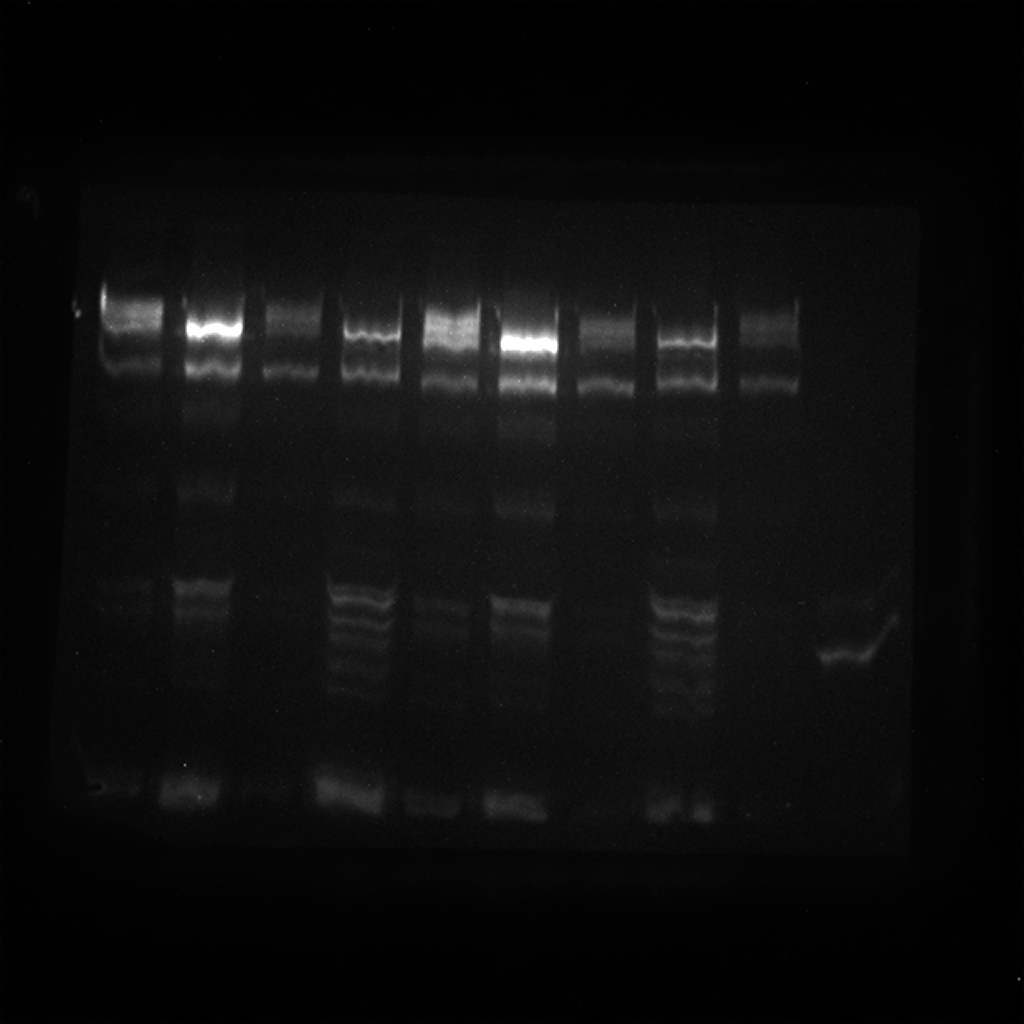

### Phostag GFP.Tif

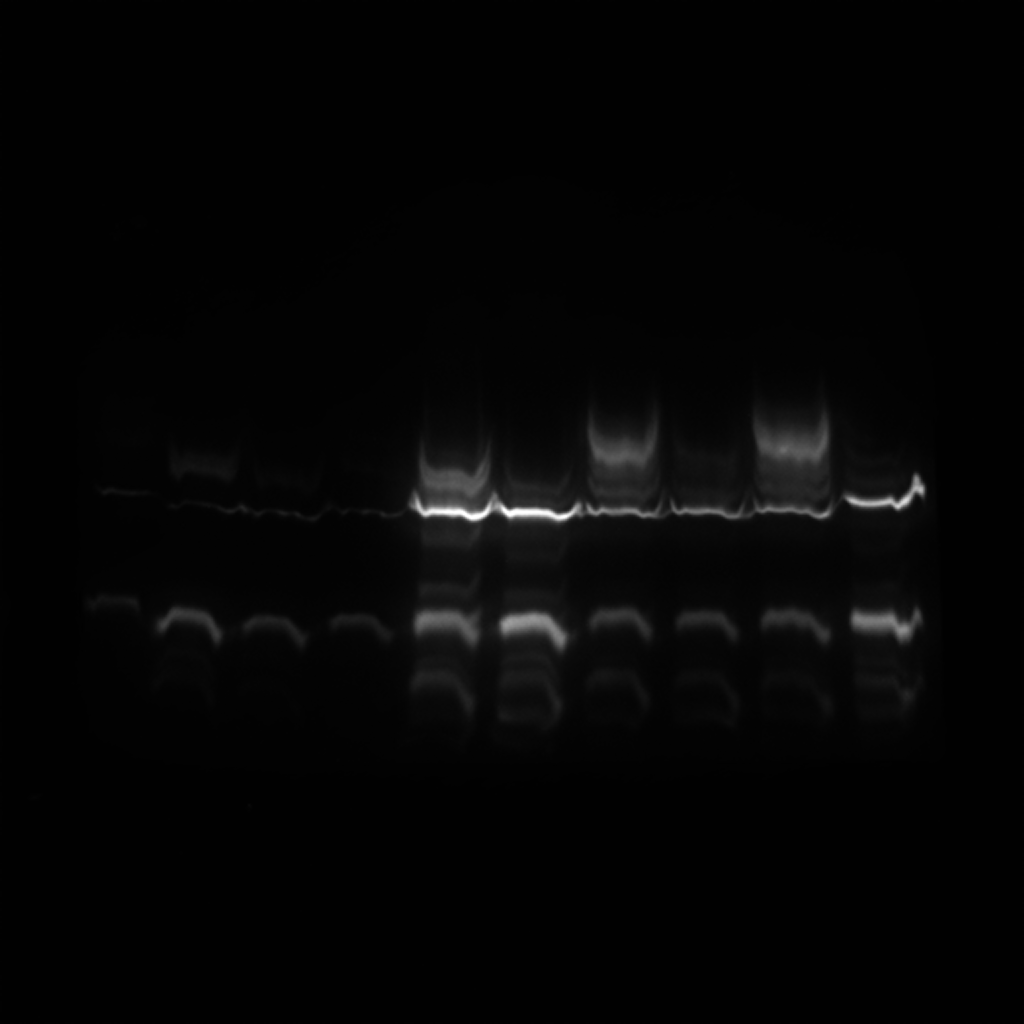

### PONCEAU.jpg

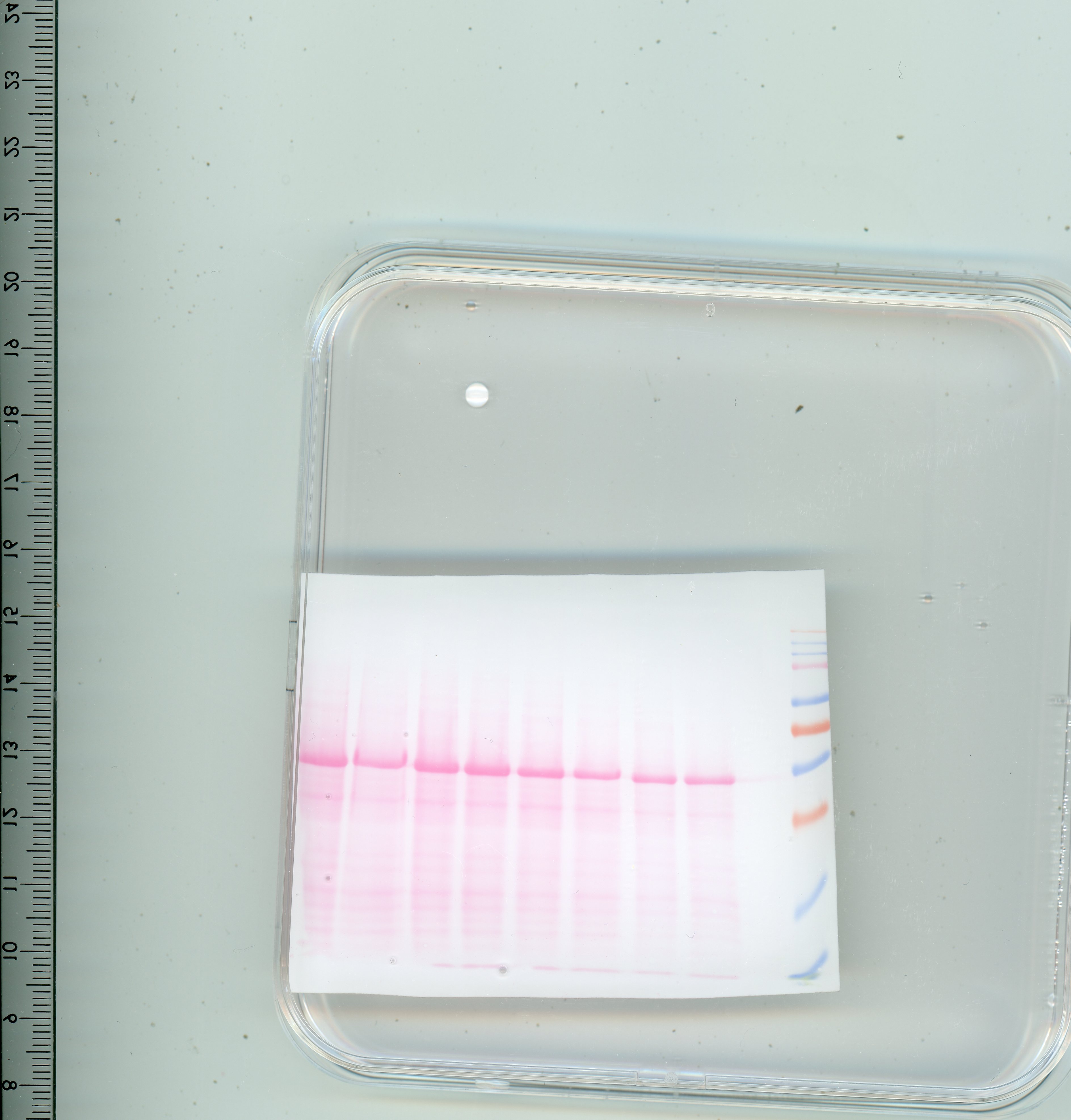

### ponceau.jpg

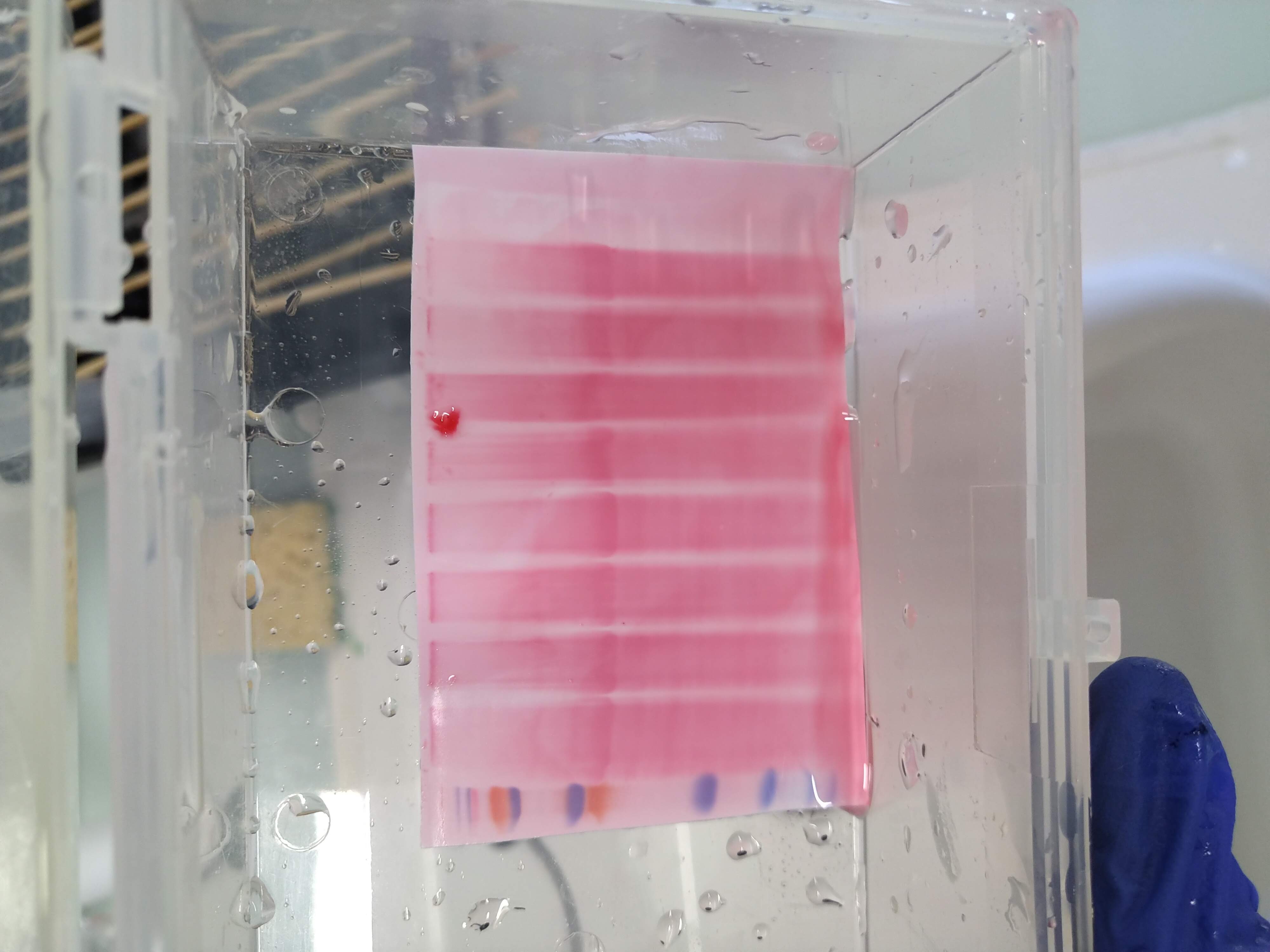

### ponceau.jpg

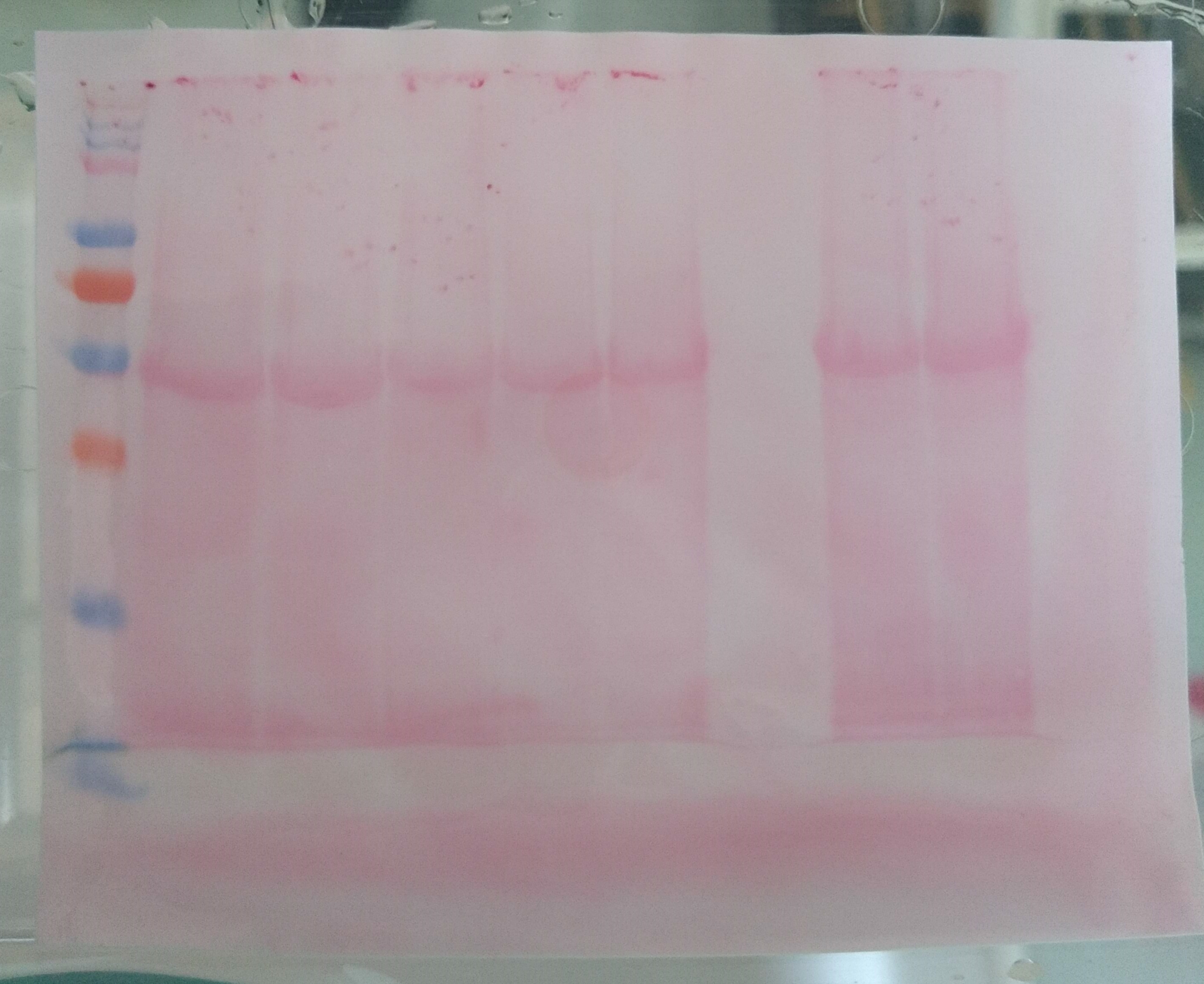

### preview ladder.Tif

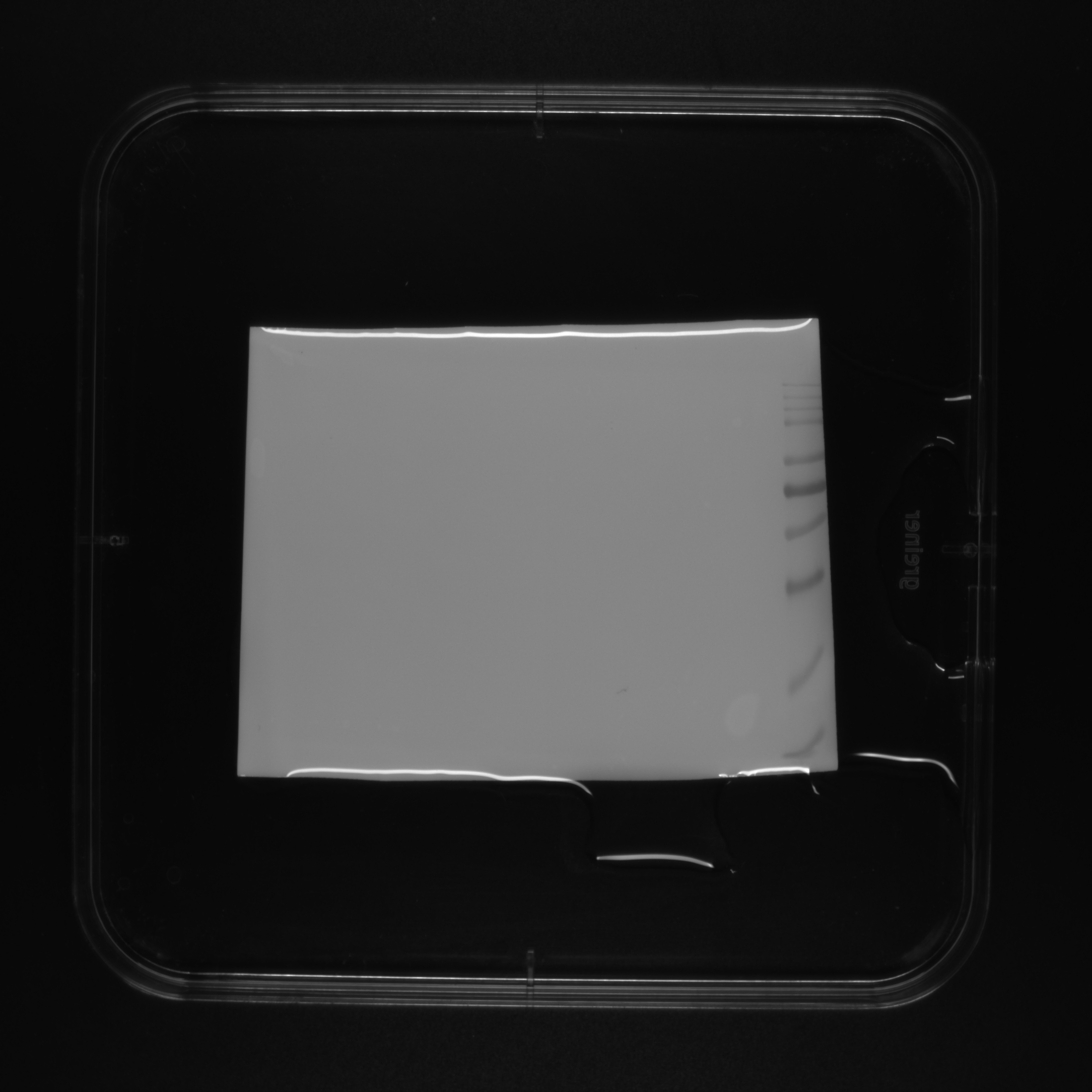

### S39_02_106BF1_exlList_bis.PNG

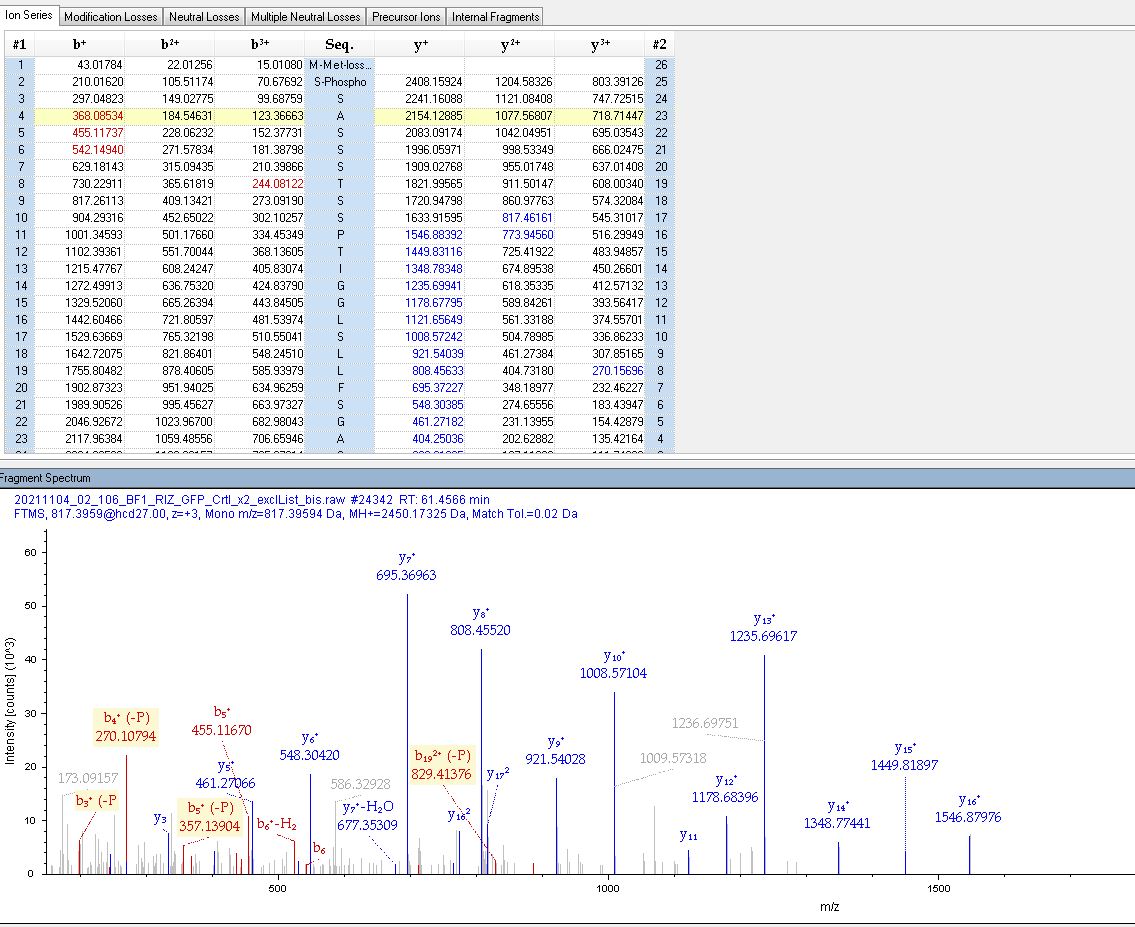

### S40_02_106BF1_exlList_bis.PNG

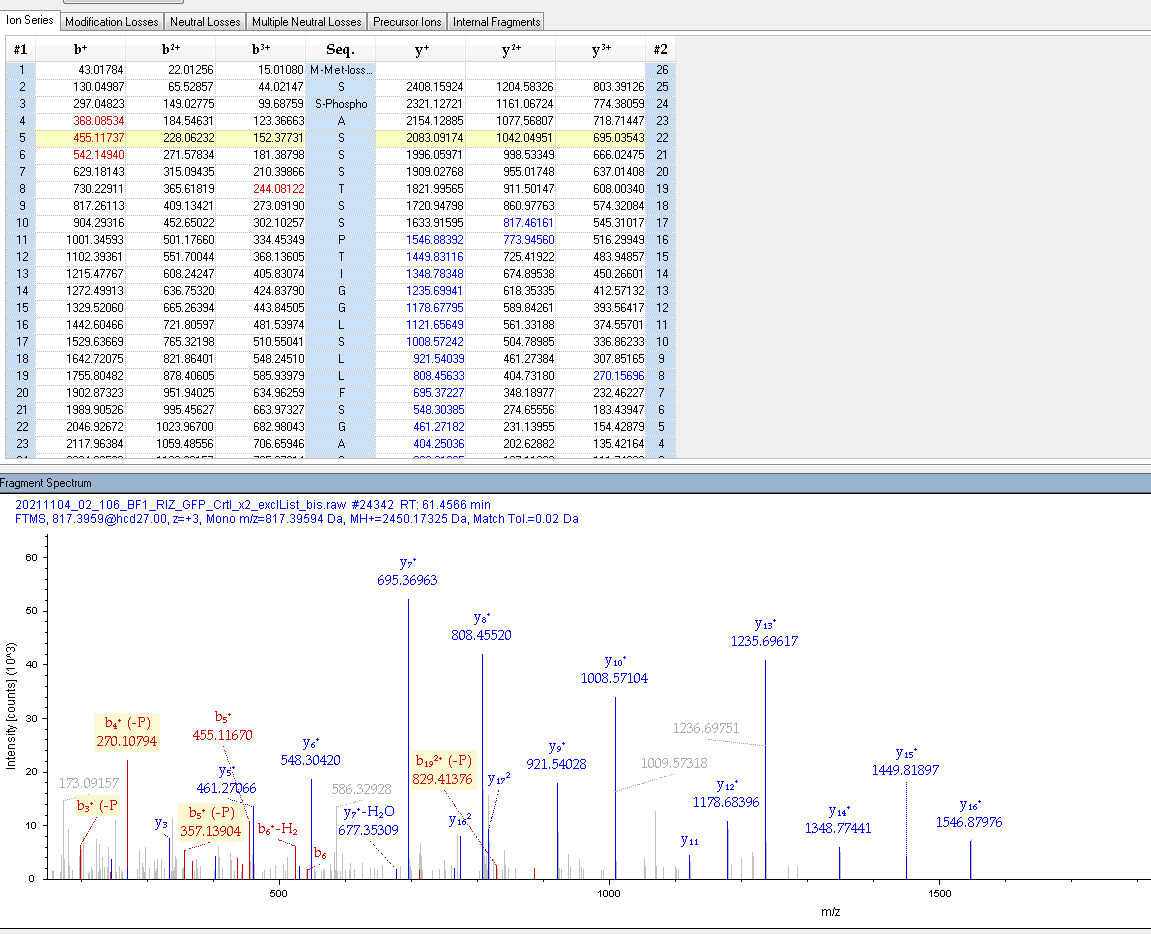

### S42_02_106BF1_exlList_bis.PNG

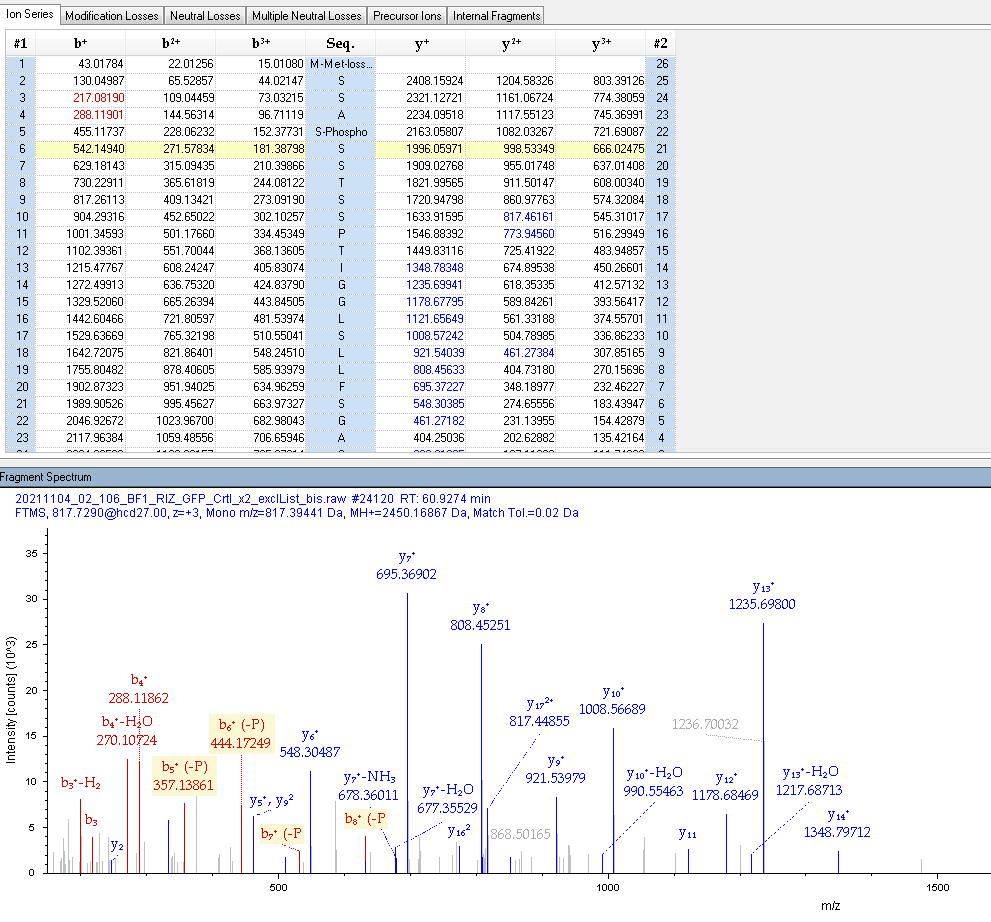

### S43_02_106BF1_exlList_bis.PNG

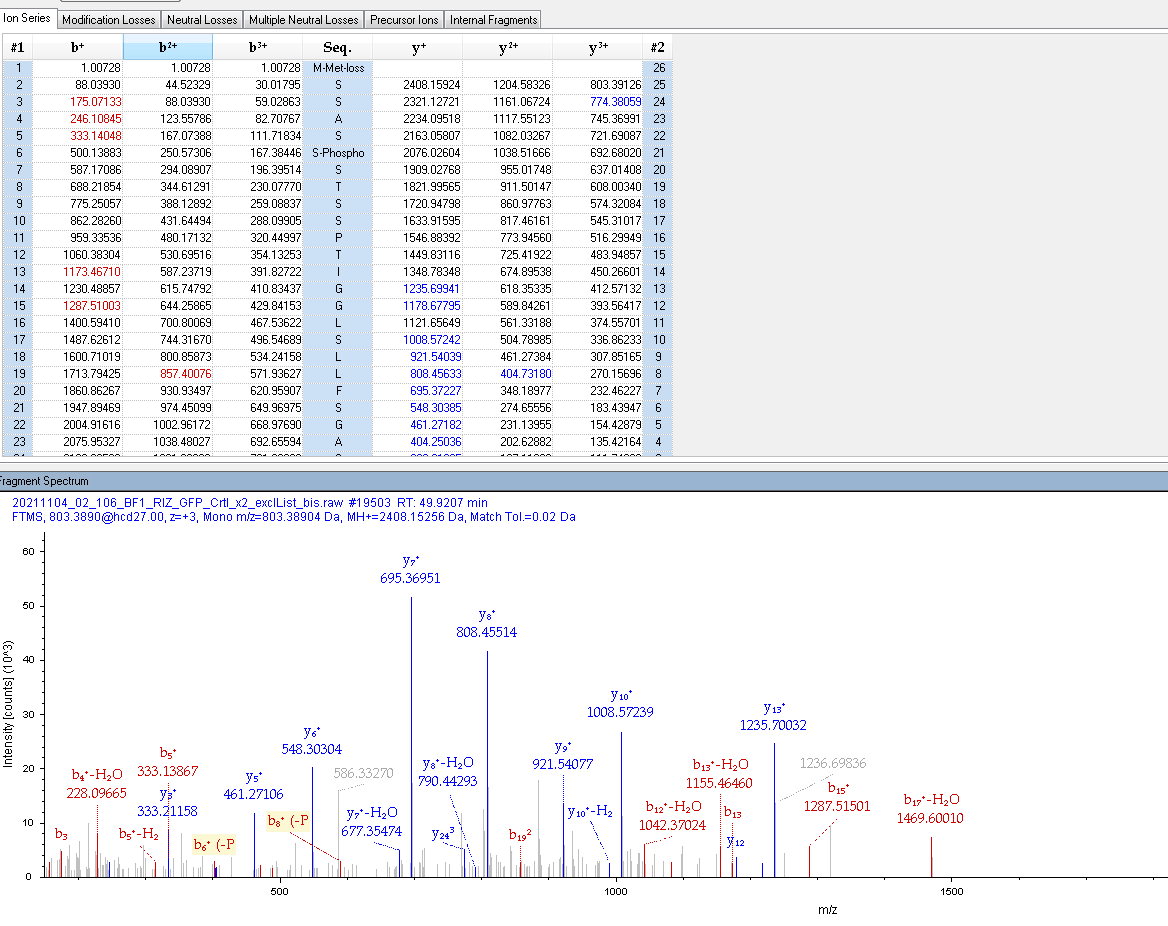

### S46_02_106BF1_exlList_bis.PNG

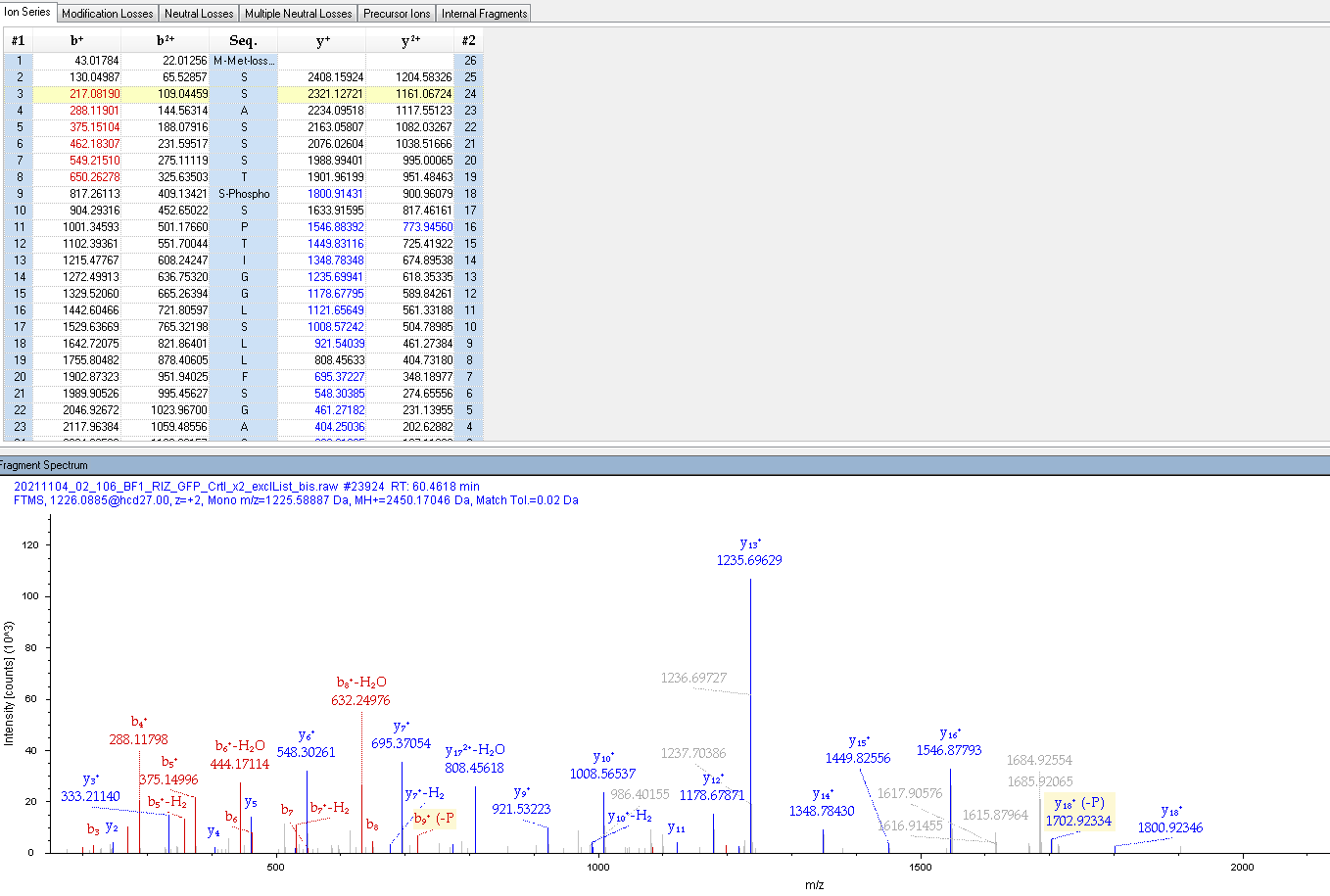

### S47_02_106BF1_exlList_bis.PNG

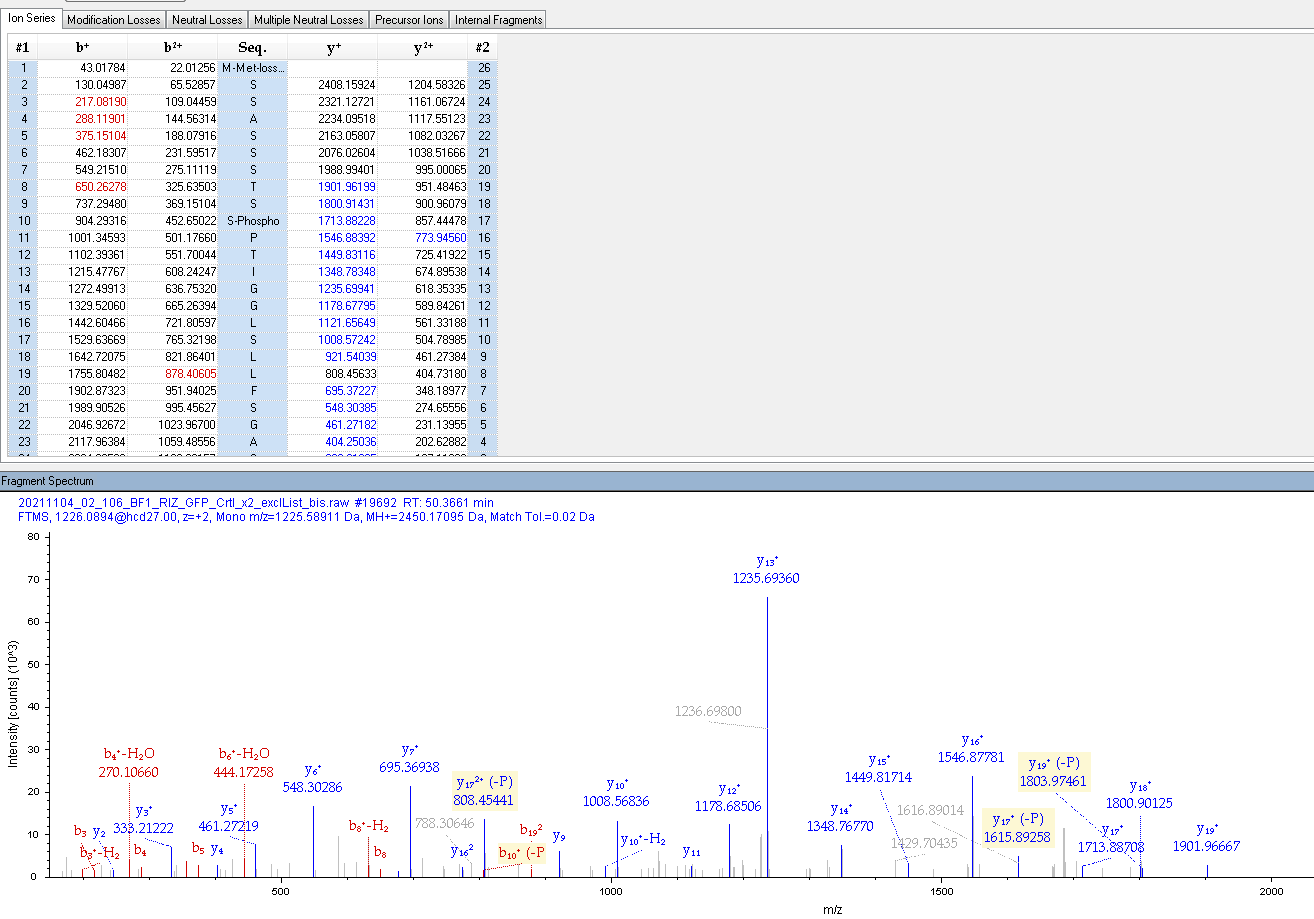

### S61_09_106BF1_exlList.PNG

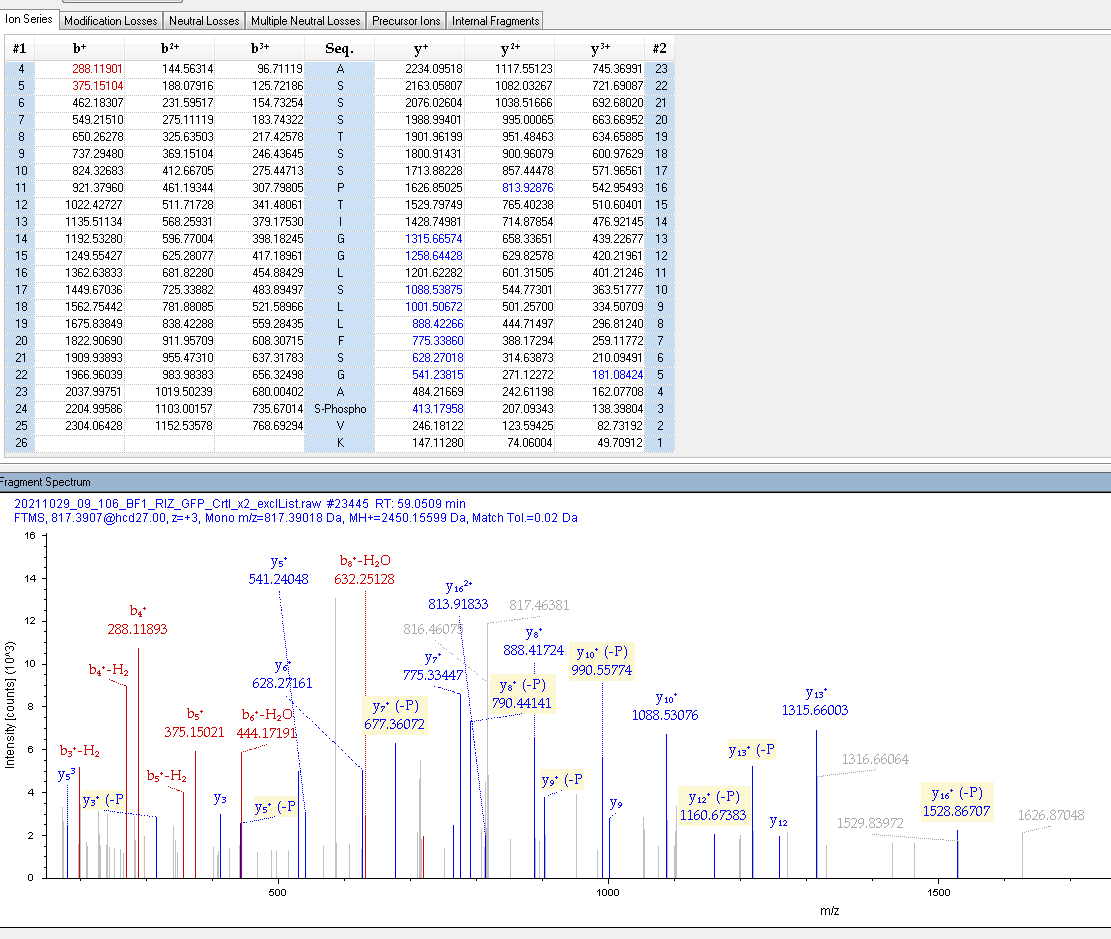

### S99_02_64BFS_trap_ter.PNG

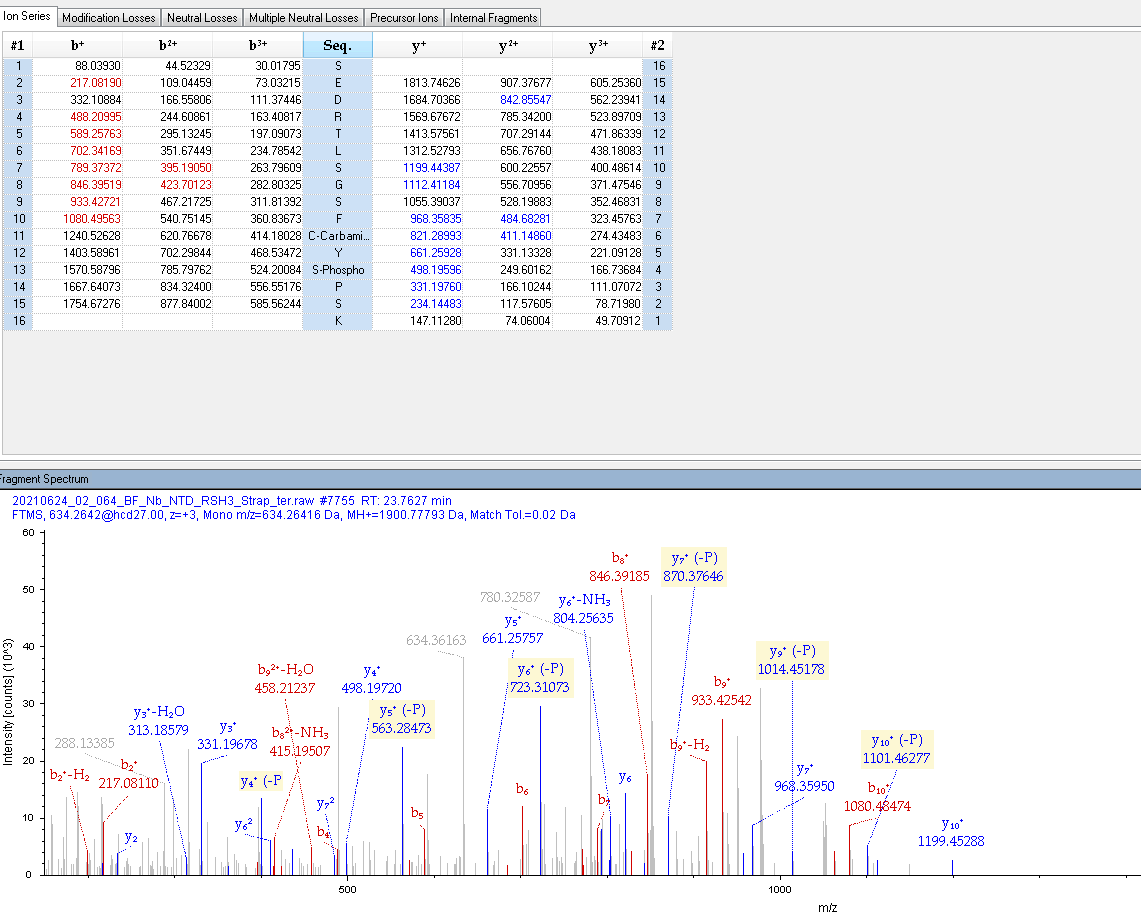

### S118&125_02_106BF1_exlList_bis.PNG

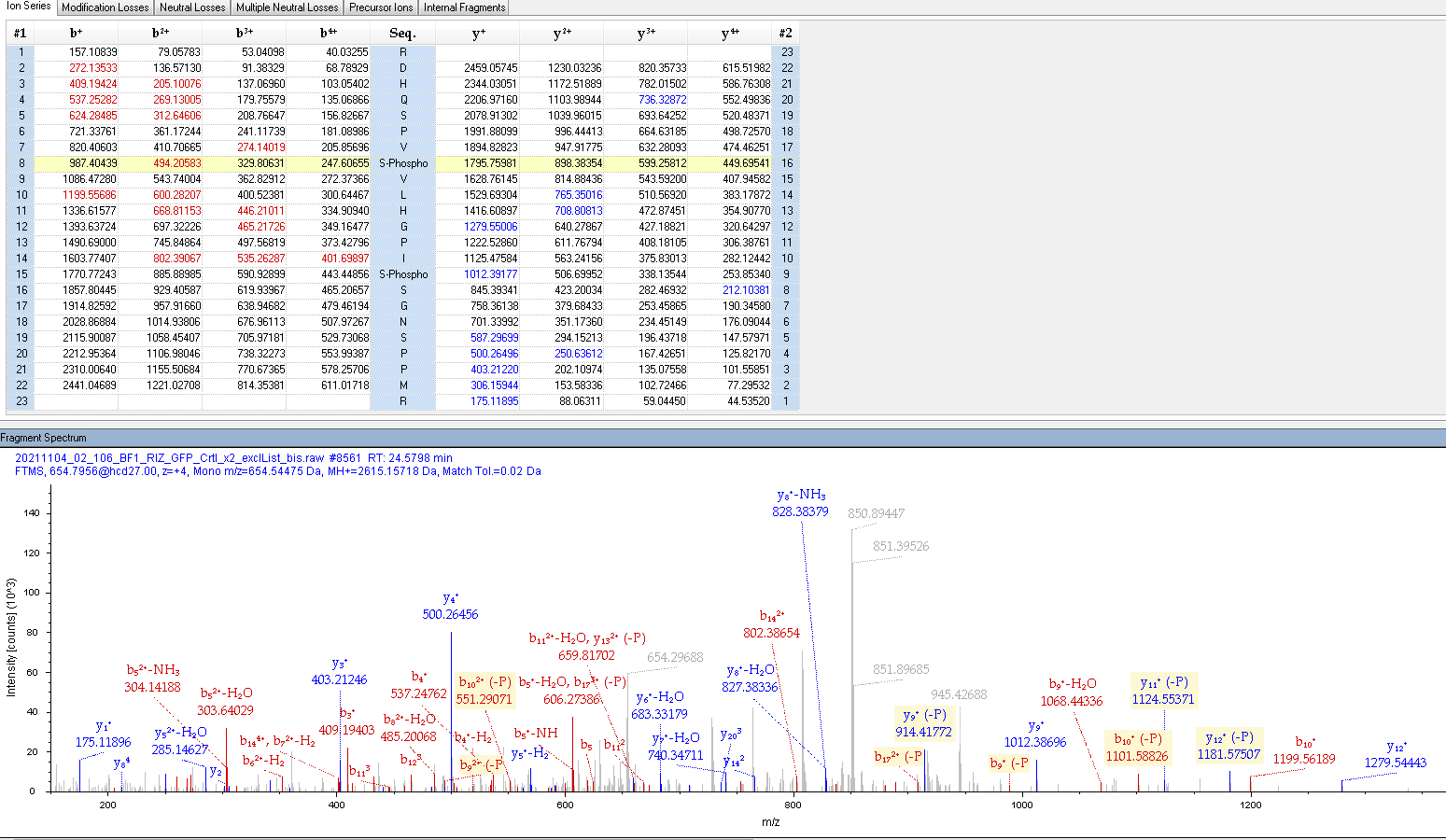

### S125_02_106BF1_exlList_bis.PNG

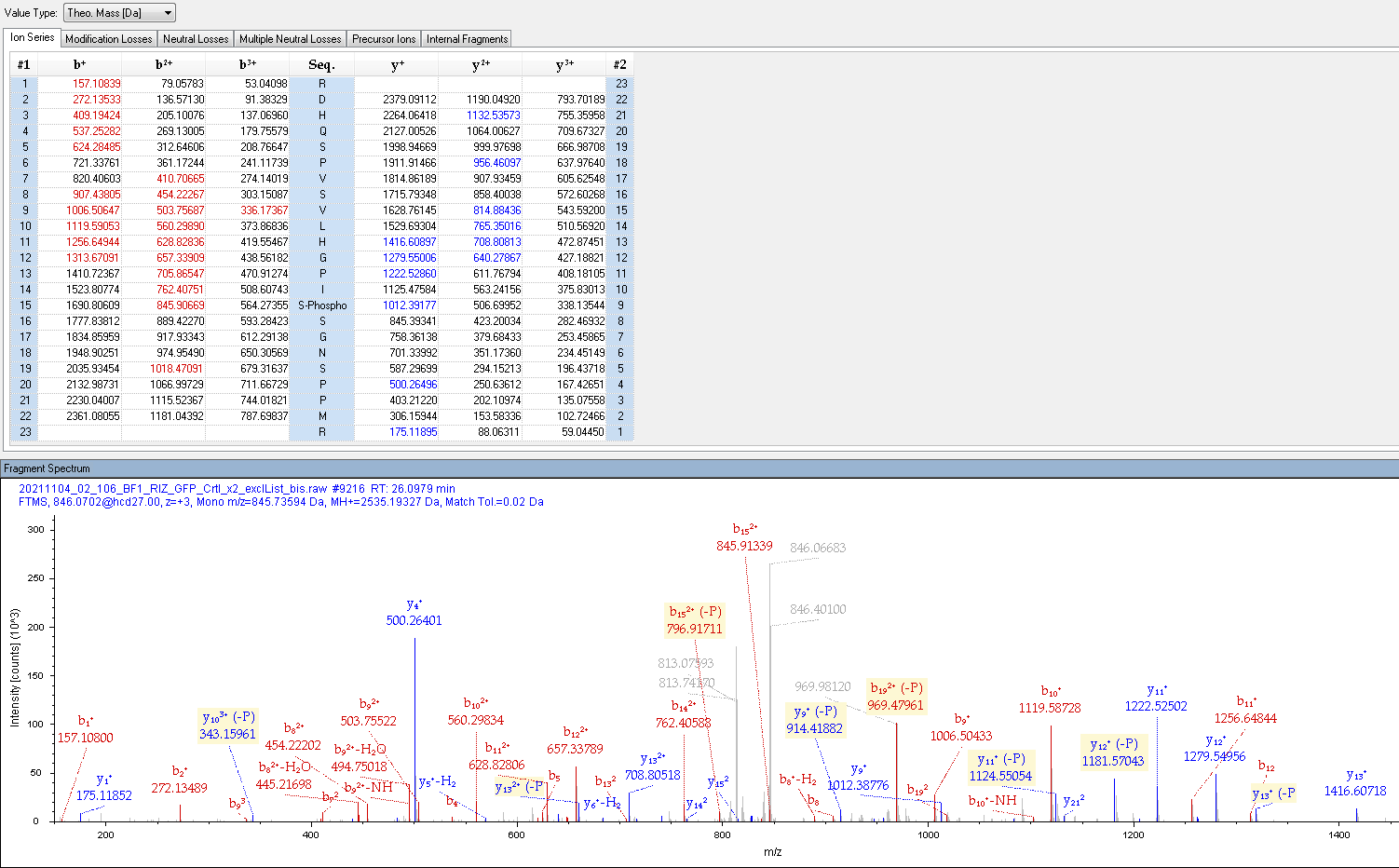

### S126_09_106BF1_exlList.PNG

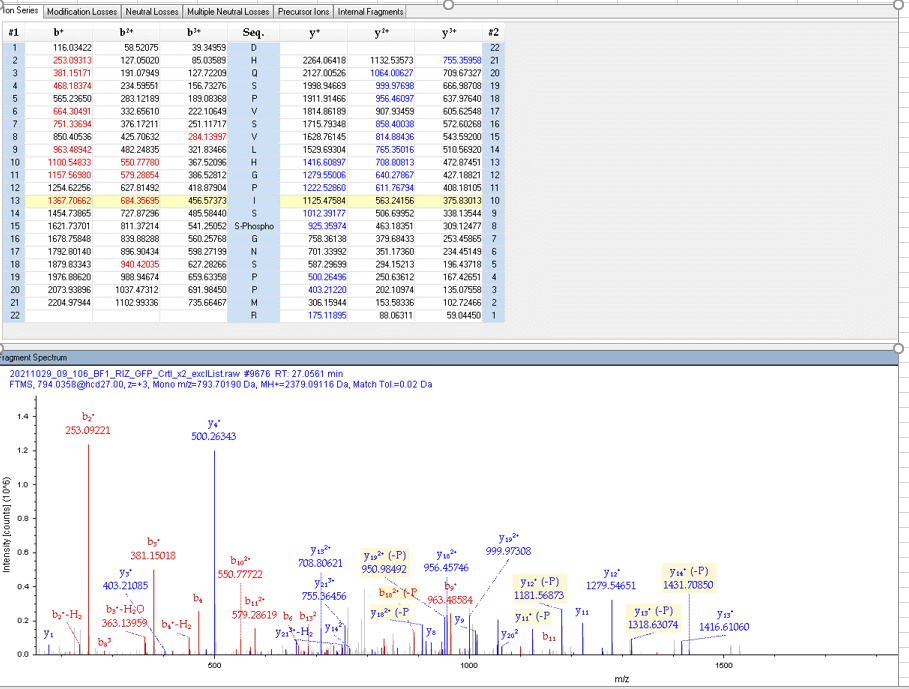

### S129_09_106BF1_exlList.PNG

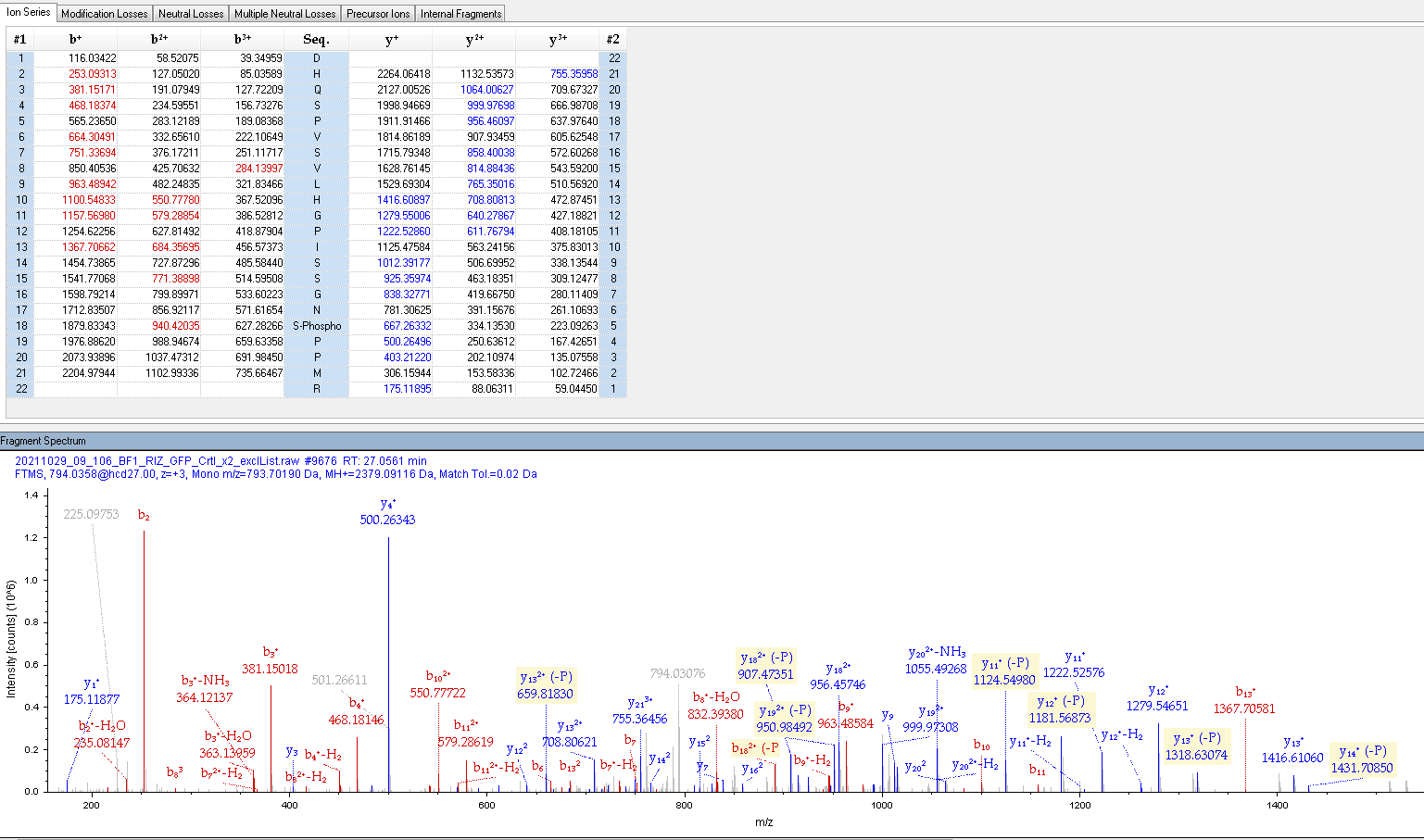

### S135_02_106BF1_exlList_bis.PNG

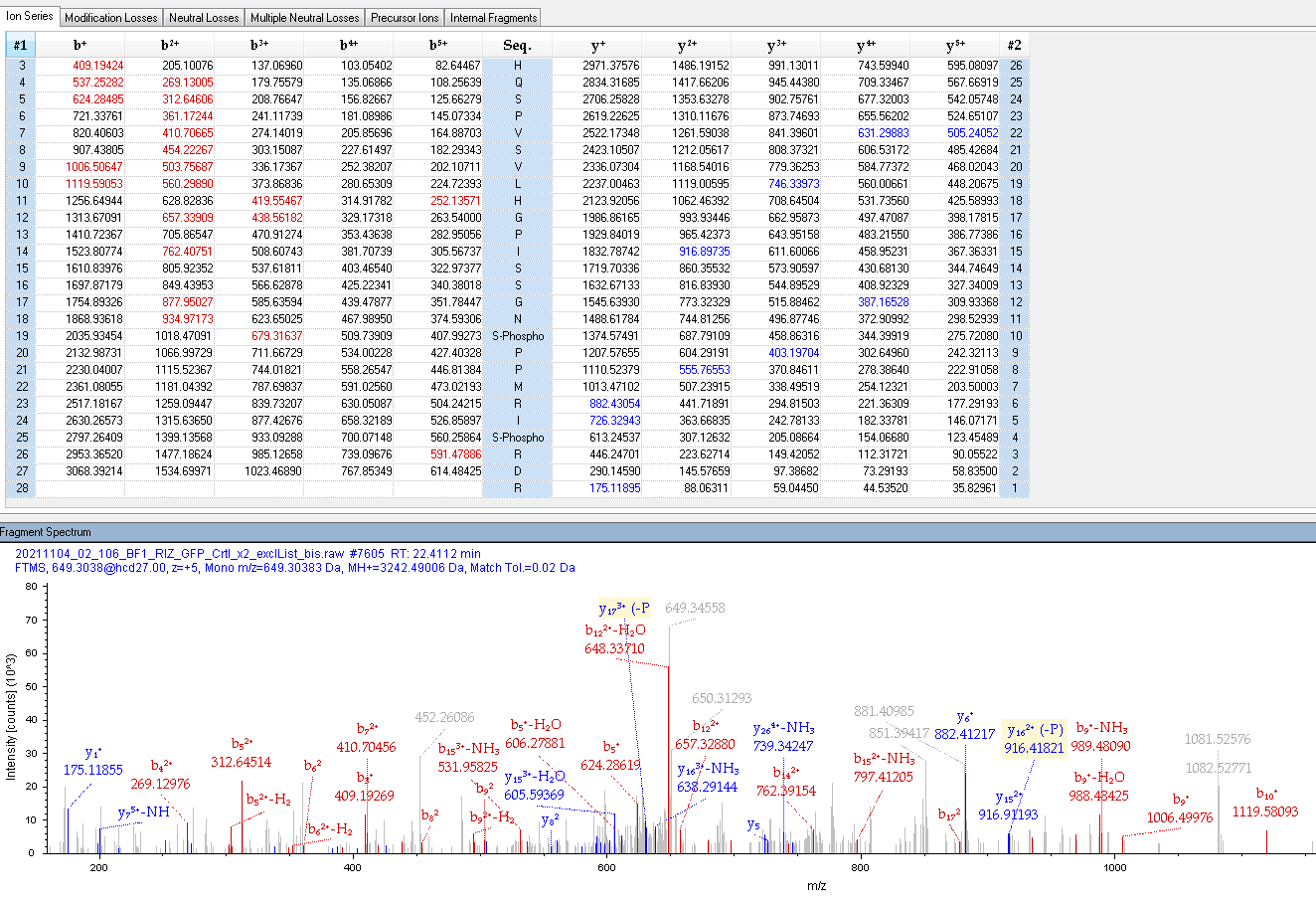

### S150_09_106BF1_exlList.PNG

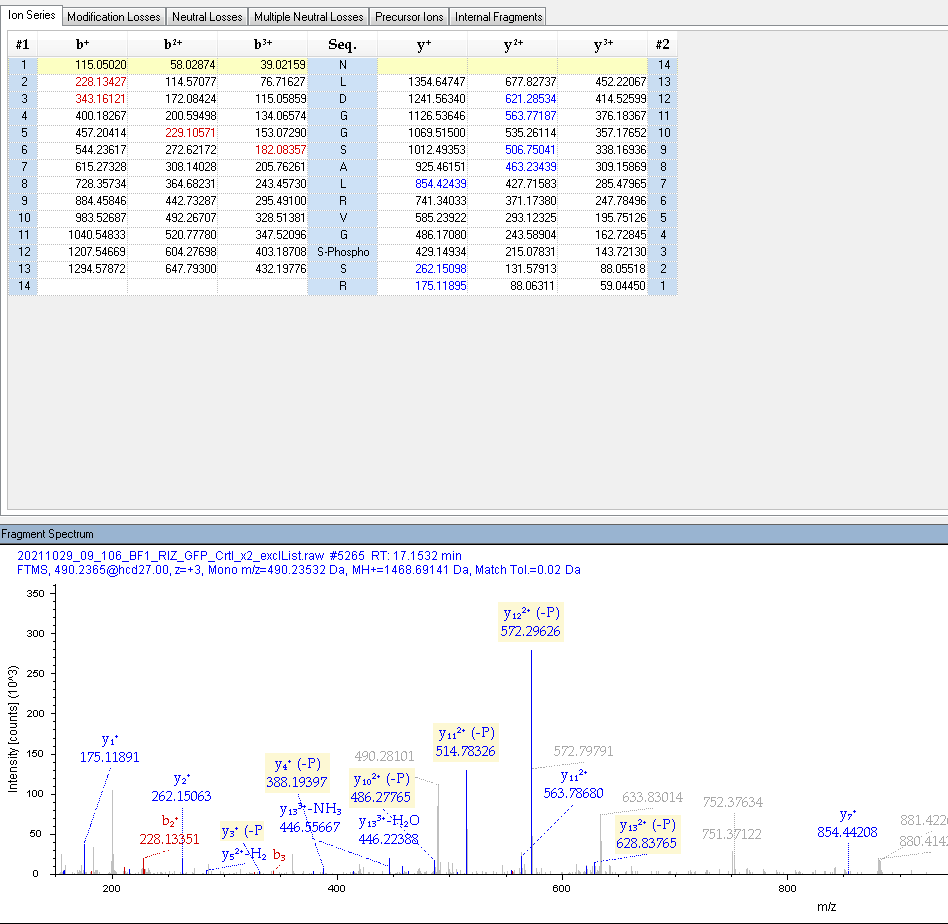

### SDS anti GFP.Tif

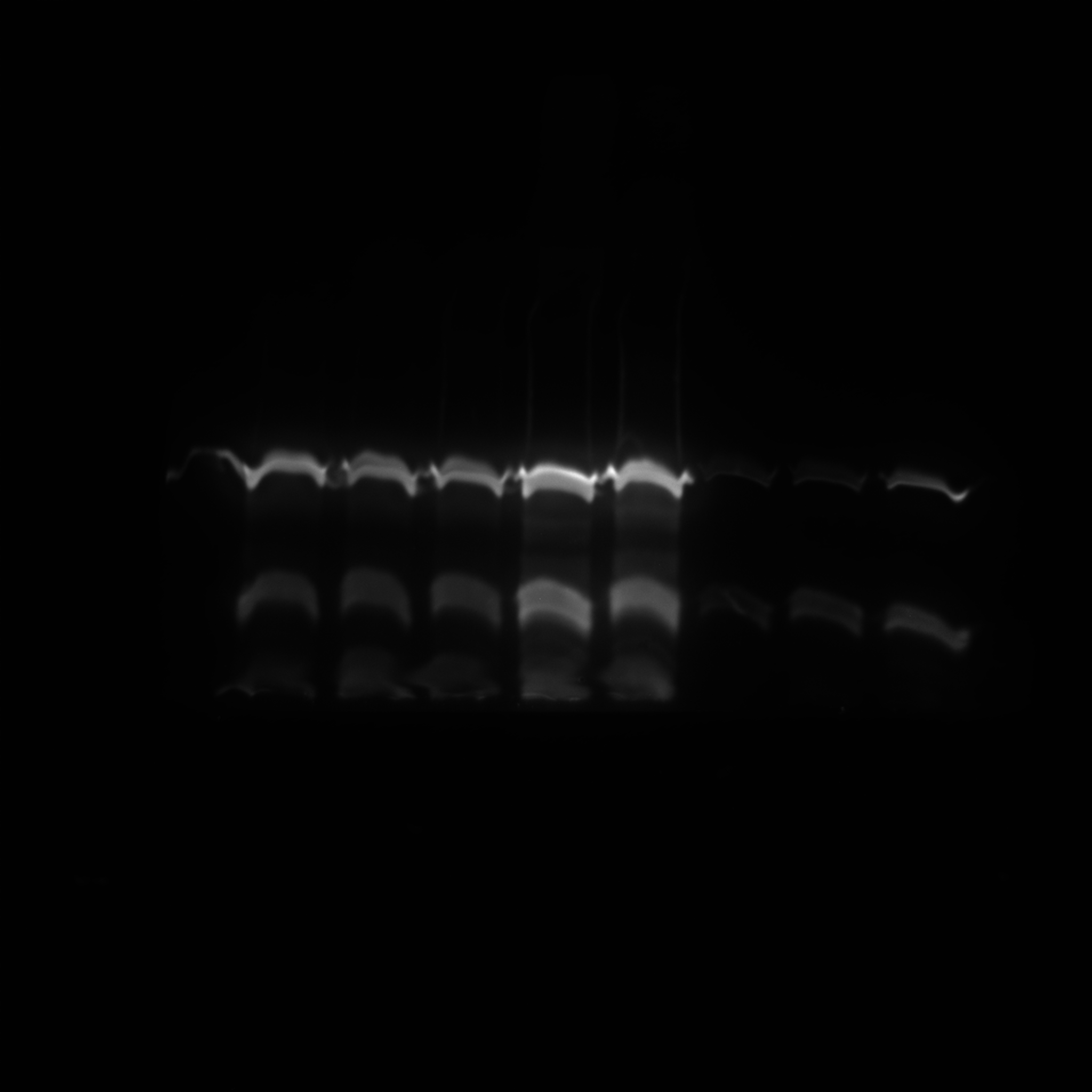

### SDS anti-GFP.Tif

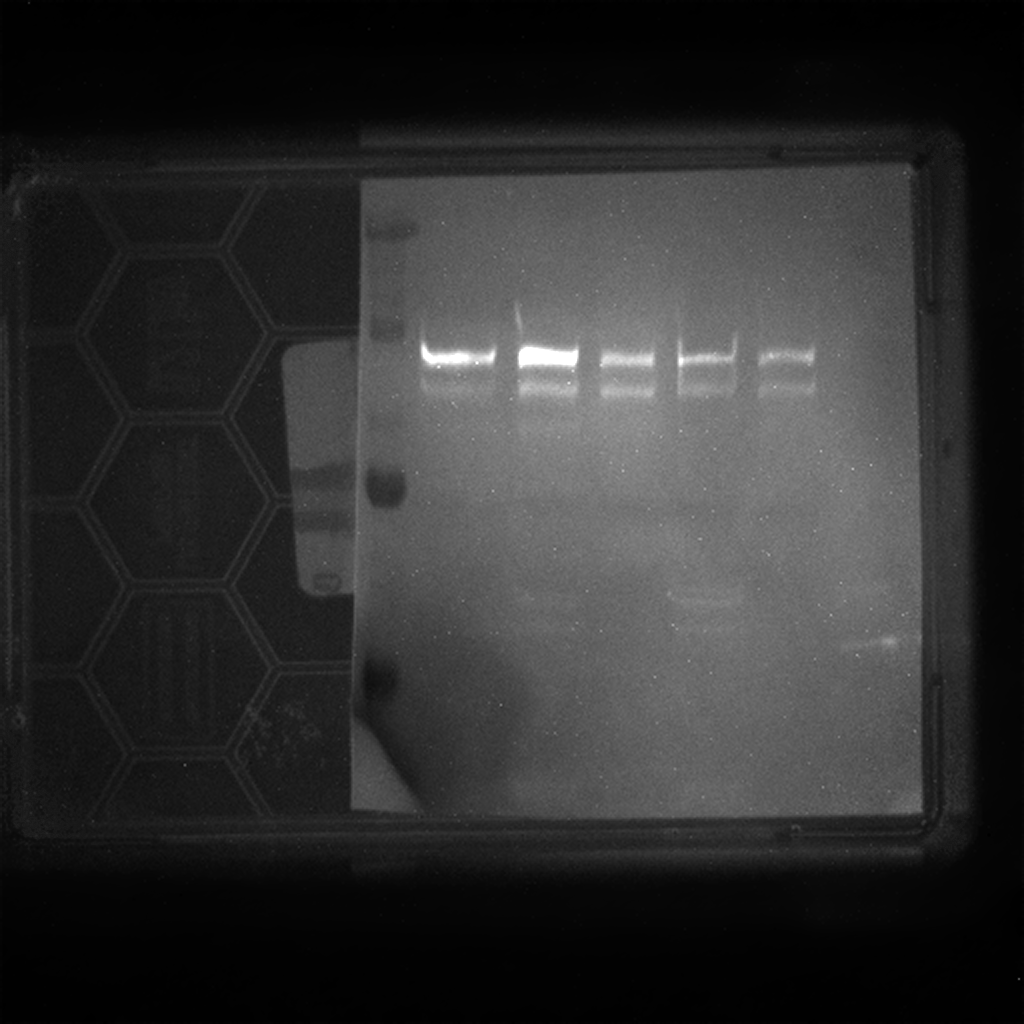

### SDS anti-GFP.Tif

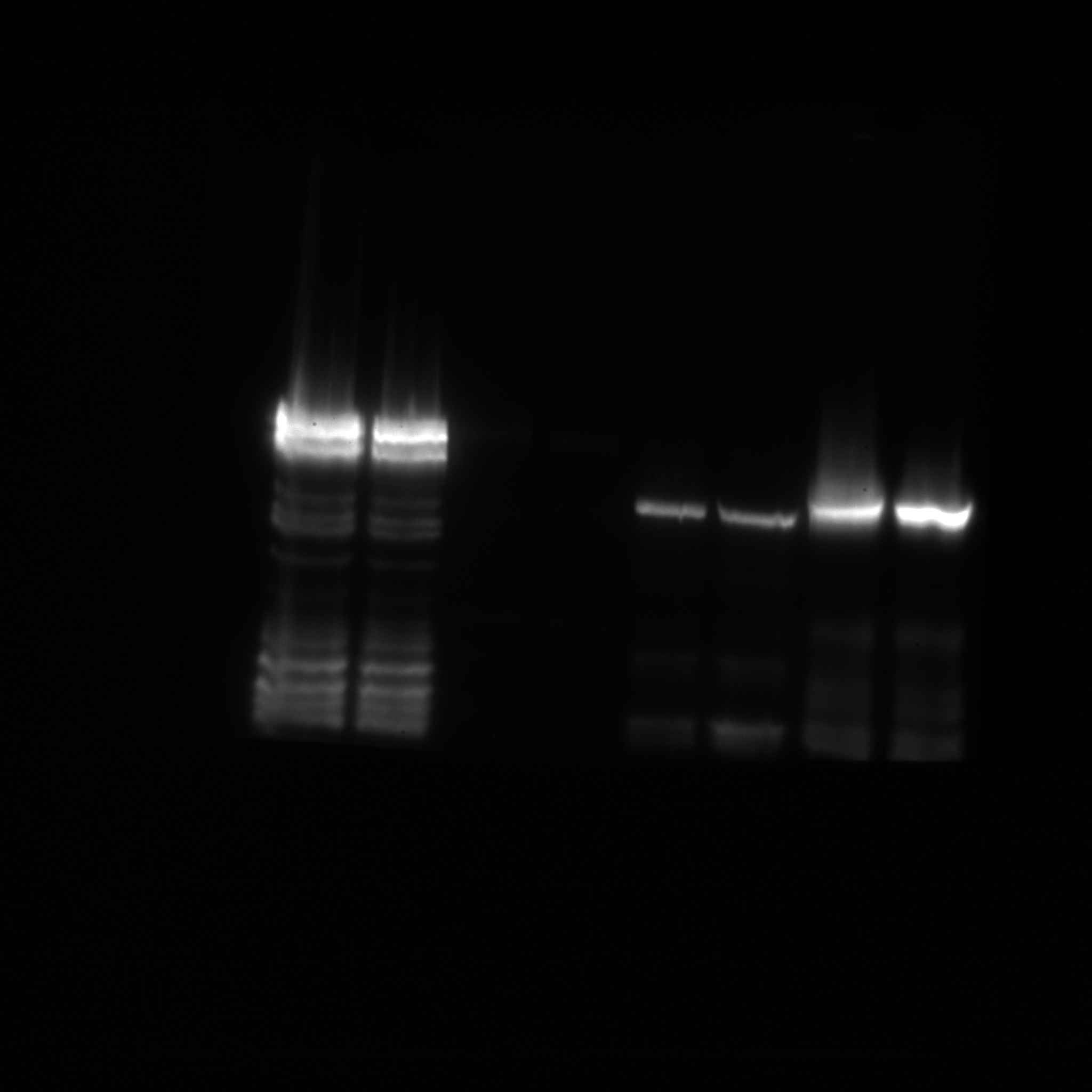

### white.Tif

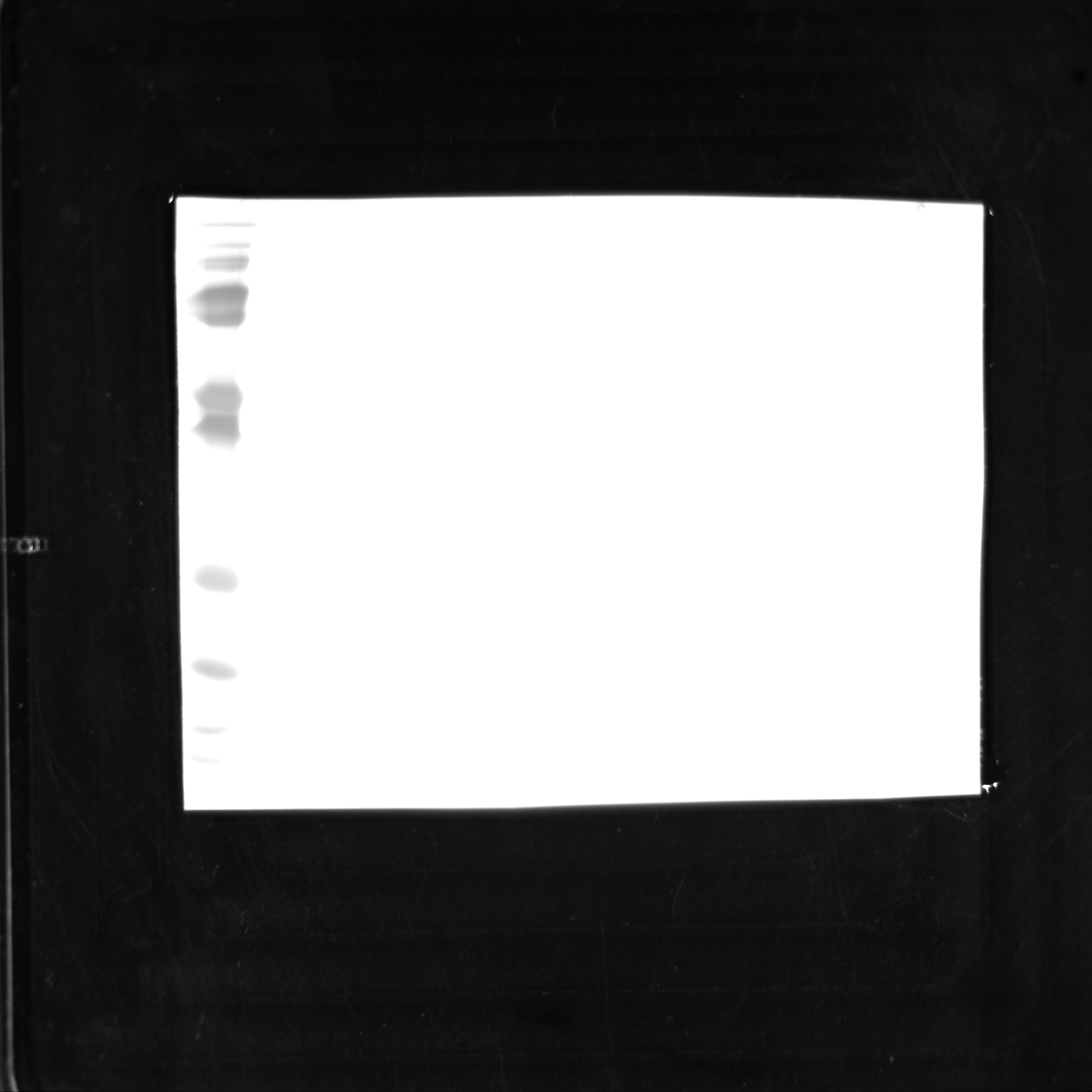
