## Supplementary Materials and Methods for "Post-translational regulation of photosynthetic activity via the TOR kinase in plants"

**Plant material**

Plant lines used and generated in this study are listed in the table of plant lines.

**Table of plant lines**

| **Plant species** | **Designation** | **Source or reference** | **Identifiers** | **Notes** |
| --- | --- | --- | --- | --- |
| *Arabidopsis thaliana* | Col-0 | Nottingham *Arabidopsis* Stock Centre (NASC) | WT, Columbia, N1093 (NASC) |  |
| *Arabidopsis thaliana* | *qrt1-2* | Nottingham *Arabidopsis* Stock Centre (NASC) | N8846 (NASC) | Col-3 ecotype, SAIL insertion parent line |
| *Arabidopsis thaliana* | *qrt1-2*/*rsh1-1* | Nottingham *Arabidopsis* Stock Centre (NASC), Sugliani et al. 2016 | TDNA insertion SAIL_391_E11, N818025 (NASC),  *rsh1* mutant | Col-3 ecotype |
| *Arabidopsis thaliana* | *qrt1-2*/*rsh2-1* | Nottingham *Arabidopsis* Stock Centre (NASC) , Sugliani et al. 2016 | TDNA insertion SAIL_305_B12, N814119 (NASC), *rsh2* mutant | Col-3 ecotype |
| *Arabidopsis thaliana* | *qrt1-2*/*rsh3-1* | Nottingham *Arabidopsis* Stock Centre (NASC), Sugliani et al. 2016 | TDNA insertion SAIL_99_G05, N862398 (NASC), *rsh3* mutant | Col-3 ecotype |
| *Arabidopsis thaliana* | *qrt1-2*/*rsh2-1 rsh3-1* | Nottingham *Arabidopsis* Stock Centre (NASC), Sugliani et al. 2016 | *rsh2,3* mutant | Col-3 ecotype |
| *Arabidopsis thaliana* | *Col-0/OX:RSH1* | Sugliani et al. 2016 | OX:RSH1, OX:RSH1-GFP (10.4) | Genomic sequence of RSH1 |
| *Arabidopsis thaliana* | *Col-0/OX:RSH3* | Sugliani et al. 2016 | OX:RSH3,  OX:RSH3-GFP.1 | Genomic sequence of RSH3 |
| *Arabidopsis thaliana* | *qrt1-2*/*rsh2-1 rsh3-1*  *OX:RSH3 line 1* | This study | *rsh2,3* OX:RSH3 line1 | Col-3 ecotype, CDS of RSH3 |
| *Arabidopsis thaliana* | *qrt1-2*/*rsh2-1 rsh3-1*  *OX:SSU-RSH3 line 1* | This study | *rsh2,3* OX:SSU-RSH3 line1 | Col-3 ecotype, CDS of RSH3 |
| *Arabidopsis thaliana* | *qrt1-2/rsh2-1 rsh3-1*  *OX:SSU-RSH3 line 2* | This study | *rsh2,3* OX:SSU-RSH3 line2 | Col-3 ecotype, CDS of RSH3 |
| *Arabidopsis thaliana* | *TOR/tor-1* | Menand et al., 2002 |  | T-DNA insertion, TOR GUS translational fusion. |
| *Arabidopsis thaliana* | *lst8-1* | Moreau et al., 2012 | SALK_02459*, lst8-1-1* | Non-viable seeds |
| *Arabidopsis thaliana* | *lst8-1 OX:RSH3* | This study | *lst8-1* OX:RSH3*,*  *lst8-1-1* OX:RSH3-GFP.1 | Not fertile, Genomic sequence of RSH3 |

**Plant growth conditions**

In each experiment, the seeds for each line were derived from the same batch of plants grown together. For growth on plates, seeds were surface sterilized with 70% ethanol 0.01% Triton X-100, rinsed with 100% ethanol, dried and placed on square culture plates containing 45 ml of 0.5× Murashige and Skoog salts (Merck Sigma-Aldrich), 0.5 g l^−1^ MES, and 0.8% Agar (Merck Sigma-Aldrich), adjusted to pH 5.7 with KOH. Plates were placed at 4°C for 2 days in the darkness, and then transferred to controlled growth conditions with 16 h/8 h photoperiod at 22 °C day/20 °C night. For non-sterile growth, *Arabidopsis thaliana* and *Nicotiana* *benthamiana* plants were grown in soil in a controlled environment at 120 µmol m^−2^ s^−1^ illumination with an 16 h/8 h photoperiod at 22 C day/20°C night, and 55% day/75% night relative humidity. Plants were treated weekly with a Coïc-Lesaint complete nutrient solution.

**Cloning**

New gene parts (level 0 modules) were amplified by PCR or synthesized directly (Twist Biosciences) and sequenced. The resulting modules are free from BsaI, BsmBI, BpiI and SapI Type IIS sites and can be mobilised in the MoClo cloning system (Patron et al., 2015). Restriction ligation reactions for the assembly of transcriptional units (Level 1) and assemblies of transcriptional units (Level 2) were performed using a single step protocol as described previously with small modifications (Weber et al., 2011) and according to the MoClo Golden Gate assembly standard (Engler et al., 2014; Gantner et al., 2018) (for detailed instructions see cloning guide in Velay et al. (2022)). Briefly, 100 fmol of each insert plasmid and 50 fmol of acceptor plasmid were mixed with restriction enzyme (BpiI or BsaI) and T4 DNA ligase in restriction enzyme buffer and 1mM ATP in 20 µl reactions and incubated at 37 °C for 5 h. A 1.5 µl aliquot was transformed into DH10B *E. coli* cells by electroporation and transformants selected on appropriate antibiotics. Correct assembly was confirmed by digestion. Full sequences of new modules are available in the supplementary data files. All other Level 0 and infrastructure modules used were described previously (Engler et al., 2014; Gantner et al., 2018; Velay et al., 2022). The RSH3 phosphodefective peptide was constructed by mutating all serine residues that we initially predicted to be susceptible to TOR-dependent regulation, as well as S61 whose phosphorylation was experimentally identified (see also Fig. S5)(Xi et al., 2021). The RSH3 phosphomimic peptide was constructed by mutating serine residues after the predicted chloroplast transit peptide cleavage site at position 64.

**Table of primers**

| **Primer name** | **Sequence** | **Direction** |
| --- | --- | --- |
| RSH3_65-715_ | ttgaagacttATCATCTTCCTCTTCTTCCTCATCG | For |
| RSH3_65-715_ | ttgaagacttCGAACTTCCCCAGCCAACC | Rev |
| RSH3_39-221_ | ttggtctcaAATGTCATCGGCGTCCTCTTCCACG | For |
| RSH3_39-221_ | ttggtctcacgaaTCGTAAAATGCTTTGATC | Rev |
| RSH3_131-715_ | ttgaagacttCCTCCGATGAGGATTTCACG | For |
| RSH3_195-715_ | atggtctcaAGGTCCATATGCTAGGGATTTG | For |
| RSH3_195-715_ | atggtctcacgaaCTTCCCCAGCCAACCATGG | Rev |
| RSH3_1-64_ | atggtctcaAATGGTGGTAGCAACGACC | For |
| RSH3_1-64_ | atggtctcaACCTatTGAGGAAGAAGAGGAAGATGATTTGAC | Rev |
| RSH3_1-96_ | atggtctcaACCTatGAAGGATCCGCTTAAGGTCC | Rev |
| RSH3_1-135_ | atggtctcaACCTatCCTCATCGGAGGACTGTTACC | Rev |
| RSH3_1-165_ | atggtctcaACCTatACATGAACCTATCGCTTTCCTGAC | Rev |
| RSH3_1-175_ | atggtctcaACCTatGACAAGAACAGAGTCTGTATCGTAATCAACAC | Rev |
| SSU-CTP_57-69_ | ttggtctcaAATGgtgtggcctccgattggaaag | For |

**Table of genetic constructions**

| **Name** | **Moclo Level** | **Notes** |
| --- | --- | --- |
| pEarleygate 103 35S:RSH3-GFP | NA | Sugliani et al., 2016 |
| TID | L0 | Velay et al., 2021 |
| RSH3_1-64_ | L-1 | Synthesized |
| RSH3_1-130_ S12A, S40A, S47A, S61A, S99A, S115A, S129A | L-1 | Synthesized |
| RSH3_1-130_ S99E, S115E, S129E | L-1 | Synthesized |
| LST8 (At3g18140) CDS | L0 | cds2ns |
| RSH3_65-715_ CDS | L0 | Synthesized |
| RSH3_65-715_ D452G CDS | L0 | Synthesized |
| RSH3_1-715_ CDS | L0 | cds1ns |
| RSH3_1-715_ S12A, S40A, S47A, S61A, S99A, S115A, S129A | L0 | cds1ns |
| RSH3_1-715_ S99E, S115E, S129E | L0 | cds1ns |
| Cyanophora RSH2/3_90-282_ | L0 | Synthesized |
| Rice RSH2/3_34-216_ | L0 | Synthesized |
| Tomato RSH2/3_43-226_ | L0 | Synthesized |
| 35S:SSU_1-79_-CFP | L1 | Velay et al., 2021 |
| 35S:TID-LST8 | L1 |  |
| 35S: LST8-TID | L1 |  |
| 35S:TID-YFP | L1 |  |
| 35S:mCHERRY-LST8 | L1 |  |
| 35S: RSH3_1-715_-GFP | L1 |  |
| 35S: RSH3_1-715_ S12A, S40A, S47A, S61A, S99A, S115A, S129A -GFP | L1 |  |
| 35S: RSH3_1-715_ S99E, S115E, S129E -GFP | L1 |  |
| 35S:RSH3_39-221_-GFP | L1 |  |
| 35S:RSH3_39-221_ S40A, S47A, S61A, S99A, S115A, S129A -GFP | L1 |  |
| 35S:SSU_57-79_-RSH3_65-221_-GFP | L1 |  |
| 35S:SSU-RSH3_65-715_-GFP | L1 |  |
| 35S:SSU-RSH3_194-715_ D452G-GFP | L1 |  |
| 35S:SSU_1-79_-GFP | L1 |  |
| 35S:GFP | L1 |  |
| 35S:RSH_1-70_-GFP | L1 |  |
| 35S:RSH_1-96_-GFP | L1 |  |
| 35S:RSH_1-135_-GFP | L1 |  |
| 35S:RSH_1-165_-GFP | L1 |  |
| 35S:RSH_1-175_-GFP | L1 |  |
| 35S:Cyanophora RSH2/3_90-282_-GFP | L1 |  |
| 35S:Rice RSH2/3_34-216_-GFP | L1 |  |
| 35S:Tomato RSH2/3_43-226_-GFP | L1 |  |

**Yeast two hybrid screening**

Yeast two-hybrid (Y2H) screening was performed by Hybrigenics Services (Paris, France). The coding sequence of LST8 (At3g18140, residues 1-305) was amplified from Arabidopsis cDNA and cloned into pB66 (GAL4 N-terminal fusion) as baits. The prey library was derived from one-week old Arabidopsis seedlings. In total 69 million (pB66_C) interactions were examined, and 89 clones processed. Four clones containing fragments of RSH2 were identified with high confidence in the interaction, and two clones containing fragments of RSH3 were identified with good confidence in the interaction.

**Transient expression by agroinfiltration**

*Agrobacterium tumefaciens* GV3101 transformed with plant expression constructs were grown at 28°C overnight in LB medium supplemented with rifampicin and a selective antibiotic. The cultures were then diluted to an OD_600_ of 0.2 in infiltration buffer containing 10 mM MES pH 5.5, 10 mM MgCl_2_ and 200 µM acetosyringone, and then infiltrated into leaves of one month old *N. benthamiana* plants using a 1 ml syringe. Infiltrated plants were returned to standard growth conditions for three days before observation or further treatment.

**Biotin labelling**

Leaf discs or whole leaves were taken from *N. benthamiana* plants three days after agroinfiltration and vacuum infiltrated with a solution of 50 µM biotin. The plant material was then floated on water and transferred to standard growth conditions for 2 hours before being rinsed with cold water and flash frozen in liquid nitrogen. Total proteins were extracted in SDS sample buffer and analyzed directly by SDS-page or further processed to isolate biotinylated proteins or for immunoprecipitation.

For the isolation of biotinylated proteins the protein extract was precipitated in four volumes of acetone, and washed twice with 400 µl acetone to remove free biotin. The proteins were then resuspended in binding buffer (100 mM Tris-HCl pH 7.5, 2% SDS, 8M urea) and incubated with 150 µl streptavidin magnetic beads (NEB) for 3 hrs with agitation. Beads were washed twice in binding buffer, twice with 1 M NaCl in 100 mM Tris-HCl pH 7.5, once with double distilled water, and once with ammonium bicarbonate pH 8.0. Success of the enrichment procedure was confirmed by analyzing 5% of beads by SDS-PAGE and immunoblotting. Enriched proteins were then identified by LC-MS/MS.

**Immunoblotting and protein detection**

Total leaf proteins were extracted in SDS sample buffer and separated by SDS-PAGE as described previously (Romand et al., 2022). Proteins were transferred onto a nitrocellulose membrane and probed with specific antibodies. Primary antibodies targeting the HA tag (monoclonal ab9110, Abcam) and GFP (polyclonal A-11122, Thermofisher) were used at a dilution of 1/5000. Biotinylated proteins were detected directly using streptavidin-horse radish peroxidase conjugate (RPN1231, Cytiva). Total proteins were visualized after separation and transfer using Ponceau Red protein stain. Original gel images are provided in the supplementary data files.

**Immunoprecipitation**

Total leaf proteins were extracted from 150 mg *N. benthamiana* powdered leaf disks in 1 ml Extraction Buffer (25mM Tris-HCl pH 7.5, 150 mM NaCl, 2 mM EDTA, 1% Triton X-100, 0.1% SDS, Complete protease inhibitor cocktail (Roche), phosphatase inhibitor cocktail 2 (Sigma), and phosphatase inhibitor cocktail 3 (Sigma)). 25 µl α-GFP nanobody:Halo:His6 coupled to Magne HaloTag beads (Promega) were added to each sample, which were incubated 2 hours at 4°C on a rotating wheel. After incubation, beads were magnetically separated, and then washed twice with Extraction Buffer and once with double distilled water. Proteins were then either analyzed by SDS-PAGE after eluting from the beads by heating with SDS sample buffer at 95°C for 10 min or analyzed by LC-MS/MS.

The α-GFP nanobody:Halo:His6 used for immunoprecipitation was produced using an α-GFP-nanobody:Halo:His6 construct (Addgene plasmid #111090) and prepared as described by Chen et al. (2018).

**Identification of peptides by LC-MS**

Proteins enriched on streptavidin or α-GFP nanobody:Halo:His6 beads were submitted to on-bead digestion. First, the beads were washed once with 160µL of 100 mM NH_4_HCO_3_,/CH3CN (v/v, 1/1), then reduced with 50 µL of 10 mM DTT in 100 mM NH_4_HCO_3_ for 45 min at 25 °C in the dark, and alkylated with 50 µL of 55 mM iodoactetamide in 100 mM NH_4_HCO_3_ for 30 min at 25 °C in the dark. The beads were washed once again as above, then resuspended in 50 µL of 25 mM NH_4_HCO_3_ with 0.025% ProteaseMAX (V/V) and 150 ng of Trypsin / LysC. The digestion was continued overnight at 37°C with agitation at 500 rpm. The free digested peptides were collected in a clean tube, the beads were washed twice with 25 µL of 0.1% in water (agitation 5 min, 500 rpm) and 25 µL of CH3CN (agitation 5 min, 500 rpm). All supernatants were pooled, dried down, resuspended in 0.05% TFA/ 2% acetonitrile in water, and quantified by the quantitative colorimetric peptide assay (Thermo Scientific). Peptides (600 ng) were injected on a reversed phase C18 column (Acclaim PepMap RSLC, 75μm x 150 mm, 2μm) and separated on a two step -linear gradient from 6% to 40% in 52 min of mobile phase B (80% acetonitrile/ 0.1% formic acid (FA) in water (v/v)) in mobile phase A (0.1% FA in water (v/v)), then from 40% to 65% of B in A for 11 min, followed by a 5-min chase at 99% of B. After ionization in the nanosource (source Easy Spray, ThermoFisher) at 1.9 kV spray voltage (capillary temperature set at 275 °C), the peptides were detected into the mass spectrometer Q-Exactive plus (Thermo Fisher) in positive ion mode. A top 10 data dependent acquisition mode was applied, alternating a scan event full MS in the Orbitrap analyzer at 70 000 resolution in a 350-1900 m/z range, and scan events of fragmentation (MS/MS) of the 10 top ion parents, in the Higher Energy Collisional Dissociation Cell, at 17 500 resolution, with a dynamic exclusion of 30 s.

Spectra were processed in Proteome Discoverer (Thermo Fisher Scientific, version: 2.4.1.15) using the algorithm Sequest HT and a peptide validator node based on Peptide Spectral Match level with maximum Delta Cn 0.05. LC-MS data were searched against an *N. benthamiana* protein database (Kourelis et al., 2019), target protein sequences (TID-LST8, LST8-TID, TID-YFP, RSH3-GFP, RSH3_39-221_-GFP), and a list of common contaminants. The protease was set as trypsin with up to 3 missed cleavages possible. Static modifications included carbamidomethylation (+57.021 Da), and dynamic modifications including Acetyl of the N-terminus (+42.011 Da), methionine oxidation (+15.995 Da), Met-loss (-131.040 Da), Met-loss+Acetyl (-89.030 Da), lysine biotinylation (+226.78 Da) in the sample containing TID, and serine or threonine phosphorylation (+79.966 Da). A maximum of 4 dynamic modifications per peptide was allowed. Phosphorylation sites on peptides were considered only at rank 1. Peptides identifying proteins were validated with the best PSM score calculated on PSM level FDR (0.01< Target FDR <0.05). Phosphopeptide identification and summaries are available in the supplementary data files, and the raw LC-MS data files will be soon available in ProteomeXchange Consortium (http://proteomecentral.proteomexchange.org) via the PRIDE partner repository with a specific dataset identifier PXD (Perez-Riverol et al., 2022).

**Detection of phosphoforms by Phos-tag**

Total leaf proteins were extracted in SDS sample buffer as described previously (Romand et al., 2022). EDTA present in the sample buffer was quenched with 10 mM MnCl_2_ before protein denaturation and loading in the gel. Phos-TAG gels were obtained by adding 25 µM Phos-TAG (Origin) and 50 µM MnCl_2_ to the 9% Acrylamide/Bis-Acrylamide SDS-PAGE preparation. Protein resolution was carried at 20 mA per gel, and after migration MnCl_2_ present in Phos-TAG gels was quenched by a 30-minute wash in 10 mM EDTA. Proteins were transferred onto a nitrocellulose membrane and probed with primary antibodies against GFP. Phos-TAG gels were compared with standard SDS-PAGE to identify phosphorylated proteins.

**TOR inhibition treatments**

We used three different TOR inhibition protocols depending on the type and growth stage of the plant. A stock solution of 10 mM AZD-8055 (Tocris) in DMSO was used for preparing buffers and media for TOR inhibition. For the 0 µM AZD-8055 control an equivalent quantity of DMSO was used.

For the inhibition of TOR in *N. benthamiana*, 6 mm leaf disks were removed from agroinfiltrated leaves and vacuum infiltrated with liquid half strength MS solution containing 10 µM AZD-8055. Then, leaf disks were floated on 10 ml of the respective treatment solution in round culture dishes under standard growth conditions for 2 hours. After incubation discs were frozen and total proteins extracted as described below.

For the inhibition of TOR in Arabidopsis seedlings, developmentally homogeneous 5-day old Arabidopsis seedlings of each genotype were transferred onto new square culture plates in a 6 by 3 grid pattern. Seedlings were treated by placing 45 1-µl drops of AZD-8055 stock solution (or the same volume of DMSO for the control) equally spaced between the seedlings for a final concentration of 10 µM AZD-8055 in the plate.

For the inhibition of TOR in mature Arabidopsis plants, entire rosettes were treated by spraying with liquid half strength MS solution containing 10 µM AZD-8055 or the equivalent quantity of DMSO. The rosette were sprayed twice at 24-hour intervals and plants were analyzed 48 hours after the first treatment.

**Nitrogen limitation treatment**

For nitrogen limitation plants were germinated on nitrogen replete half strength MS solution. After 5 days seedlings were transferred to square culture plates containing nitrogen replete (+N) (0.5× Murashige and Skoog salts [Caisson Labs], 1% sucrose, 0.5 g l−1 MES, and 0.4% Phytagel [Merck Sigma-Aldrich], adjusted to pH 5.7 with KOH) or nitrogen limiting (−N) (+N medium diluted 1/25 in 0.5× Murashige and Skoog medium without nitrogen [Caisson Labs], 0.5 g l−1 MES and 0.4% Phytagel [Merck Sigma-Aldrich], adjusted to pH 5.7 with KOH) growth medium. Plants were then returned to standard growth conditions.

**Microscopy**

Leaf discs were analyzed 3 days after agroinfiltration. The discs were mounted in perfluorodecalin (Merck Sigma-Aldrich) as described previously (Littlejohn and Love, 2012). Capture of fluorescence images was performed using the AxioImager APO Z1 microscope (Zeiss) using the following filters: chlorophyll, excitation 625–655 nm, emission 665–715 nm; mCHERRY, excitation 533–558 nm, emission 570–640 nm; GFP/GFP, excitation 455–495 nm, emission 505–555 nm; YFP, excitation 455–495 nm, emission 515–555 nm and CFP, excitation 431–441 nm, emission 460–500 nm. Standard exposure times of 10 ms for chlorophyll and 50-200 ms for fluorescent proteins was kept for all observations. No fluorescence bleed-through was observed between the different fluorescent protein channels. Images were captured from different regions of each inoculated leaf, and from at least two leaves per experiment. 10 µm deep Z stacks composed of 21 slices were acquired and then converted into maximum intensity projections in ZEN (Zeiss). Post-acquisition image processing was then performed Image J (Schneider et al., 2012; Schindelin et al., 2012). Unprocessed images were used for the quantification of normalized fluorescence intensities. The integrated signal density was calculated for the EGFP channel. Fluorescence localized in nuclei was divided by the average fluorescence present in the nuclei-attached chloroplast.

**Chlorophyll fluorescence measurements**

Plants were dark adapted for 20 min and chlorophyll fluorescence was measured in a Fluorcam FC 800-O imaging fluorometer (Photon System Instruments). PSII maximum quantum yield (Fv/Fm) was calculated as (Fm − Fo)/Fm. All experiments on photosynthetic parameters were repeated independently two to five times with similar results.

**Nucleotide quantification**

Nucleotides were extracted from about 150 mg of plant material, and quantified by HPLC–MS/MS using stable isotope labeled ppGpp and GTP standards as described previously (Bartoli et al., 2020).

**Phylogenetic inference**

Multiple-sequence alignments of homologous proteins were performed using MAFFT v7.40262 with option –auto (Katoh et al., 2019). Alignments were then trimmed to include only the N-terminal region up to the start of the (p)ppGpp hydrolase domain. Phylogenetic reconstructions were created using maximum likelihood with the IQ-TREE web server version 1.6.1164 using default settings, with LG + F + R7 automatically selected as the best fit evolutionary model based on BIC values by ModelFinder (Trifinopoulos et al., 2016). Branch support was tested using two methods: ultrafast bootstrap approximation using 1000 bootstraps, and the non-parametric Shimodaira–Hasegawa–like approximate likelihood-ratio test (aLRT). The alignments and trees are available in the supplementary data files.

**Creation of *rsh_2,3_* complementation lines**

RSH3_65-715_ was amplified by PCR and assembled with the RSH3_1-64_ region in a BsaI MoClo reaction to make a 35S:RSH3_1-715_-GFP-35S terminator Level 1 module. RSH3_65-715_ was also assembled with the CTP of the Rubisco small subunit 1A NT2 module in a BsaI MoClo reaction to make a 35S:SSU-RSH3_65-715_-GFP-35S terminator Level 1 module. The Level 1 modules were assembled with the OLE1:RFP reporter module(Engler et al., 2014; Shimada et al., 2010) in a BpiI MoClo reaction to make Level 2 multigenic constructs. The resulting constructs were transferred into *Agrobacterium tumefaciens* (strain GV3101) and used to transform *rsh_2,3_* plants by floral dipping. The majority of the recovered lines were silenced or showed very low expression. The following non-silenced lines were obtained: two *rsh_2,3_* OX:RSH3-GFP lines, four *rsh_2,3_* OX:SSU-RSH3-GFP lines, and one *rsh_2,3_* P- OXRSH3-GFP line.

**Data analysis**

Graphs and statistical tests were generated in Python (Python Software Foundation, <https://www.python.org/>) using the Panda (Reback et al., 2022), Matplotlib (Hunter, 2007) and Seaborn (Waskom, 2021) libraries. Statistical tests were performed using the Pingouin (Vallat, 2018) library. A Games-Howell Post-hoc test was adopted for non-parametric data comparisons and a pairwise T-test using the Benjamini/Hochberg FDR correction for multiple comparisons of data with normal distributions. Scripts and source data are available in the supplementary data files.

For tor/tor-1 (Menand et al., 2002)

**References**

**Bartoli, J., Citerne, S., Mouille, G., Bouveret, E., and Field, B.** (2020). Quantification of guanosine triphosphate and tetraphosphate in plants and algae using stable isotope-labelled internal standards. Talanta **219**: 121261.

**Chen, C., Masi, R.D., Lintermann, R., and Wirthmueller, L.** (2018). Nuclear Import of Arabidopsis Poly(ADP-Ribose) Polymerase 2 Is Mediated by Importin-α and a Nuclear Localization Sequence Located Between the Predicted SAP Domains. Frontiers in Plant Science **9**.

**Engler, C., Youles, M., Gruetzner, R., Ehnert, T.-M., Werner, S., Jones, J.D.G., Patron, N.J., and Marillonnet, S.** (2014). A golden gate modular cloning toolbox for plants. ACS Synth Biol **3**: 839–843.

**Gantner, J., Ordon, J., Ilse, T., Kretschmer, C., Gruetzner, R., Löfke, C., Dagdas, Y., Bürstenbinder, K., Marillonnet, S., and Stuttmann, J.** (2018). Peripheral infrastructure vectors and an extended set of plant parts for the Modular Cloning system. PLoS One **13**: e0197185.

**Hunter, J.D.** (2007). Matplotlib: A 2D Graphics Environment. Computing in Science Engineering **9**: 90–95.

**Katoh, K., Rozewicki, J., and Yamada, K.D.** (2019). MAFFT online service: multiple sequence alignment, interactive sequence choice and visualization. Brief Bioinform **20**: 1160–1166.

**Kourelis, J., Kaschani, F., Grosse-Holz, F.M., Homma, F., Kaiser, M., and van der Hoorn, R.A.L.** (2019). A homology-guided, genome-based proteome for improved proteomics in the alloploid Nicotiana benthamiana. BMC Genomics **20**: 722.

**Littlejohn, G.R. and Love, J.** (2012). A Simple Method for Imaging Arabidopsis Leaves Using Perfluorodecalin as an Infiltrative Imaging Medium. J Vis Exp: 3394.

**Menand, B., Desnos, T., Nussaume, L., Berger, F., Bouchez, D., Meyer, C., and Robaglia, C.** (2002). Expression and disruption of the Arabidopsis TOR (target of rapamycin) gene. Proceedings of the National Academy of Sciences **99**: 6422–6427.

**Patron, N.J. et al.** (2015). Standards for plant synthetic biology: a common syntax for exchange of DNA parts. New Phytol **208**: 13–19.

**Perez-Riverol, Y. et al.** (2022). The PRIDE database resources in 2022: a hub for mass spectrometry-based proteomics evidences. Nucleic Acids Res **50**: D543–D552.

**Reback, J. et al.** (2022). pandas-dev/pandas: Pandas 1.4.3.

**Romand, S. et al.** (2022). A guanosine tetraphosphate (ppGpp) mediated brake on photosynthesis is required for acclimation to nitrogen limitation in Arabidopsis. Elife **11**: e75041.

**Schindelin, J. et al.** (2012). Fiji - an Open Source platform for biological image analysis. Nat Methods **9**: 10.1038/nmeth.2019.

**Schneider, C.A., Rasband, W.S., and Eliceiri, K.W.** (2012). NIH Image to ImageJ: 25 years of Image Analysis. Nat Methods **9**: 671–675.

**Shimada, T.L., Shimada, T., and Hara-Nishimura, I.** (2010). A rapid and non-destructive screenable marker, FAST, for identifying transformed seeds of Arabidopsis thaliana. Plant J **61**: 519–528.

**Trifinopoulos, J., Nguyen, L.-T., von Haeseler, A., and Minh, B.Q.** (2016). W-IQ-TREE: a fast online phylogenetic tool for maximum likelihood analysis. Nucleic Acids Research **44**: W232–W235.

**Vallat, R.** (2018). Pingouin: statistics in Python. Journal of Open Source Software **3**: 1026.

**Velay, F., Soula, M., Mehrez, M., Belbachir, C., D’Alessandro, S., Laloi, C., Crete, P., and Field, B.** (2022). MoBiFC: development of a modular bimolecular fluorescence complementation toolkit for the analysis of chloroplast protein–protein interactions. Plant Methods **18**: 69.

**Waskom, M.L.** (2021). seaborn: statistical data visualization. Journal of Open Source Software **6**: 3021.

**Weber, E., Engler, C., Gruetzner, R., Werner, S., and Marillonnet, S.** (2011). A Modular Cloning System for Standardized Assembly of Multigene Constructs. PLoS One **6**: e16765.

**Xi, L., Zhang, Z., and Schulze, W.X.** (2021). PhosPhAt 4.0: An Updated ArabidopsisArabidopsis Database for Searching Phosphorylation Sites and Kinase-Target Interactions. In Plant Phosphoproteomics: Methods and Protocols, X.N. Wu, ed, Methods in Molecular Biology. (Springer US: New York, NY), pp. 189–202.
